# Uncovering the Role of the *scs* Pilus Within the Complex Surface Architecture of *Stenotrophomonas maltophilia*

**DOI:** 10.64898/2026.03.13.711640

**Authors:** Radhika Bhaumik, Gregory G. Anderson, Seema Mattoo

## Abstract

*Stenotrophomonas maltophilia* is an emerging multidrug-resistant pathogen that encodes numerous surface structures mediating attachment, biofilm formation, motility, and virulence. How individual adhesive systems function within this complex and potentially redundant network remains unclear. Here, we investigated the *scs* locus, a conserved chaperone-usher pilus system, to define its contribution to surface-associated behaviors and infection. Deletion of *scs* alone produced minimal effects on biofilm formation under standard laboratory conditions. However, significant and reproducible phenotypic differences emerged when *scs* loss was combined with mutations in other pilus systems or when bacteria were evaluated in infection-relevant environments. Across assays of biofilm formation, surface piliation, motility, flagellar gene expression, and virulence, *scs* contributed differentially to enhancing the adhesion and motility defects associated with the *smf-1* pili, *cbl* pili, and *fliC* flagellar gene loci. Transmission electron microscopy corroborated these findings, revealing differential expression and distinct alterations in predominantly the pilus architecture among mutant strains. These pilus associated phenotypes also largely corroborated infection in a *Galleria melonella* model. Given the abundance of adhesion associated gene loci in the pan-*S. maltophilia* genome, our results demonstrate that the *scs* locus functions as a context-dependent determinant of attachment and virulence, revealing a potential crosstalk between components of the surface landscape of the bacteria.

**Importance:** *Stenotrophomonas maltophilia* is an emerging multidrug-resistant pathogen that relies on diverse surface structures, including chaperone-usher pili, to drive attachment, biofilm formation, and infection. Its rising clinical prevalence underscores the need to define how multiple chaperone-usher pili contribute to virulence and interact with other appendages such as flagella, which together shape pathogenic behaviors. Our work on the *scs* chaperone-usher pilus system advances this understanding by revealing its context-dependent role in virulence. We demonstrate how *scs* modulates defects in *smf-1* and *cbl* pilin loci, and affects the *fliC* flagellar locus to reshape the cell surface pilus landscape, thereby illuminating how coordinated surface-structure interactions drive pathogenicity.

## Introduction

*Stenotrophomonas maltophilia* is an emerging multidrug-resistant pathogen capable of causing a wide range of human infections ^1,2^. Most notably, it is a common infectious agent in the lungs of individuals with cystic fibrosis (CF), with prevalence rates ranging from 8.7% to 16% reported by countries in Europe and North America^3,4,5^. *S. maltophilia* also initiates many other types of infections, especially in immunocompromised individuals, such as those with prior antibiotic use, patients on ventilators or with catheters, individuals with HIV, or those with malignancies or trauma ^2^ ^6^ ^7^ ^8^ ^9,10^. In many such cases, *S. maltophilia* isolates display extremely high levels of multidrug resistance^1^. One of the most common pathogenic mechanisms used by this bacterium is biofilm formation, which enables persistent survival on biotic and abiotic surfaces and contributes to chronic human infections, such as those occurring in the CF lung^11,12^.

Prior studies have revealed various factors involved in *S. maltophilia* biofilm formation. For example, *rmlA*, *rmlC*, *xanB*, and *spgM* contribute to forming the lipopolysaccharide layer of *S. maltophilia* which in turn affects biofilm formation ^13^ ^14^ ^15^ ^16^ ^17,18^. Additionally, a quorum sensing molecule called diffusible signaling factor (DSF) influences biofilm formation, presumably by regulating biofilm associated genes^19^. Environmental iron also plays a key role in biofilm development, signaling gene expression changes through the *fur* locus^20^. Our previous studies have shown that the glycolytic enzyme phosphoglycerate mutase, encoded by gene *gpmA*, plays an important role in *S. maltophilia* biofilm formation, through attachment or by mediating growth states in the presence of changing nutrient environments ^21,22^. Mutations in genes for each of the above factors result in a decrease in biofilm levels, although how these factors cooperate in biofilm development remains unclear.

The main mechanism by which *S. maltophilia* adheres, colonizes, and establishes biofilm is through its pili ^14^ ^23^ ^15,24^. So far, the main pilus recognized in *S. maltophilia* is SMF-1, a chaperone/usher pilus (CUP) associated with attachment to a variety of surfaces, agglutination of red blood cells, and biofilm formation ^23,25^. Sequencing of amplicons from CF-derived *S. maltophilia* strains indicate that the *smf-1* pilin gene, at locus *smlt0706* in type strain K279a, is highly prevalent among clinical isolates^26^. The SMF-1 pilus is also widespread among environmental strains of *S. maltophilia* ^27^ ^28,29^.

We have previously shown that the *smf-1* pilus gene is part of a putative CUP operon (smlt0706–0709) in the *S. maltophilia* type strain K279a. Building on this finding, we sought to investigate whether additional pili are present in this pathogen and assess their contribution to virulence. Through bioinformatic analyses we found two additional CUP operons in *S. maltophilia* type strain K279a which we name CBL (located at *smlt3830-3833*), as mentioned in our previous study^30^, and SCS (located in *smlt1508-1513*) (described in this study). These pilin genes are nearly universally present in clinical and environmental isolates as well. Additionally, we identified many other CUP operons spread throughout the *S. maltophilia* pan genome. Finally, we found an intriguing connection between the presence of *smf-1*, *cblA*, and *scs* pilus genes and flagellar expression and motility, which corresponded with infection. These data highlight potential crosstalk between virulence factors that dictate pathogenic mechanisms in *S. maltophilia*.

## 1. Materials and Methods

### 1.1 Identification of pilus operons

Pilus operons were initially identified in *S. maltophilia* type strain K279a, by searching the annotated genome on the NCBI database using search terms “pili”, “pilin”, “pilus”, “fimbriae”, “fimbrial”, “chaperone/usher”, “chaperone”, and “usher”. We determined whether these operons were present in other *S. maltophilia* strains by using the BLASTn function on the NCBI website to search for homologies to K279a putative pilus nucleotide sequences in all fully sequenced *S. maltophilia* genomes present in the NCBI database (as of November 10th, 2025). Subsequently, we used the same search terms to identify additional, novel pilus operons and genes from all fully sequenced *S. maltophilia* genomes in the NCBI database.

### 1.2 Bacterial strains and culture conditions

For our experiments, we used *S. maltophilia* clinical strain K279a, a fully sequenced type strain^31^. *Escherichia coli* strain S17-1 was used for the creation and maintenance of the genetic constructs^32^. Bacterial strains were cultured in LB medium, with gentamicin supplementation as needed to maintain plasmids (10 μg/mL for plasmids in *E. coli* and 70 μg/mL for plasmids in *S. maltophilia*). Isogenic deletions were created in *smf-1* and *cblA* genes in *S. maltophilia* K279a using an allelic replacement technique, as previously described ^21^ ^22,30^. We also generated the pΔ*scs* (with primer sets 1513LFor/1513LRev and 1513RFor/1513RRev) deletion plasmid (**Table 1**). This construct was used to create a Δ*scs* strain following the same strategy outlined in our previous study ^30^ which we confirmed with 1513For/1513Rev primer sets (**Table 1**). Using the pΔ*smf-1*, pΔ*cblA*, and pΔ*scs* deletion plasmid constructs, we similarly generated the double mutant strains Δ*smf-1* Δ*scs* and Δ*cblA* Δ*scs* ^21,22^, in a manner similar to the construction of previously described Δ*smf-1* Δ*cblA*^30^. Bacteria were grown overnight in LB liquid medium with shaking at 37*°*C prior to inoculation in experiments.

**Table 1.**
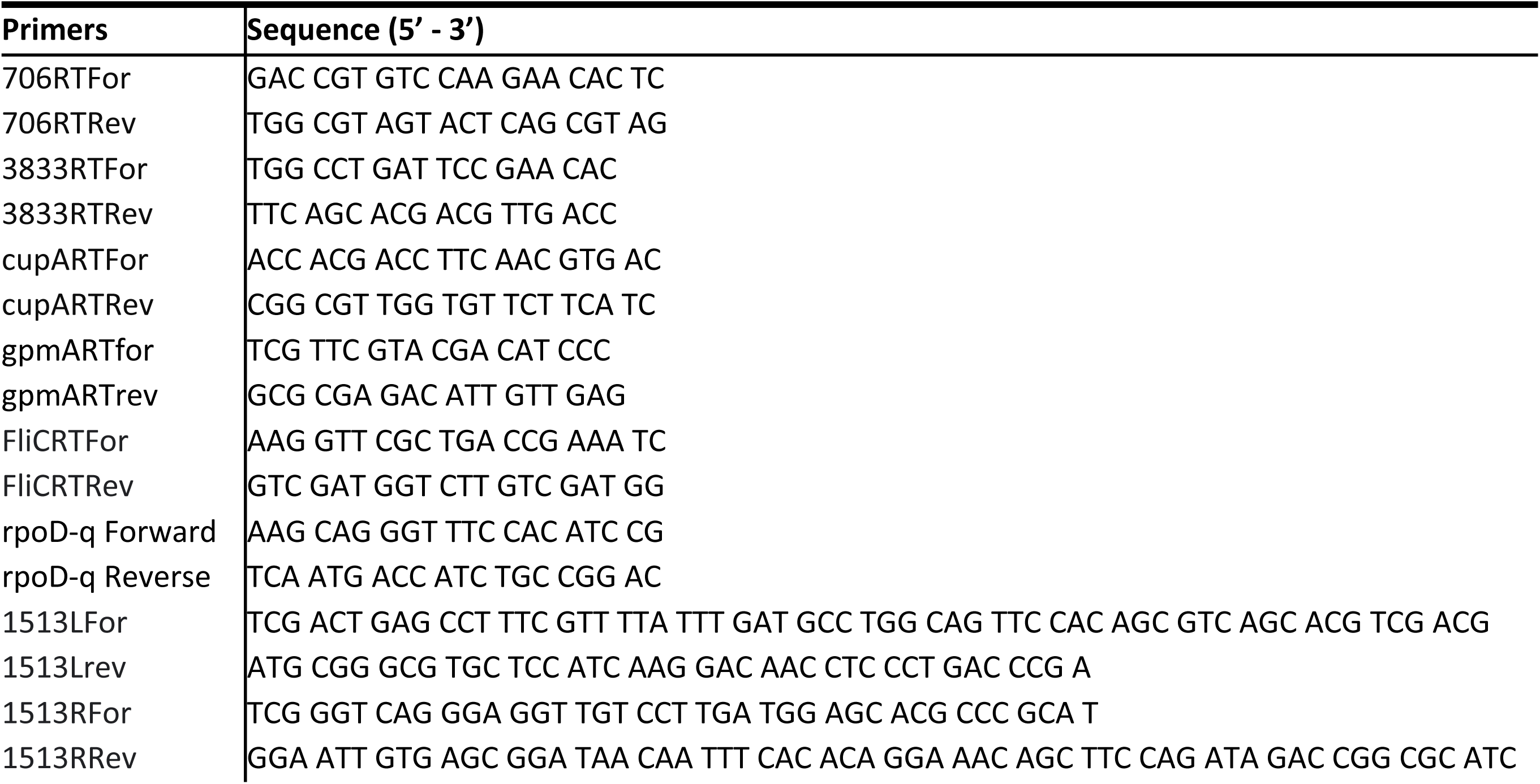
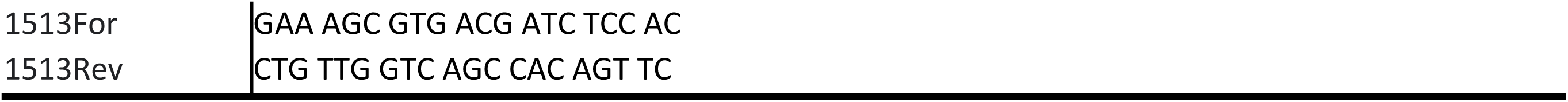
Primers used in this study.

### 1.3 Quantification of biofilm formation on polystyrene microtiter plate

For performing biofilm assays, overnight grown bacteria were diluted 1:100 into fresh LB medium ^33,34^. These diluted strains were then added at a volume of 100μL in individual wells of a 96-well polystyrene microtiter plate (Greiner Bio-One; Monroe, NC) and incubated statically for the indicated time point at 37°C. Each strain was added in quadruplicate in the plate. After the incubation, the plates were stained with 0.1% crystal violet (CV) for 15 min to detect bacterial cell attachment and biofilm formation, as described^34^. The wells were washed to remove any unattached stain and dried overnight. The stain was then solubilized using 30% acetic acid and 125μL solubilized stain were transferred to a new 96-well polystyrene microtiter plate. The intensity of the stain was then quantified with a SpectraMaxM2 spectrophotometer (Molecular Devices; Sunnyvale, CA) by measuring optical density at 550nm (OD₅₅₀). We performed a similar biofilm assay in Synthetic Cystic Fibrosis Sputum Medium (SCFM2) that mimics the chemical composition of CF lung environment^35^.

### 1.4 Transmission Electron Microscopy (TEM) analysis

Bacterial cultures were grown overnight in 5mL of LB medium at 37°C until reaching an OD₆₀₀ of 1. Cells were then washed twice with 1X DPBS and negatively stained with 2% phosphotungstic acid on carbon-Formvar nickel grids^23^. A 3μL sample (∼6000 bacterial cells) was applied to the grids and imaged using a Tecnai 12 transmission electron microscope (FEI, Hillsboro, OR) equipped with a LaB₆ crystal at 120kV. Pili were manually quantified from five independent micrographs per strain.

### 1.5 Galleria mellonella survival assay

*In vivo* virulence of *S. maltophilia* was assessed using a *Galleria mellonella* infection model as previously described^20^. Final instar *G. mellonella* larvae were obtained from BugCo (Ham Lake, MN). Bacterial cultures were grown overnight in 5 mL of LB medium at 37°C to an OD₆₀₀ of 1, followed by two washes in 1X PBS and resuspension in the same buffer. Larvae were injected with 10 μL of each bacterial strain at a concentration of 10⁵ CFU/10µL using a 20 µL Hamilton syringes, targeting the last prolegs. Each bacterial strain was tested in five larvae, per experiment. After infection, larvae were incubated in filter paper-lined Petri dishes at 37°C and monitored daily for survival over a 7-day period. Larvae were scored as dead based on lack of movement and increased melanization. A control group was injected with 10 µL of 1X PBS to account for any mortality due to mechanical injury; all PBS-injected larvae survived throughout the experimental time course. Survival curves were generated using the Kaplan-Meier method, and statistical significance was determined using the log-rank test (GraphPad Prism version 9.0, Software Inc., La Jolla, CA)

### 1.6 Alignment of pilus gene sequences

The *smf-1*, *cblA*, and *scs* gene sequences from all completed *S. maltophilia* genomes listed in the NCBI database were taken and aligned in Jalview.Ink which also provided percent identity to strain K279a.

### 1.7 PCR testing of pilus gene sequences

A bank of 51 clinical *S. maltophilia* isolates was obtained from Dr. Valerie Waters, at the Hospital for Sick Kids and St Michael’s Hospital, Toronto. To analyze these strains for the presence of *smf-1*, *cblA*, and *scs* genes, cultures of these isolates were grown overnight at 37°C in LB medium, with shaking, and their chromosomal DNA was isolated with the Puregene Yeast/Bact. Kit B (Qiagen) according to the manufacturer’s instructions. 1μL genomic DNA from *S. maltophilia* K279a was used as template for a PCR reaction using primer sets 706RTFor/706RTRev, 3833RTFor/3833RTRev, and cupARTFor/cupARTRev (**Table 1**), to test for *smf-1*, *cblA*, and *scs* genes, respectively. As a control transcript, the *gpmA* gene (*smlt1430*; phosphoglycerate mutase) ^21^was similarly tested, using primer pairs gpmARTfor/gpmARTrev. The PCR products were electrophoresed through a 1% agarose gel. *∼*800bp bands were considered a positive result for the presence of these pilus genes.

### 2.8 Motility assays

#### 2.8.1 Swimming motility

*S. maltophilia* swimming motility was assayed on tryptone broth with 0.3% (w/v) agar. 5μL of overnight grown bacterial culture were inoculated half-way into the plate via puncturing by the pipette tip carrying the bacteria. The plates were incubated at 37°C for 24 hours. The diameters of the swimming zones were measured along 3 different axes, and we recorded the overall diameter of an individual colony as the average of these 3 measurements.

#### 2.8.2 Swarming motility

*S. maltophilia* swarming motility was assayed on nutrient broth with 0.5% (w/v) agar. 2.5μL of overnight grown bacterial culture were dispensed onto the surface of the agar plates. The plates were incubated at 37°C for 24 hours. The diameters of the swimming zones were measured along 3 different axes, and we recorded the overall diameter of an individual colony as the average of 3 measurements.

#### 2.8.3 Twitching motility

*S. maltophilia* twitching motility was assayed on trypticase soy broth (TSB) with 1% (w/v) agar. 2.5μL of overnight grown bacterial culture were inoculated by puncturing the agar completely and dispensing the culture at the bottom of the agar at the Petriplate surface. The plates were incubated at 37°C for 72 hours. The gel was then loosened and scooped out with a spatula without disturbing the twitching zones. The exposed zones were stained with 0.1% crystal violet for 15 minutes and were then washed to remove any unattached stain. The diameters of the swimming zones were measured along 3 different axes, and we recorded the overall diameter of an individual twitching zone as the average of these 3 measurements.

### 2.9 Expression of *fliC* gene

The different strains of *S. maltophilia* were analyzed for their expression of flagellin gene *fliC* (s*mlt2304*), encoding the FliC monomer that comprises major structural component of the flagellar filament ^24,36^. *S. maltophilia* K279a wild type, Δ*smf-1*, Δ*cblA*, Δ*scs*, Δ*smf-1* Δ*cblA*, Δ*smf-1* Δ*scs*, Δ*cblA* Δ*scs,* and Δ*smf-1* Δ*cblA* Δ*scs* were grown in LB medium overnight at 37°C with shaking, and their RNA was isolated with the RNAeasy Kit (Qiagen) according to manufacturer’s instructions. The RNA was then converted to cDNA by SuperScript III First-Strand Synthesis System for RT-PCR (Invitrogen) according to manufacturer’s instructions. RNA was then removed by incubation with RNaseH for 20 min. *fliC* transcript level in each strain was assessed semi-quantitatively using 1 μL cDNA from each strain as template in a PCR reaction with primer pair FliCRTFor/FliCRTRev (**Table 1**)^33^. PCR was carried out in Bio-Rad T100 Thermal Cyclers as follows: 1 cycle of 95°C for 10 mins, followed by 25 cycles of denaturation at 95°C for 30 secs, annealing at 58°C for 30 secs, and extension at 72°C for 1 min. Finally, we incubated the tubes at 72°C for 10 mins and stored them at 4°C. As a control transcript, the *rpoD* gene (s*mlt4165*; RNA polymerase sigma-70 factor) ^37^was similarly tested, with primer pairs rpoD-q Forward/ rpoD-q Reverse (**Table 1**). PCR products were electrophoresed through a 1% agarose gel.

### 2.10 Growth kinetics

Bacterial strains were cultured overnight in LB at 37°C and then diluted 1:100 into fresh LB or SCFM2 medium. 100μL aliquots of each strain were transferred into quadruplicate wells of a 96-well polystyrene microtiter plate. The plates were incubated at 37°C in a SpectraMax M2 spectrophotometer, where absorbance at OD₆₀₀ was measured every 30 minutes for 20 hours, with 5 seconds of shaking prior to each reading.

### 2.11 Statistical analyses

Each experiment was performed at least three times with triplicate or quadruplicate samples for each strain. Student’s t-test was used to determine statistical significance. A difference was considered statistically significant at a p value of <0.05. As mentioned, a log-rank test was used to determine statistical significance for the *G. mellonella* infection survival, with p<0.05 considered significant.

## 2. Results

### 2.1 *scs* gene is important in biofilm formation

Our previous studies identified a role for SMF-1 and Cbl pili in *S. maltophilia* in biofilm formation^30^. Using a bioinformatic approach searching the NCBI database (see Materials and Methods), we identified another putative CUP gene cluster in strain K279a: *smlt1508-1513*. Gene proximity within these clusters suggests potential operonic organization (**Fig. 1**). While pending experimental evaluation, this operonic organization is supported by operon prediction software ((http://microbesonline.org/operons). The *smlt1513* locus exhibits homology to the Csu pilus of *Acinetobacter baumannii* ^38,39^, and we refer to this system as SCS (*<u>S</u>tenotrophomonas* <u>Cs</u>u pili) (**Fig. 1**). To assess the role of the putative pilin gene *scs* (*smlt1513*) in *S. maltophilia* biofilm formation, we generated an isogenic deletion mutant using allelic replacement to create the Δ*scs* strain. Biofilm assays performed in LB medium in 96-well polystyrene microtiter plates demonstrated that, in contrast to the Δ*smf-1* and Δ*cblA* strains, the Δ*scs* strain exhibited biofilm levels comparable to wild-type (WT), suggesting that the *scs* gene does not play a significant role in biofilm formation under the tested conditions (**Fig. 2A**). This lack of biofilm phenotype is likely why we did not identify this locus in our original transposon screen for biofilm deficient mutants.^21^. We also examined the role of *scs* gene in biofilm formation in SCFM2 medium. SCFM2 is a physiologically relevant setting as it closely replicates the nutrient-rich environment of CF lung sputum, containing experimentally measured concentrations of ions, free amino acids, glucose, lactate, mucin, lipids, proteins, DNA, and other components. At the early (4 hr) time point, the *Δscs* strain exhibited reduced biofilm formation compared to the wild type (WT); however, by 24 hr, biofilm formation in this mutant equaled or even surpassed WT levels (**Fig. 3A**).

**Figure 1.**
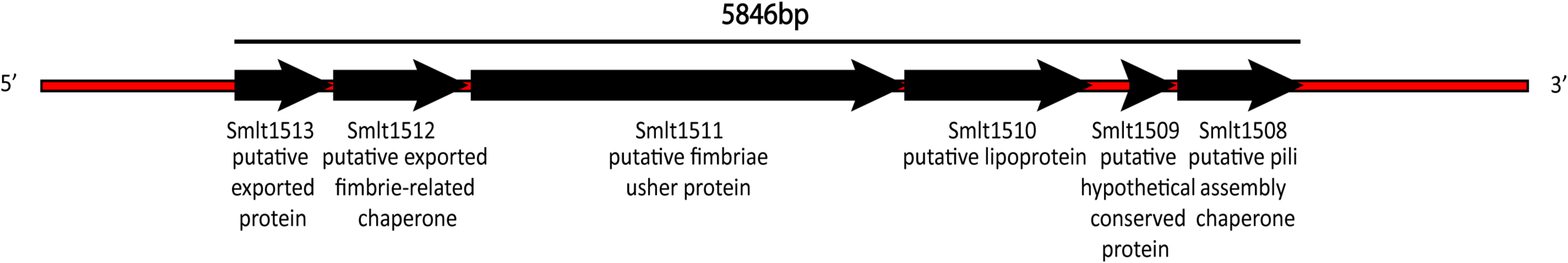
Gene organization of the putative *S. maltophilia scs* chaperone usher pilus operon. Gene numbers are listed within each box, and putative functions are listed below. Box sizes indicate relative length of each gene. The *scs* operon is transcribed on the complementary strand.

**Figure 2.**
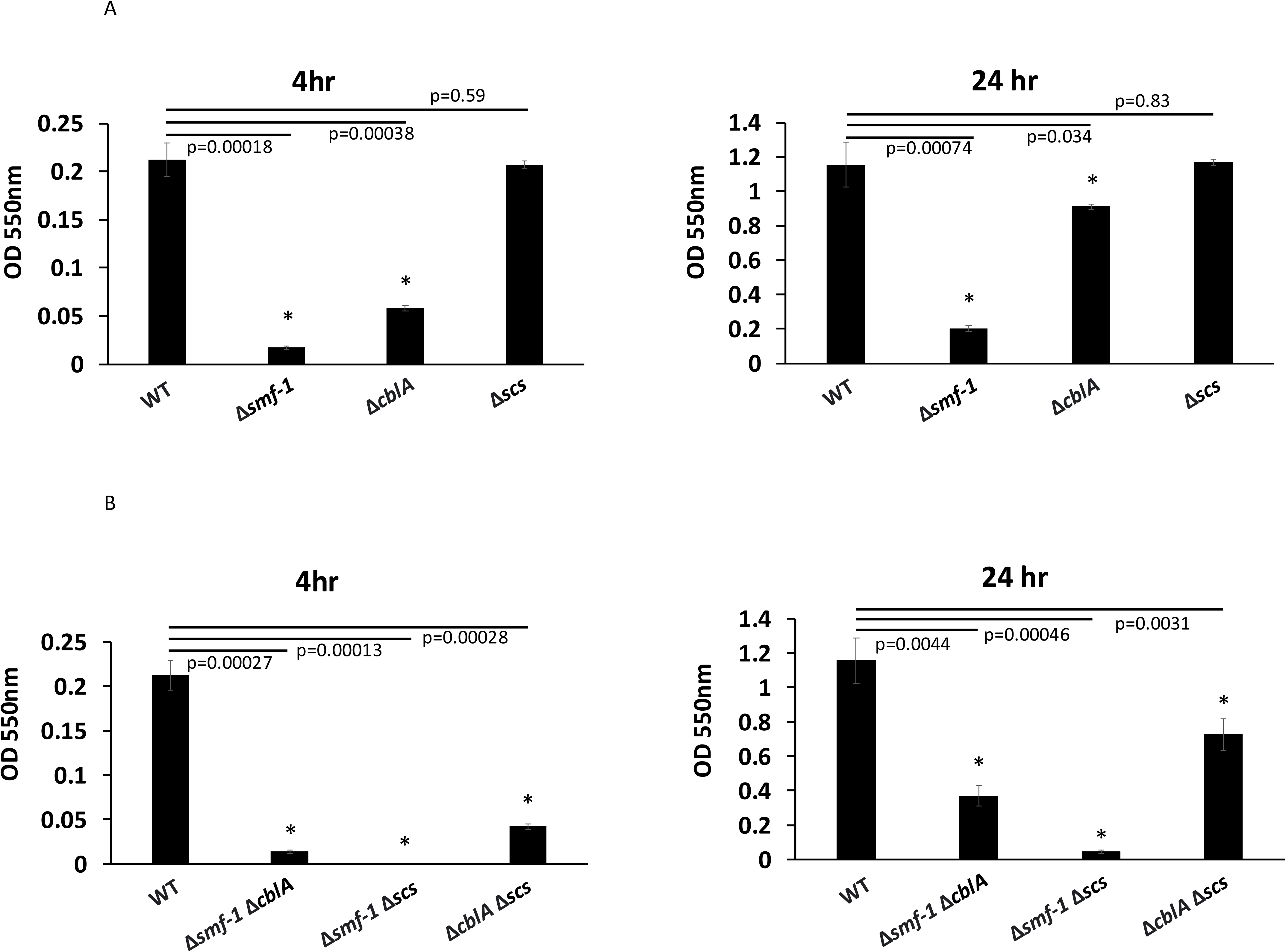
Isogenic deletion of pili genes dramatically reduces *S. maltophilia* biofilm formation in LB medium. Biofilms of *S. maltophilia* (A) wild type (WT), Δ*smf-1,* Δ*cblA,* Δ*scs* and (B) double mutants Δ*smf-1* Δ*cblA,* Δ*smf-1* Δ*scs*, and Δ*cblA* Δ*scs* in LB medium for 4 hr (left panels) or 24 hr (right panels). *p<0.05. These figures are the combination of three independent experiments (n=3) carried out in quadruplicate and error bars indicate standard deviations.

**Figure 3.**
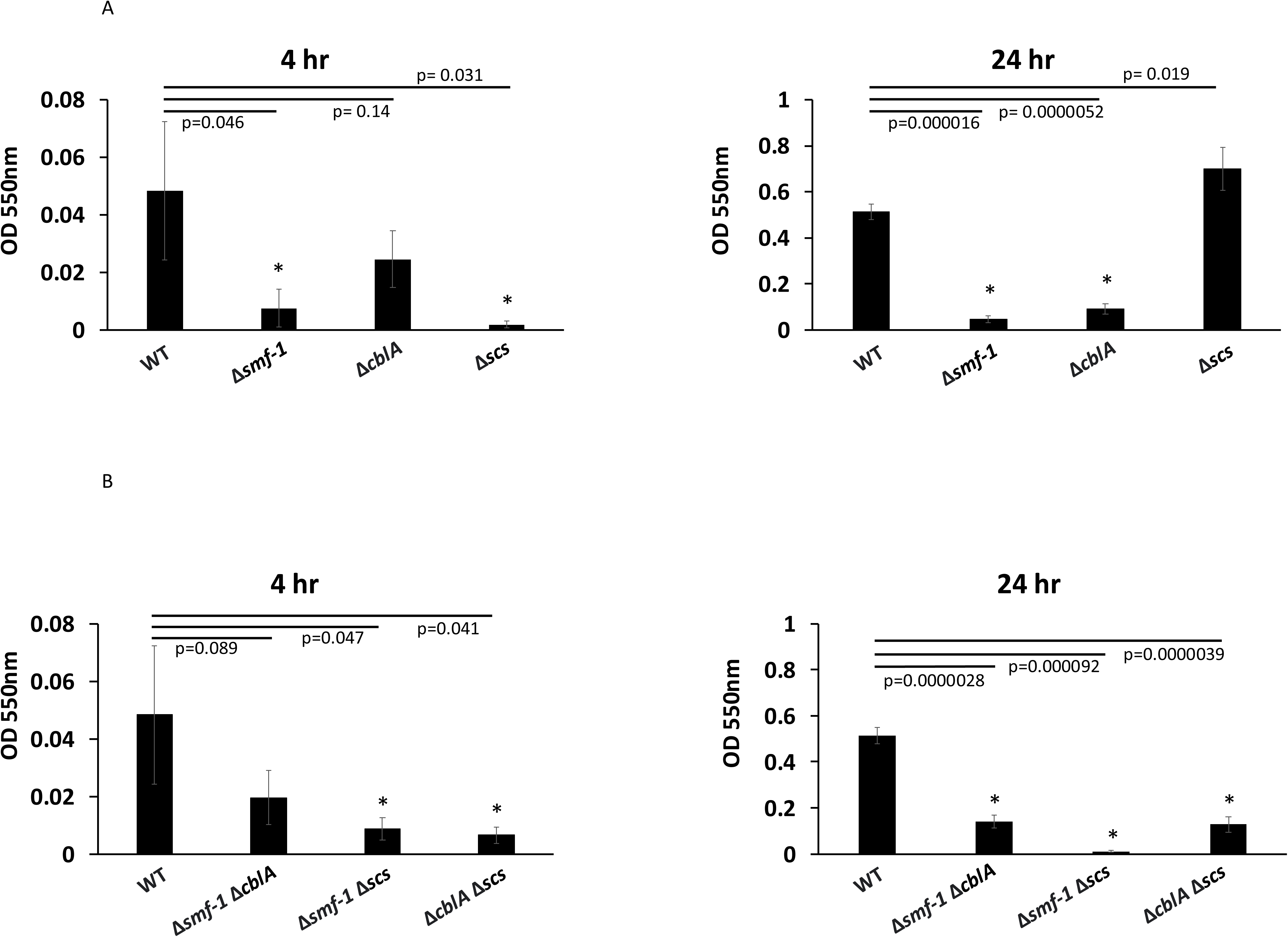
Isogenic deletion of pili genes dramatically reduces *S. maltophilia* biofilm formation in SCFM2 medium. Biofilms of *S. maltophilia* (A) wild type (WT), Δ*smf-1,* Δ*cblA,* Δ*scs* and (B) double mutants Δ*smf-1* Δ*cblA,* Δ*smf-1* Δ*scs*, and Δ*cblA* Δ*scs* in SCFM2 medium for 4 hr (left panels) or 24 hr (right panels). *p<0.05. These figures are the combination of three independent experiments (n=3) carried out in quadruplicate and error bars indicate standard deviations.

We next tested whether Scs pili showed phenotypes in combination with deletion of other pilus loci, as had been previously observed for a double mutant Δ*smf-1 ΔcblA* strain^30^. Therefore, we created the other two double mutant constructs: Δ*smf-1* Δ*scs* and Δ*cblA* Δ*scs*. We saw a significant decrease in the biofilm levels between each double mutant strain and the wild type at 4 hr and 24hr (**Fig. 2B**). At both time points, combination of Δ*scs* to Δ*smf-1* or Δ*cblA* showed an added reduction in biofilm levels compared to wild type or Δ*smf-1* or Δ*cblA* single mutants. Given the Δ*scs* single mutant does not demonstrate a biofilm defect, but only manifests a decreased biofilm when combined with Δ*smf-1* or Δ*cblA* suggests a plausible coordination of biofilm formation between these three pilus systems (**Fig. 2A**). Importantly, growth curve experiments demonstrated that all mutants grew at the same rate as wild type (**Supplementary Fig. 1**).

Interestingly, when biofilm was assessed in the more physiologically relevant SCFM2 media, we saw a significant decrease in biofilm formation with the Δ*scs* single mutant only at early (4 hr) time points compared to wild type (**Fig. 3A**). Accordingly, all double mutants also showed a defect in biofilm formation at both 4 hr and 24 hr time points when compared to wild type (**Fig. 3B**). All mutants showed similar growth kinetics to wild type in SCFM2 medium (**Supplementary Fig. 2**).

Taken together, these data emphasize a contributory role of Scs pili during biofilm formation.

### 2.2 TEM characterization of Δ*scs*

To examine pilus expression on the bacterial surface, we observed the wild type and mutant strains by TEM. The wild type qualitatively exhibited distinct pili structures, categorized into shorter chaperone/usher pili, consistent with previous reports on SMF-1^23^, and longer type IV pili ^40^ (**Fig. 4**), as previously described^30^. The Δ*scs* and Δ*cblA* Δ*scs* strains displayed similar pilus levels to the wild type. Manual quantification of these TEM images further supported these findings, revealing an average of approximately 90, 95, and 92 chaperone/usher pili for the wild type, *Δscs,* and Δ*cblA* Δ*scs* strains, respectively. In contrast, there was complete loss of the shorter chaperone/usher pili ^23^ on the surface of the Δ*smf-1* Δ*scs*, or Δ*smf-1* Δ*cblA* strains. While chaperone/usher were not detected in some of the Δ*cblA* strain cells, other cells exhibited low-level expression, with an average of approximately ∼5 pili observed (**Supplementary Fig. 3**). These observations are consistent with the reduced biofilm levels observed for the corresponding single and double mutant strains (**Fig. 2**), where strains exhibiting reduced biofilm showed little to no detection of chaperone/usher pili and were dominated by longer type IV pili. It should be noted that these assessments represent qualitative observations based on TEM imaging and previous literature^30^; definitive identification of chaperone/usher or type IV pili would require additional structural, immunological and biochemical experimental validation beyond the scope of our studies.

**Figure 4.**
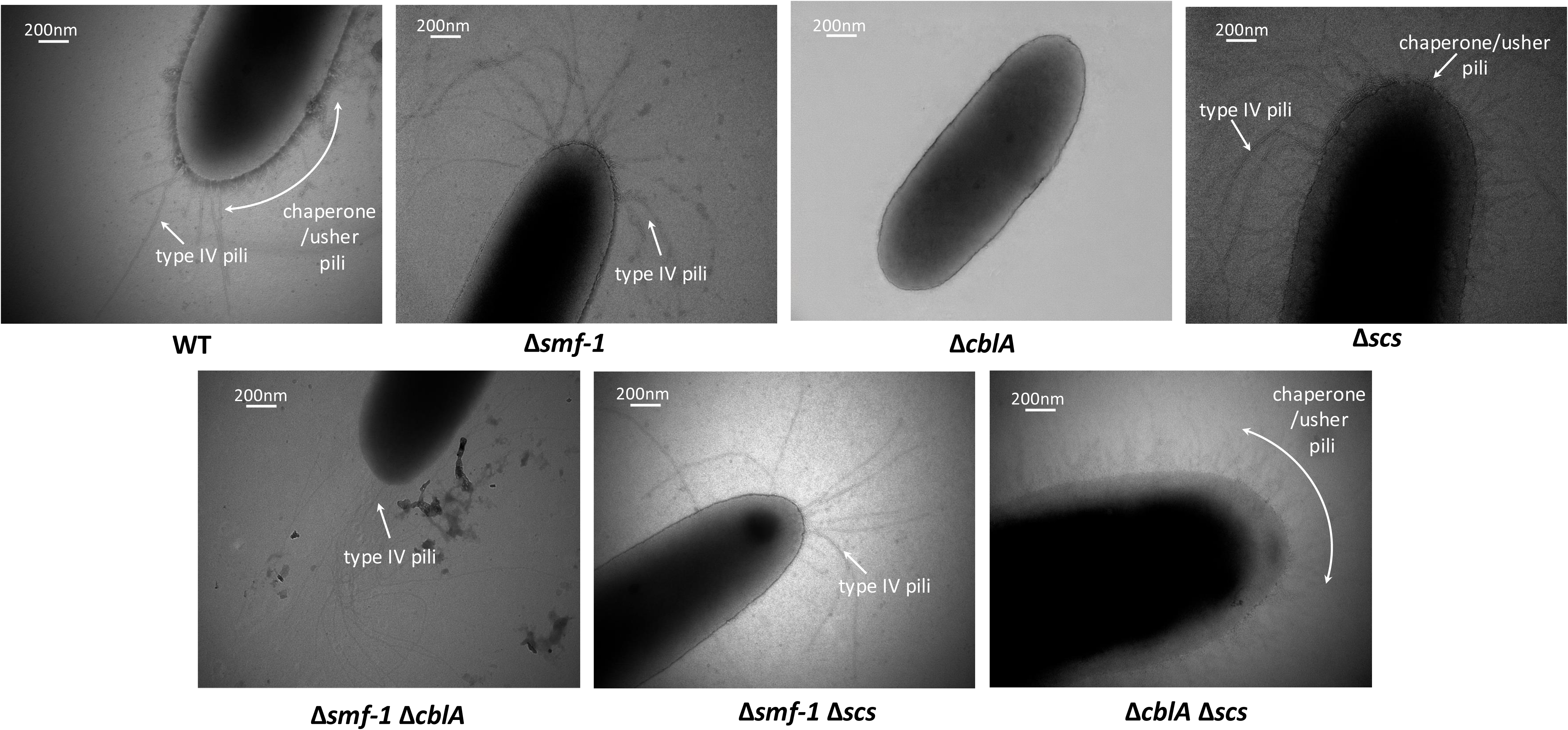
TEM studies show the presence of chaperone/usher pili (short and hairy) and type IV pili (long) in different bacterial strains. TEM images of WT, Δ*smf-1,* Δ*cblA,* Δ*scs*, double mutants Δ*smf-1* Δ*cblA,* Δ*smf-1* Δ*scs*, and Δ*cblA* Δ*scs* strains. Strains are indicated underneath each micrograph. Shorter pili correlate to chaperone/usher pili and longer structures correlate with type IV pili, as seen in previous studies. See text for details. Scale bars are as indicated on the micrographs.

Our TEM evaluations demonstrate that the loss of an individual pilus gene leads to distinct cell surface alterations that are not compensated by the remaining pilus systems.

### 2.3 Pilus mutants display variable motility patterns

Motility plays a major role in the establishment of biofilm formation^41^. First, bacterial cells perform flagellar-mediated swimming motility to find a favorable contact surface. Upon establishing attachment, the bacteria utilize flagella for swarming across the surface, while employing type IV pili to twitch, pulling neighboring bacteria closer to form clumps. This clustering initiates biofilm proliferation. We were curious whether our pilus mutations affected bacterial motilities. Thus, we tested *S. maltophilia* strains K279a wild type, Δ*smf-1*, Δ*cblA*, Δ*scs*, Δ*smf-1* Δ*cblA*, and Δ*smf-1* Δ*scs* in agar-based swimming, swarming, and twitching assays. While we observed variable motility levels in single mutants compared to wild type, these trends were in general not statistically significantly different. Only the Δ*smf-1* Δ*cblA* and Δ*smf-1* Δ*scs* mutants displayed any significant (p<0.05) decrease in twitching motility (**Fig. 5A**). We saw a similar trend for swimming and swarming motilities. While not statistically significant, the Δ*scs* showed the highest swimming and swarming motility amongst all strains compared to the wild type (**Fig. 5B**, **5C**), while Δ*smf-1*, Δ*smf-1* Δ*cblA*, and Δ*smf-1* Δ*scs* displayed decreased swimming and swarming compared to wild type – matching their respective biofilm phenotypes (**Fig. 2**).

**Figure 5.**
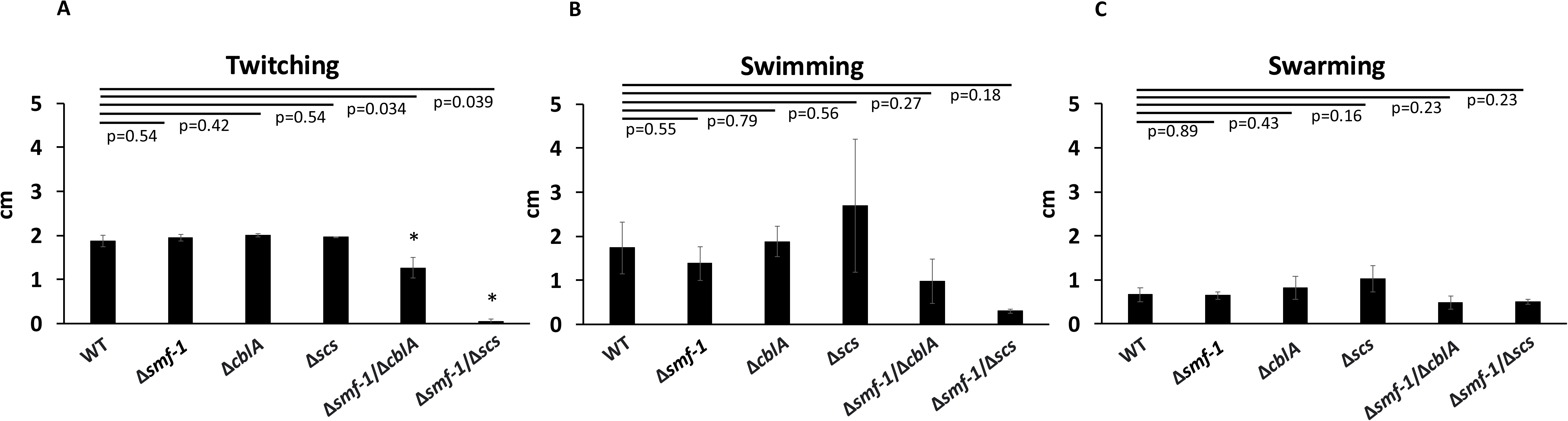
Motility assays of *S. maltophilia* K279a WT and mutant strains. WT, Δ*smf-1*, Δ*cblA*, Δ*scs*, Δ*smf-1* Δ*cblA*, and Δ*smf-1* Δ*scs* were assayed in (A) twitching (B) swimming and (C) swarming motility experiment. *p<0.05, compared to wild type. These figures are the combination of three independent experiments (n=3) carried out in quadruplicate and error bars indicate standard deviations.

### 2.4 The Δ*scs* strain exhibits high levels of *fliC* transcript

Considering the motility trends seen in **Fig. 4**, and their correlation with biofilm phenotypes (**Fig. 2**), we speculated that pilus mutation could affect flagellar gene expression. Studies have indicated that flagella play a vital role in *S. maltophilia* motility^41^. The flagella also aid *S. maltophilia* attachment in certain circumstances, such as colonization of mouse tracheal mucus^42^. The *S. maltophilia fliC* gene encodes flagellin, the major structural component of flagella^24^, and it is found in approximately 90% of *S. maltophilia* isolates^43^. Thus, we performed a semi-quantitative expression analysis of the *fliC* transcript in *S. maltophilia* wild type, Δ*smf-1*, Δ*cblA*, Δ*scs*, Δ*smf-1* Δ*cblA*, Δ*smf-1* Δ*scs*, and Δ*cblA* Δ*scs,* to determine whether pilus mutation affects flagellar gene expression level. We found that the Δ*scs* strain showed a much more intense band than wild type and other pilus mutant strains (**Fig. 6**), which matches with our previous result of higher swimming and swarming motility in this strain (**Fig. 4B**, **4C**) and the higher biofilm levels (**Fig. 2**), compared to wild type. Conversely, we found that the Δ*smf-1* Δ*scs,* double mutant exhibited the faintest band, coinciding with its most defective motility and severely impaired biofilm formation among all the strains tested. We also saw a low intensity band for Δ*smf-1* and medium intensity bands for wild type and Δ*cblA* Δ*scs*. Taken together, these results align well with the motility assays and biofilm levels described above.

**Figure 6.**
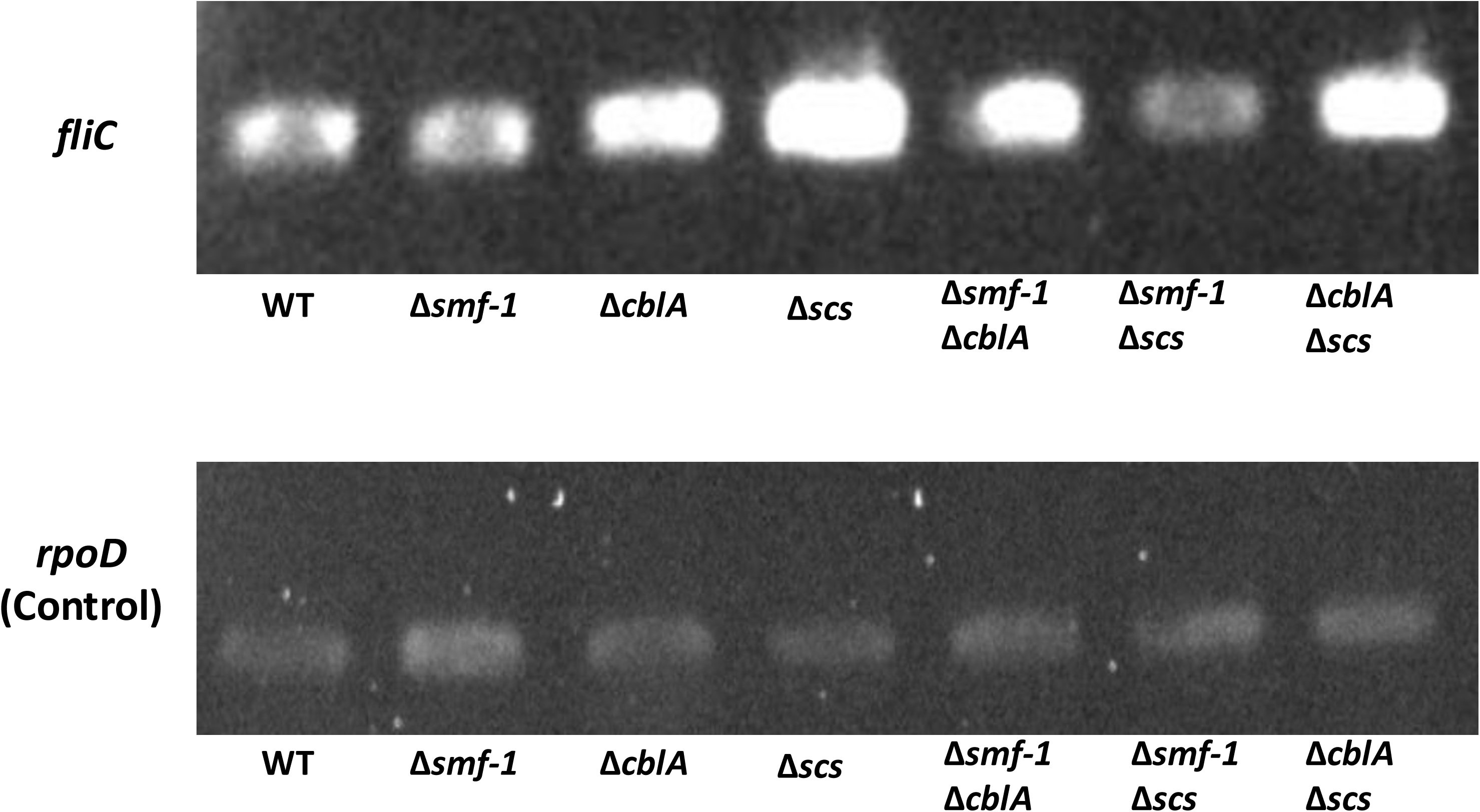
Semiquantitative expression analysis of *fliC* in pilus mutant strains. (Top) PCR analysis of flagellin gene sequence *fliC* for *S. maltophilia* wild type (WT) and mutant strains. The Δ*scs* strain shows the highest intensity and Δ*smf-1* Δ*scs* strain shows the lowest intensity of *fliC* transcript. (Bottom) *rpoD* is an expression control transcript. We do not see any difference in *rpoD* band intensity for any of the strains.

### 2.5 Pilus mutants show increased virulence in a *Galleria mellonella* infection model

*Galleria mellonella* has been widely used as an infection model to study host-pathogen interactions, including those of *S. maltophilia* ^20,44^. We infected *G. mellonella* larvae with wild type and pilus mutant strains, as described in Materials and Methods. In contrast to the larvae infected with wild-type *S. maltophilia* that exhibited 0% mortality (**Fig. 7**) we saw increased virulence for the Δ*smf-1* Δ*scs* double mutant strain, as well as for the Δ*smf-1* Δ*cblA* strain (80% mortality after 1 day). The ∼60% mortality observed in larvae infected with the Δ*smf-1* strain indicates that, despite intact *scs* and *cblA* loci, the presence of these pili alone is insufficient to mitigate the increased virulence associated with *smf-1* deletion. Furthermore, infection with the Δ*scs* and Δ*cblA* Δ*scs* strains showed a trend toward decreased survival compared to wild type; however these differences were not statistically significant. Importantly, all larvae in the control group (injected with 1X PBS) survived throughout the experiment, confirming that mechanical injury did not contribute to mortality. These data emphasize the contribution of *scs* pili during infection and virulence associated phenotypes.

**Figure 7.**
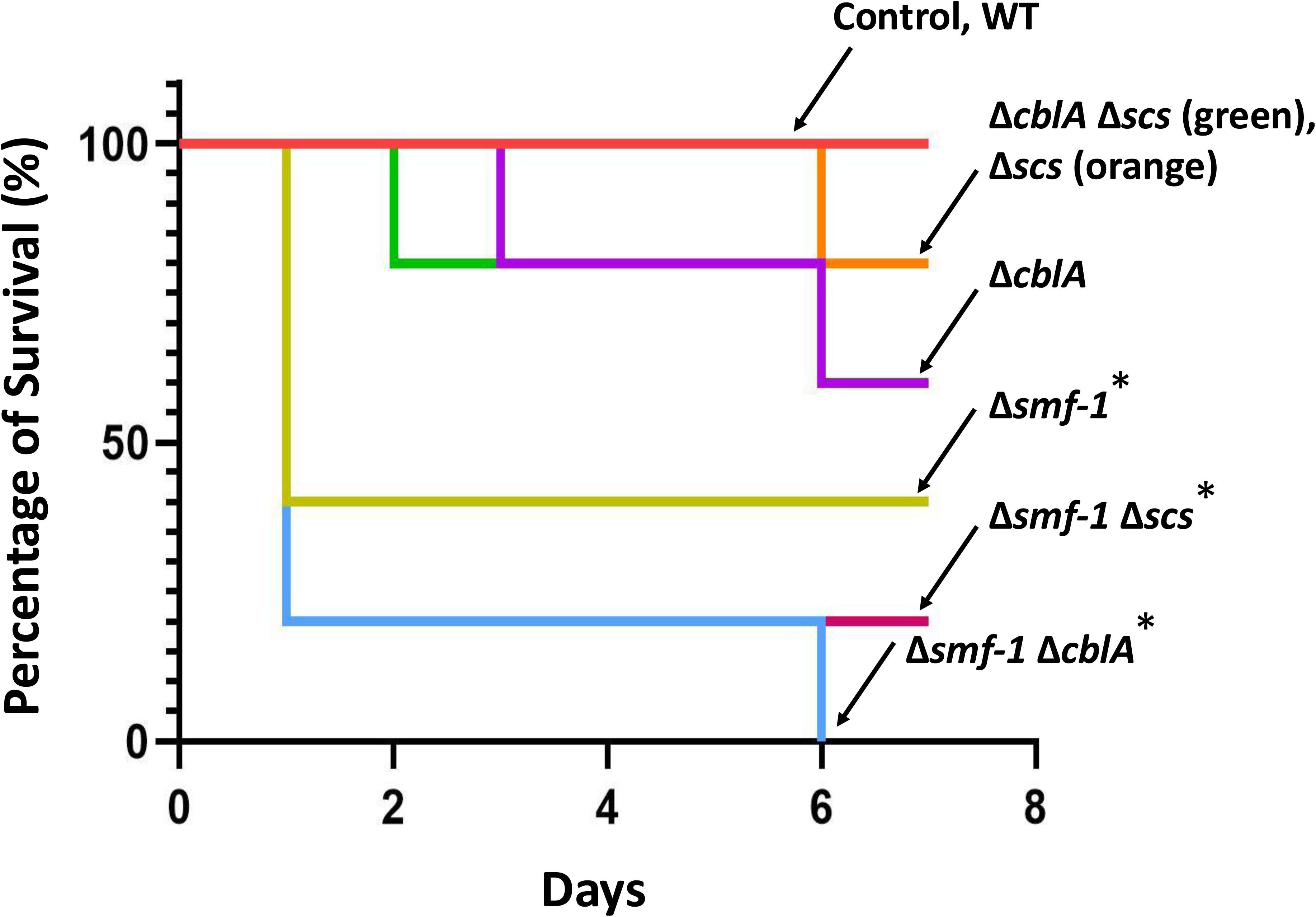
Pilus mutants are highly virulent in the *G. mellonella* infection assay. *G. mellonella* larvae were infected with wild type (WT), Δ*smf-1*, Δ*cblA*, Δ*scs*, Δ*smf-1/*Δ*cblA*, Δ*smf-1*/Δ*scs*, and Δ*cblA*/Δ*scs* strains and survival was monitored over 7 days. The survival graph is representative of three independent experiments (n=3) carried out in quintuplicate. *p<0.05 compared to the WT.

### 2.6 *S. maltophilia* chaperone usher pilus genes are widespread across tested clinical strains

Previous studies suggested that the *smf-1* gene is widespread in the CF clinical isolates and in environmental isolates ^45,46^. We wanted to similarly test the prevalence of *cblA* and *scs* pilin genes in clinical isolates. Testing 51 clinical isolates using PCR (see Materials and Methods), we found that the pilin genes from each of these pilus systems are present in more than 90% strains (**Fig. 8**). We also aligned the *smf-1*, *cblA*, and *scs* gene sequences from all completed clinical and environmental strains of *S. maltophilia* strain genomes listed in NCBI. We observed that the percentage identity between all the sequences was more than 90%, which indicated that these genes were conserved across strains (**Supplementary Fig. 4**). These data suggest that these pilin genes are likely important for cellular lifestyles, including potential infection, and thus possible targets for novel therapeutics.

**Figure 8.**
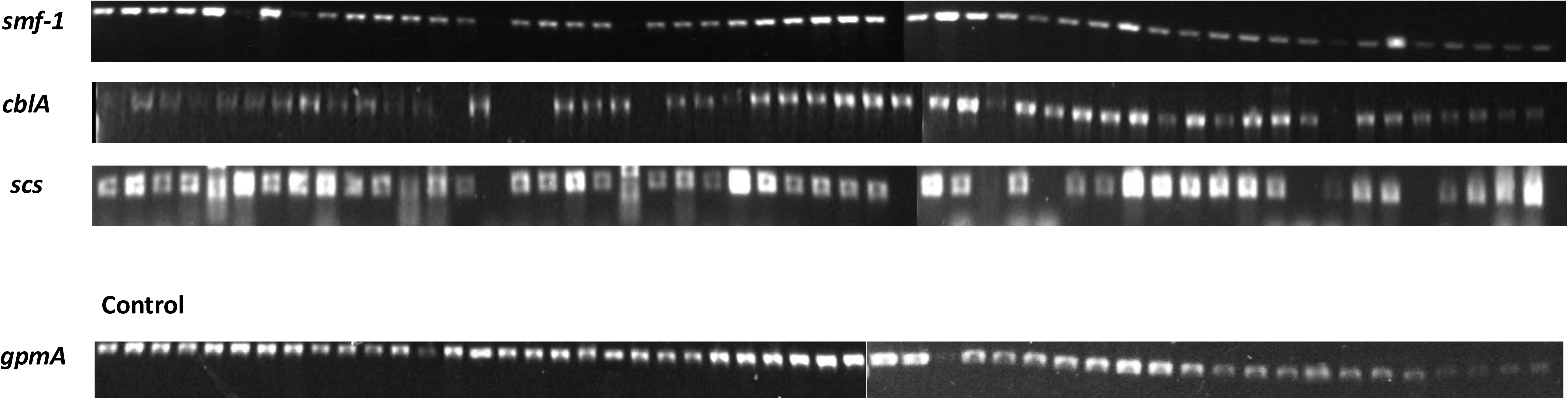
PCR analysis of *smf-1*, *cblA*, and *scs* pili genes in 51 clinical isolates. Bands observed in polyacrylamide gels show that more than 90% of tested clinical *S. maltophilia* isolates have the *smf-1*, *cblA*, and *scs* genes.

### 2.7 Clinical isolates lacking *smf-1* gene displayed lowered biofilm levels

Based on PCR analyses (**Fig. 8**), some *S. maltophilia* clinical isolates did not yield a PCR product for *smf-1* or *cblA* genes. We considered these isolates intrinsically deficient for *smf-1* or *cblA* genes. To assess whether the absence of these genes affected biofilm levels, we conducted biofilm assays of wild type, Δ*smf-1*, Δ*cblA*, and Δ*scs* along with two other clinical isolates, HSC20 (devoid of *smf-1*) and HSC139 (devoid of *cblA*). We observed that during both the early (4 hr) and late (24 hr) time points, the HSC20 (*smf-1*‾) had much lower biofilm levels compared to the wild type strain (**Fig. 9**). This result suggests that the *smf-1* pilin gene is vital in the establishment and maturation of *S. maltophilia* biofilm. In contrast, HSC139 (*cblA*‾) demonstrated biofilm levels equivalent to that of wild type at 4 hr but significantly less than wild type at 24 hr, although the isogenic deletion strain Δ*cblA* showed decreased biofilm at both time points (**Fig. 2**). This suggests that the *cblA* gene may play a significant role in sustaining long-term biofilm formation in *S. maltophilia*.

**Figure 9.**
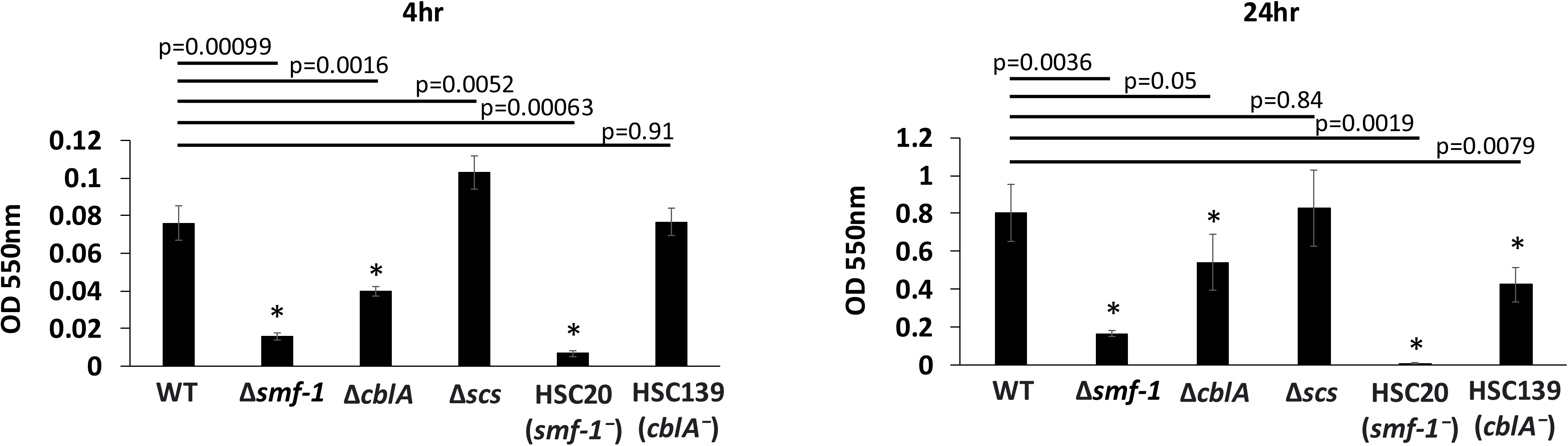
Pilus defective strains form less biofilms in the 96-well plate assay. Biofilms were formed with *S. maltophilia* wild type (WT), Δ*smf-1*, Δ*cblA*, Δ*scs*, HSC20 (*smf-1*‾), and HSC139 (*cblA*‾) for 4 hr or 24 hr. *p<0.05. These figures are the combination of three independent experiments (n=3) carried out in quadruplicate and error bars indicate standard deviations.

### 2.8 Identification of pilus genes

We next asked whether the *smf-1*, *cblA*, and *scs* pilus systems existed in other clinical and environmental *S. maltophilia* strains. By using nucleotide sequences for *smf-1*, *cblA*, and *scs* in BLAST searches of all fully sequenced *S. maltophilia* strains in the NCBI database (see Materials and Methods), we discovered that *smf-1* is contained in nearly 85% of all strains, *cblA* is found in ∼76% of strains and *scs* is found in ∼80% of all strains (**Supplementary Table 1**). Further bioinformatic searches of all fully sequenced *S. maltophilia* strains in the NCBI database (see Material and Methods) revealed a plethora of additional pilus systems dispersed among the strains. Additional putative chaperone/usher pilus systems include an F-17 like pilus (∼29% of strains), fimbriae, fimbrial biogenesis and assembly proteins (∼75-90% of strains), and additional uncharacterized pilus operons (which we designate CupA2, Cup2-Cup10, CS1) found in 1-48% of strains (**Supplementary Tables 2, 3**). Furthermore, there are individual genes annotated as “chaperone”, “usher”, “pili” or “fimbriae” found in about 73% of the strains (**Supplementary Table 2**).

In addition to chaperone/usher pilus systems, we discovered two type IV pilus systems: a canonical type IV and a Tad pilus, found in 75.86% and 75% of strains, respectively (**Supplementary Table 3**). Type IV pili have been described previously in *S. maltophilia*; it is unclear whether these pili correspond to the two loci we identified ^47,48^. We also found a putative type II pilus assembly system in about 72% of the strains (**Supplementary Table 3**).

Finally, we discovered the presence of putative afimbrial adhesins (*afaD* and *afaD2*) (**Supplementary Table 3**) scattered across *S. maltophilia* strains (45.21% and 14.78% respectively). In other Gram-negative bacteria, afimbrial adhesins have been reported to bind to several host receptors ^49,50^. To our knowledge, these afimbrial adhesins have not hitherto been identified and are currently uncharacterized in *S. maltophilia*.

## 3. Discussion

*S. maltophilia* is emerging as a major multidrug-resistant nosocomial pathogen. Surface associated appendages such as pili and flagella play a critical role in bacterial adherance to surfaces, contributing to colonization and perhaps biofilm formation. Like many other Gram-negative bacteria, *S. maltophilia* utilizes biofilm formation during pathogenesis. In fact, more than 98% of *S. maltophilia* isolates can form a biofilm and 46% of these are strong biofilm producers^11^. In this study we found that *S. maltophilia* encodes an extensive repertoire of pilus-associated genes, highlighting the importance of adhesive structures in this organism (**Supplementary Tables 1-3)**.

Among all the *S. maltophilia* pilus systems listed, the three most widely spread across strains are *smf-1*, which expresses the SMF-1 pili; *cbl*, which has similarity to the cable pilus in *Burkholderia cepacia* ^51,52^; and *scs*, which has similarity to Csu pilus in *Acinetobacter baumannii* ^38,39^. *cblA* is used a DNA marker for virulent isolates of *B. cepacia* infecting CF patients ^51^, and Csu pili in *A. baumanii* have been reported to be indispensable for biofilm growth^53^. The conservation of these homologous systems in *S. maltophilia* strains suggests that they may play similarly important roles in surface attachment and persistence.

In this study, functional characterization of the *scs* locus revealed its context-dependent role in biofilm formation. We observed that deletion of *scs* alone had little effect on biofilm formation of this bacterium (**Fig. 2**). However, pronounced defects emerged when the *scs* deletion was combined with other pilus mutations in both abiotic surfaces and infection relevant models (**Figs. 2,3**). These findings are consistent with our earlier observations using *smf-1* and *cblA* mutants ^30^. The reduction in these biofilm phenotypes was seen without affecting planktonic growth (**Supplementary Fig. 1 and 2**), indicating a bacterial attachment specific effect. Our results highlight potential crosstalk and combinatorial effects of surface expressed pili in *S. maltophilia*.

In addition to attachment, motility is a key feature of many pathogenic bacteria, allowing them to migrate to favorable niches, interact with other microorganisms, evade host immune response, and cause disease ^54,55^. Motility allows bacteria to find appropriate environmental conditions to transition to sessile communities for biofilm differentiation ^56^. Given the established role of pili, in surface-associated motility, we examined whether deletion of pilus genes affected motility behaviors in *S. maltophilia*. A comparable pattern was observed for both swimming and swarming motilities, aligning with the strain that exhibited the highest level of biofilm formation. The higher motility of the Δ*scs* (**Fig. 5**), in keeping with its increased biofilm capacity (**Fig. 2**), suggests an increased efficiency of this mutant to find optimal environments for pathogenesis. The increased motility of the Δ*scs* mutant was accordingly associated with an increase in *fliC* gene expression (**Fig. 6**). Together, these observations support a model in which loss of the *scs* pilin enables compensatory expression of alternative pili (Smf-1, Cbl, and perhaps others). We propose a regulatory mechanism where these compensatory events then modulate flagellar gene regulation, resulting in increased *fliC* expression and motility in the Δ*scs* mutant. This hypothesis is further supported by the fact that the combined loss of *scs* and *smf-1* pili suppresses motility (**Fig. 5**) and biofilm formation (**Fig. 2**), and shows a corresponding decrease in flagellar expression (**Fig. 6**). Our results highlight the need for future studies exploring this connection between pilus and flagellar expression, and whether it impacts pathogenesis and infection.

Data presented in **Fig. 2** and **Fig. 3** further highlight the complex coordination of pilus expression in *S. maltophilia*. Specifically, while deletion of *scs* alone had little effect on biofilm formation, the *scs* mutation in combination with other pilus mutations resulted in the lowest biofilm levels observed in both LB and SCFM2 media (**Supplementary Fig. 5**). Similarly, as observed previously ^30^, the presence of chaperone-usher pili in these strains correlates directly with their biofilm-forming ability. While Δ*scs* (retaining *smf-1* and *cblA*) shows robust chaperone/usher piliation, Δ*smf-1* Δ*scs* (retaining *cblA*) exhibits little to none, and Δ*cblA* Δ*scs* shows intermediate piliation, mirroring its biofilm phenotype (**Fig. 4**). These observations indicate that piliation phenotypes cannot simply be attributed to the additive effects of individual pilin gene mutations. The absence of compensatory effects among pilin genes points to a more intricate regulatory network, potentially involving transcriptional coordination or physical and mechanical interactions between these pilus systems.

We also observed virulence effects of pilus mutation in the *G. mellonella* infection model (**Fig. 7**). Bacterial pathogens often exhibit an inverse regulatory relationship between virulence factor secretion and biofilm formation, allowing them to adapt between acute and chronic infection states ^57^ ^58,59^. In our previous manuscript ^30^, we observed that pilus-deficient *S. maltophilia* mutants displayed increased virulence in a *G. mellonella* infection model, suggesting a potential trade-off between biofilm formation and acute infection. Our current study reinforces this pattern, as Δ*smf-1* Δ*scs* mutants again exhibit heightened virulence compared to the wild-type strain. This consistent phenotype suggests that pilus loss may alter regulatory or metabolic pathways that promote dissemination and acute toxicity. These findings point toward a broader shift in infection strategy triggered by disruption of pilus-associated structures. The high prevalence of these pilus genes across *S. maltophilia* strains (**Fig. 8**), and the high percentage of relatedness across strains (**Supplementary Fig. 4**), further indicates that their putative importance for attachment and biofilm formation in a variety of environments, including during infection. Within this prevalence, the clinical strains that lacked either the *smf-1* or *cblA* gene were notably deficient in forming robust biofilms (**Fig. 9**), further supporting the functional importance of these pili in surface attachment and biofilm development.

Apart from the *smf-1*, *cblA*, and *scs* pili structures, we found the presence of additional chaperone/usher pilus operons, putative pilin subunits, putative fimbrial assembly and biogenesis proteins, type IV pili, Tad like pili, afimbrial adhesins, putative type II pilus assembly structures, and other individual pilus genes and adhesive structures, within the pan genome (**Supplementary Tables 2-3**). We speculate that the diversity of these structures makes the bacteria extremely versatile and robust in carrying out adherence and biofilm formation. Additionally, the numerous pili could be involved in broad host cell tropism, transferring genetic material, mediating phage transduction, and performing numerous motility activities^60^. Increased attachment to various host cells or environmental surfaces could lead to increased disease transmission and progression^61^. It could also assist in evading host immune defenses and establishing greater number of infections.

Importantly, the extensive repertoire of adhesive and pilus-associated systems provides a plausible explanation for the context-dependent role of the *scs* locus observed in this study. Crosstalk among multiple adhesive structures may allow functional compensation or modulation such that deletion of *scs*, either as a single mutation or in combination with other loci, results in distinct phenotypic outcomes depending on the genetic and environmental context.

Overall, our study sheds light on the functional role of the *scs* locus as a key determinant of attachment in *S. maltophilia*, revealing that its contribution is highly context dependent. Rather than acting in isolation, *scs* likely functions within a complex network of adhesive and pilus-associated systems, where crosstalk and functional redundancy shape attachment and biofilm phenotypes. These findings contribute to a better understanding of how this pathogen works within the host environment to initiate infection. Further elucidating pilus production and regulation, and how these impact other virulence activities in *S. maltophilia*, can open new avenues for development of distinctive therapeutic strategies to counter this deadly pathogen.

## 4. Acknowledgements.

We acknowledge Jordan Hall, Alli Beard, Oliver Harrigan, Layla Ramos-Hegazy, Harjot Kaur, Eli Klein, Cierra Isom, and Blake Fort for their assistance. This work was supported by RSFG and BRIDGE funding from IUPUI to G. Anderson, and by the Life Sciences Summit Initiative from Purdue University and the Hypothesis Fund Award to S. Mattoo. We also acknowledge Dr. Valerie Waters for the kind donation of clinical isolates. We kindly thank Dr. Quyen Quoc Hoang and Derrick A. Gray for helping us with the TEM experiments.

## 5. Credit Authorship Contribution Statement

**Radhika Bhaumik-** Conceptualization, Methodology, Investigation, Formal analysis, Data curation, Writing – original draft, Writing – review & editing; **Gregory G. Anderson-** Conceptualization, Methodology, Supervision, Project administration, Formal analysis, Writing – review & editing, Funding acquisition; **Seema Mattoo-** Conceptualization, Formal analysis, Resources, Supervision, Writing – review & editing, Project administration, Funding acquisition.

## 6. Ethics Statement and Declaration of AI assisted writing

All scientific content, data interpretation, and conclusions were generated, verified, and approved by the authors. However, the authors used artificial intelligence–assisted language editing tools to improve clarity, grammar, and overall readability of the manuscript.

## 9. Supplementary Figure Legends

**Supplementary Fig. 1.**
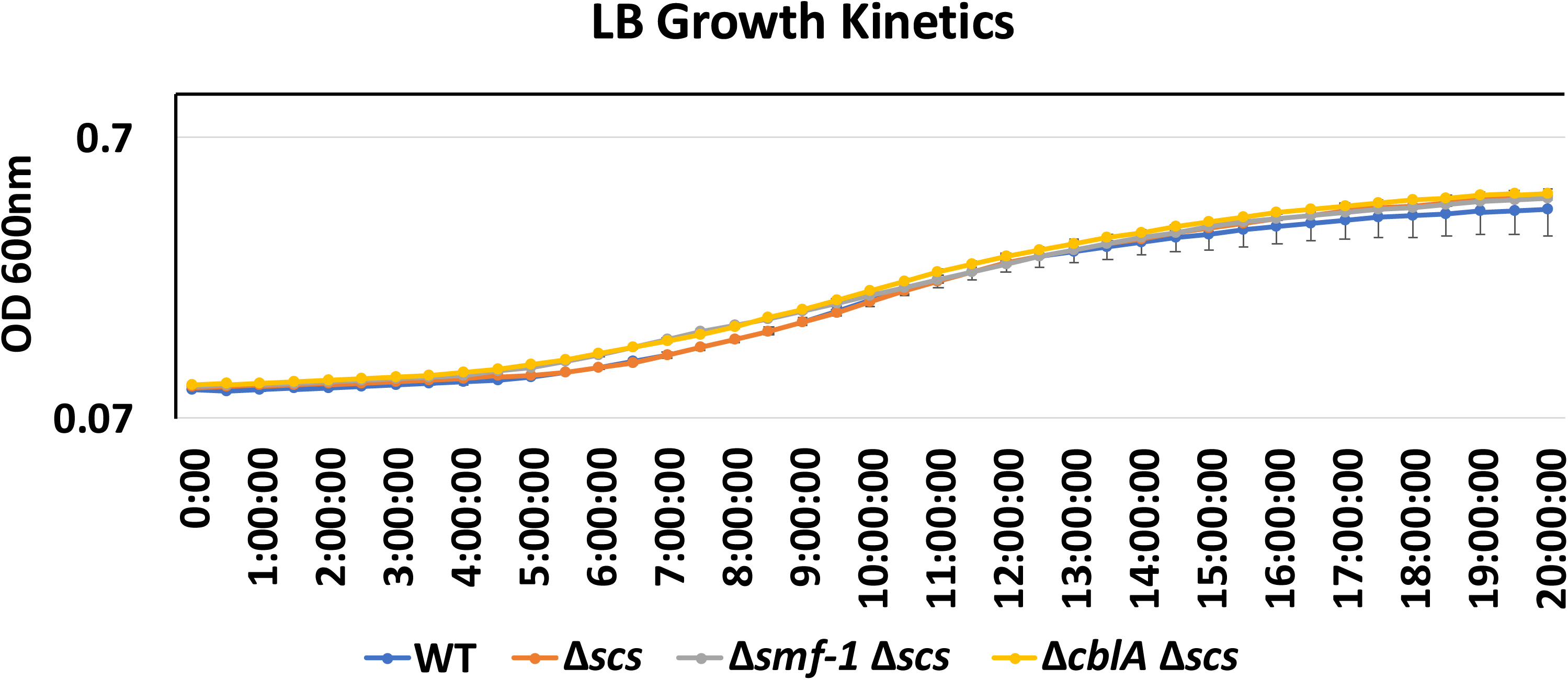
Pilin gene deletions do not affect bacterial growth in LB medium. Growth kinetics of WT, Δ*scs,* Δ*smf-1* Δ*scs*, and Δ*cblA* Δ*scs*, in LB medium. Absorbance was measured for 20 hr at OD₆₀₀. This figure is representative of a combination of three independent experiments (n=3) carried out in quadruplicate and error bars indicate standard deviations. No significant difference was found. (p>0.05).

**Supplementary Fig. 2.**
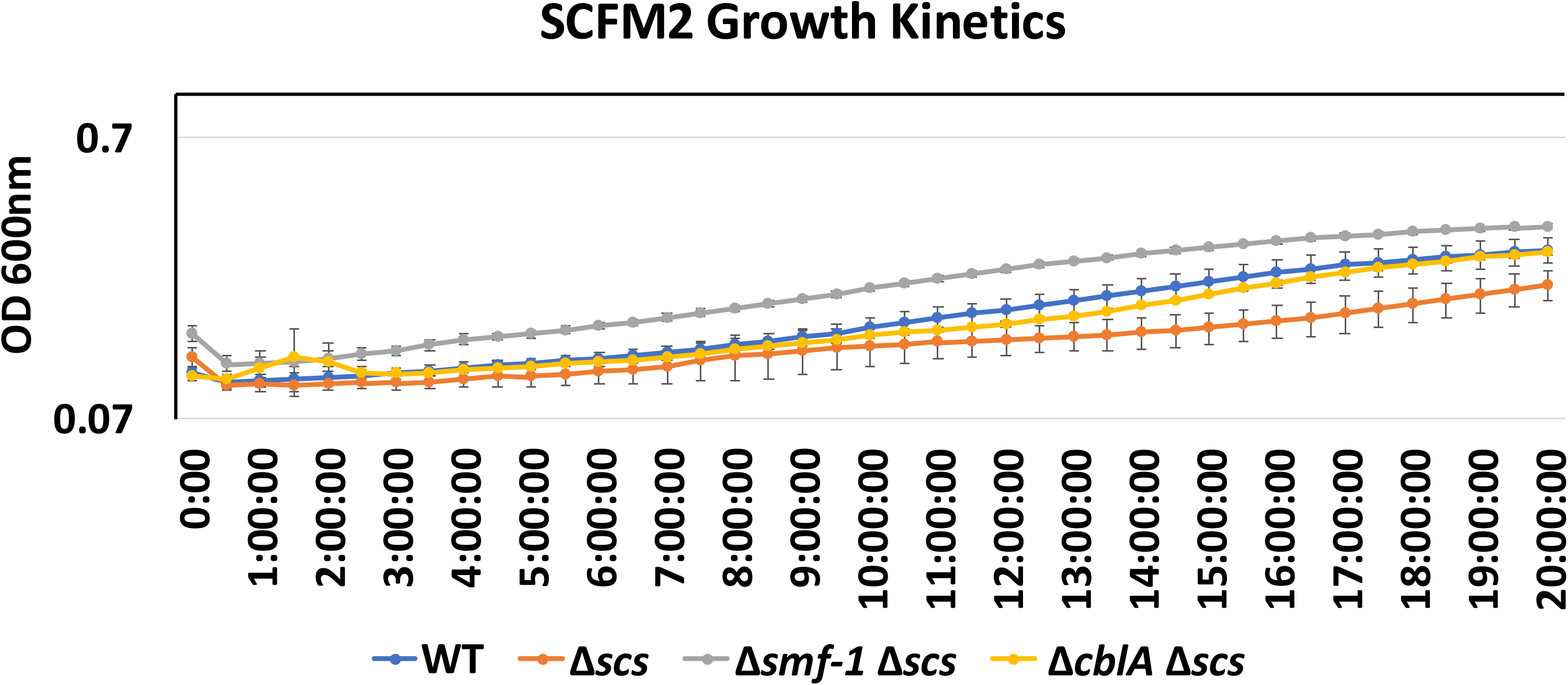
Pilin gene deletions do not affect bacterial growth in SCFM2 medium. Growth kinetics of WT, Δ*scs,* Δ*smf-1* Δ*scs*, and Δ*cblA* Δ*scs*, in SCFM2 medium. Absorbance was measured for 20 hr at OD₆₀₀. This figure is representative of a combination of three independent experiments (n=3) carried out in quadruplicate and error bars indicate standard deviations. No significant difference was found. (p>0.05).

**Supplementary Fig. 3.**
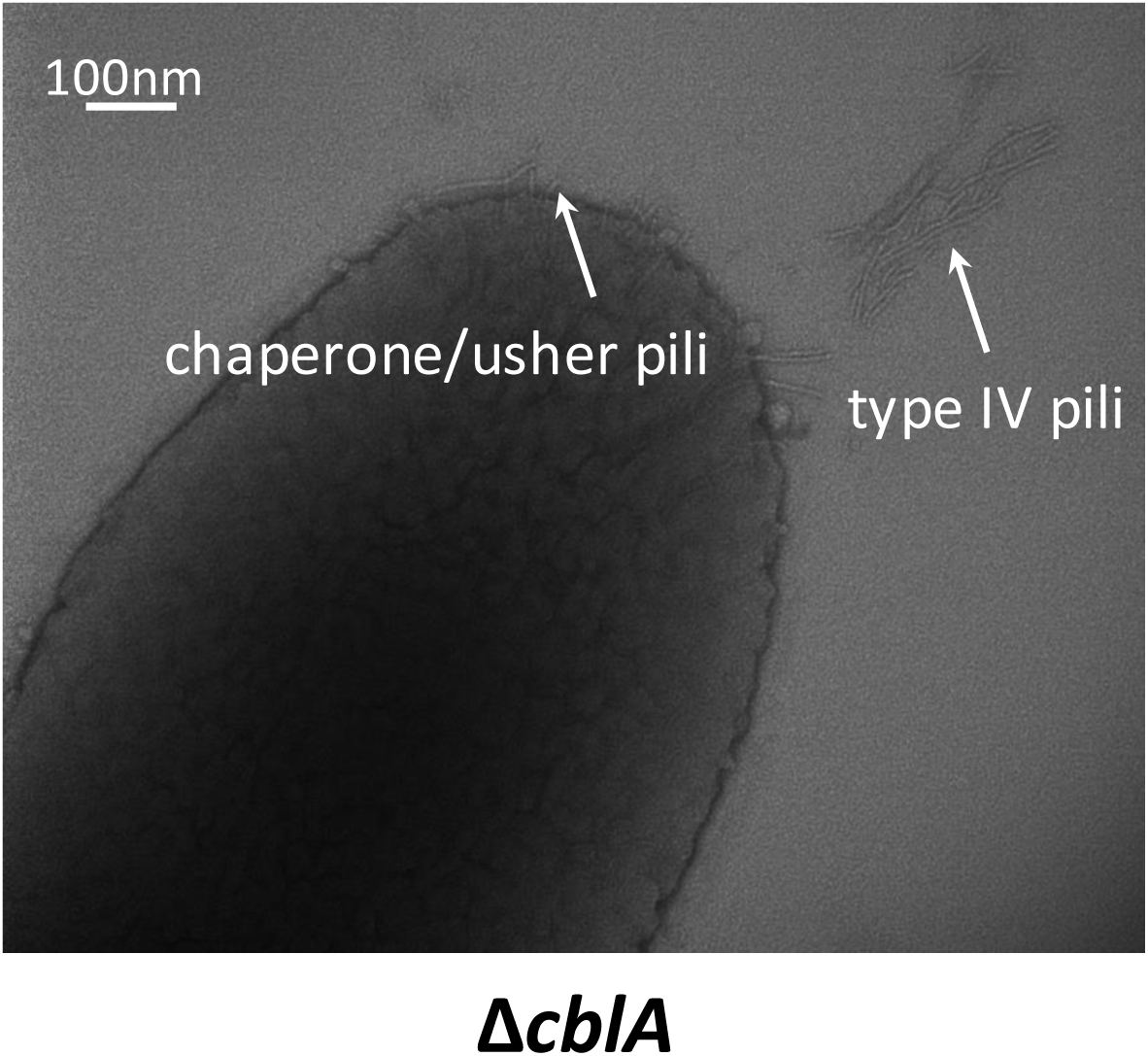
TEM studies show the presence of chaperone/usher pili (short and hairy) and type IV pili (long) in Δ*cblA* strain. TEM images of Δ*cblA* strain. Shorter pili correlate to chaperone/usher pili and longer structures correlate with type IV pili, as seen in previous studies. See text for details. Scale bars are as indicated on the micrographs.

**Supplementary Fig. 4A.**
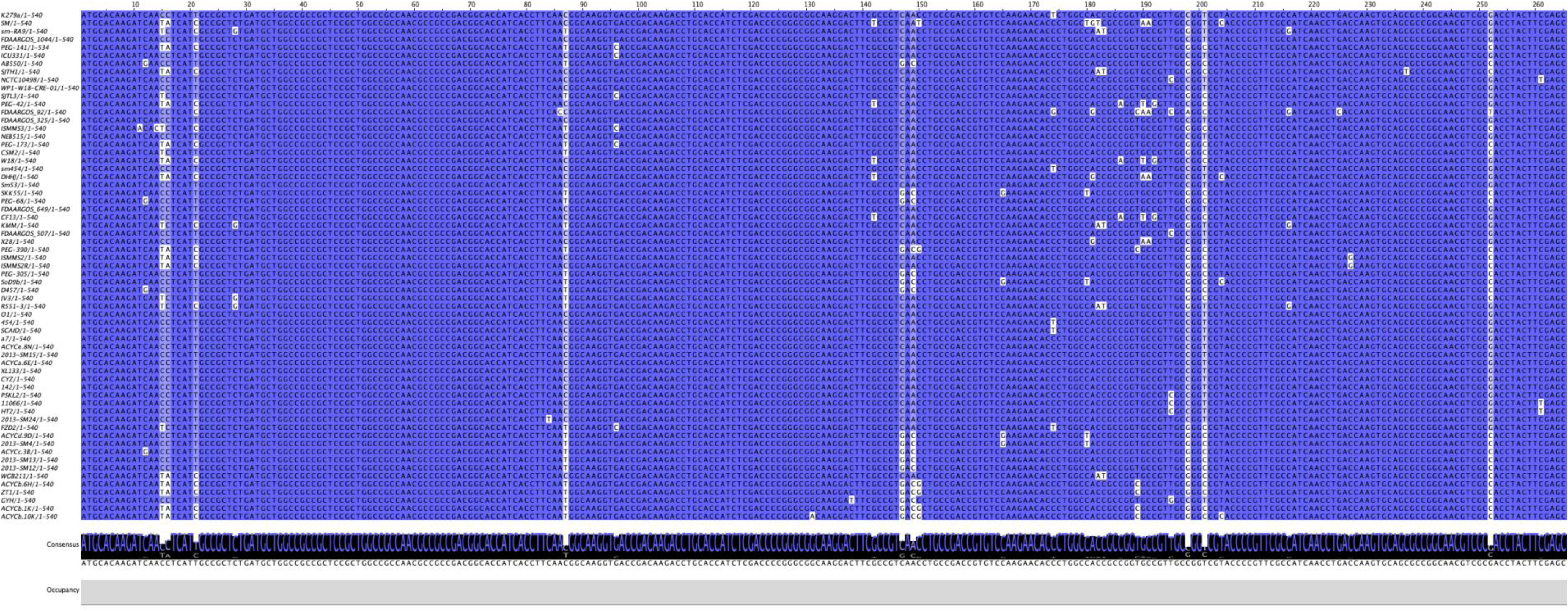
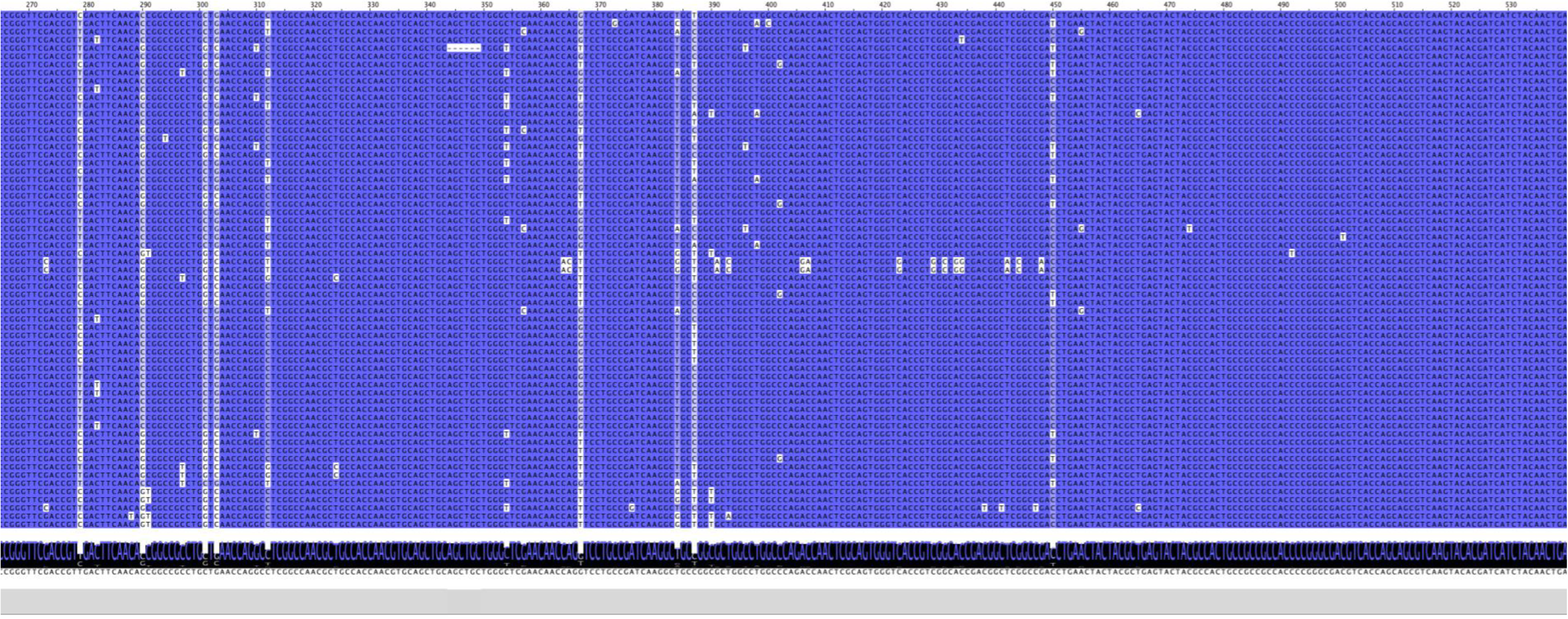
Multiple sequence alignment of the *smf-1* gene across fully sequenced *Stenotrophomonas maltophilia* strains. Nucleotide sequences of the *smf-1* gene from all fully sequenced *S. maltophilia* strains available in NCBI were aligned using Jalview. Shades of blue indicate the degree of sequence identity at each nucleotide position, with darker blue representing higher conservation and lighter blue representing lower conservation across strains.

**Supplementary Fig. 4B.**
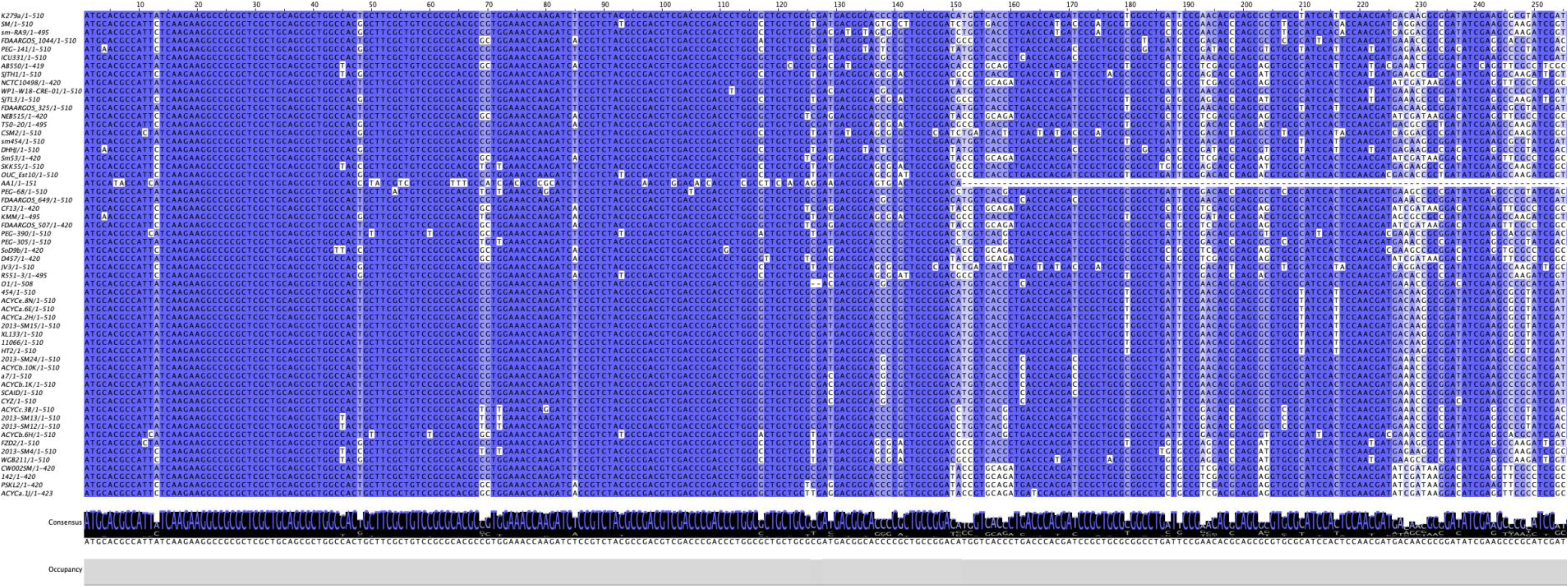
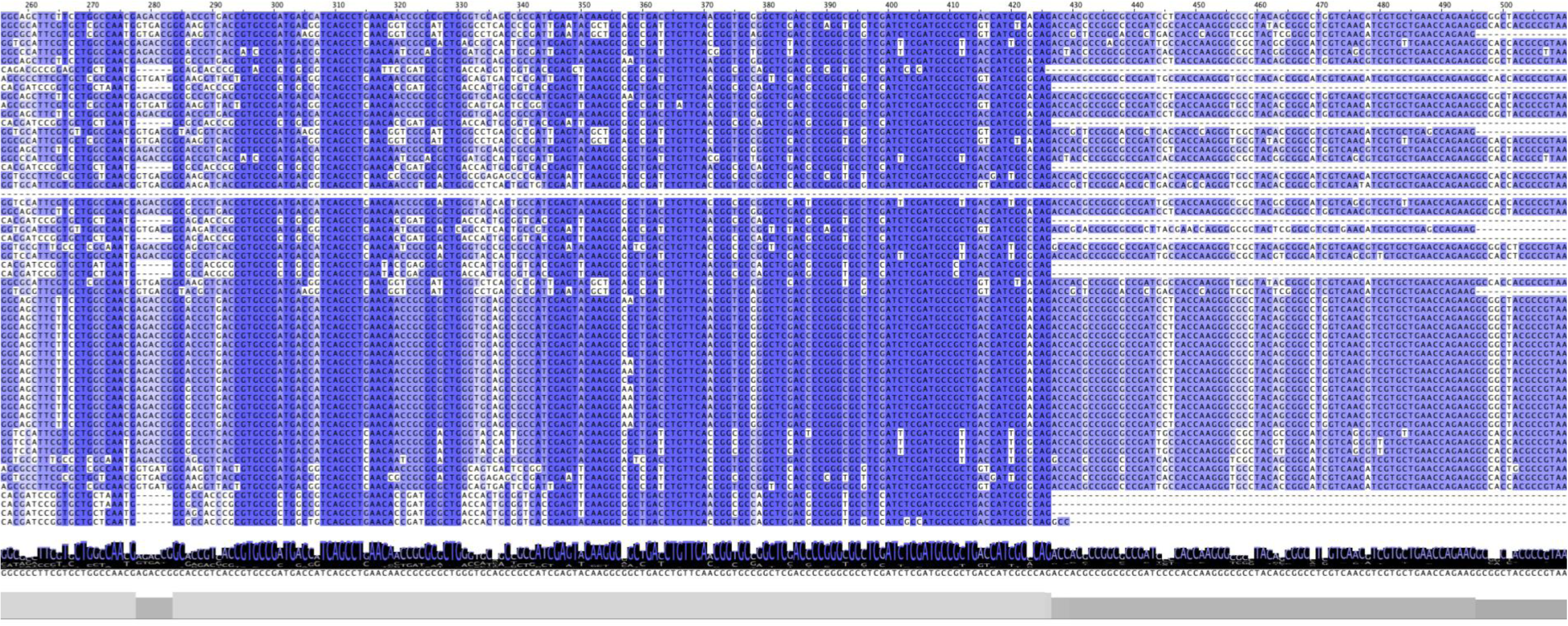
Multiple sequence alignment of the *cblA* gene across fully sequenced *Stenotrophomonas maltophilia* strains. Nucleotide sequences of the *cblA* gene from all fully sequenced *S. maltophilia* strains available in NCBI were aligned using Jalview. Shades of blue indicate the degree of sequence identity at each nucleotide position, with darker blue representing higher conservation and lighter blue representing lower conservation across strains.

**Supplementary Fig. 4C.**
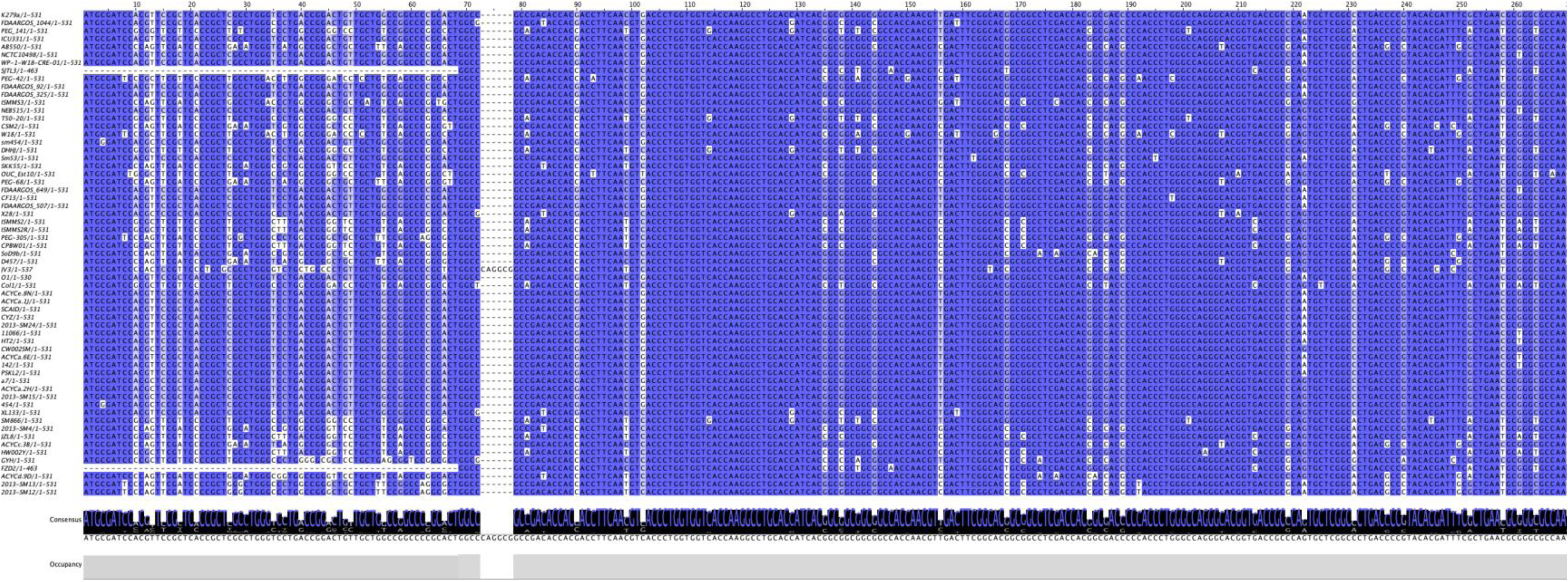
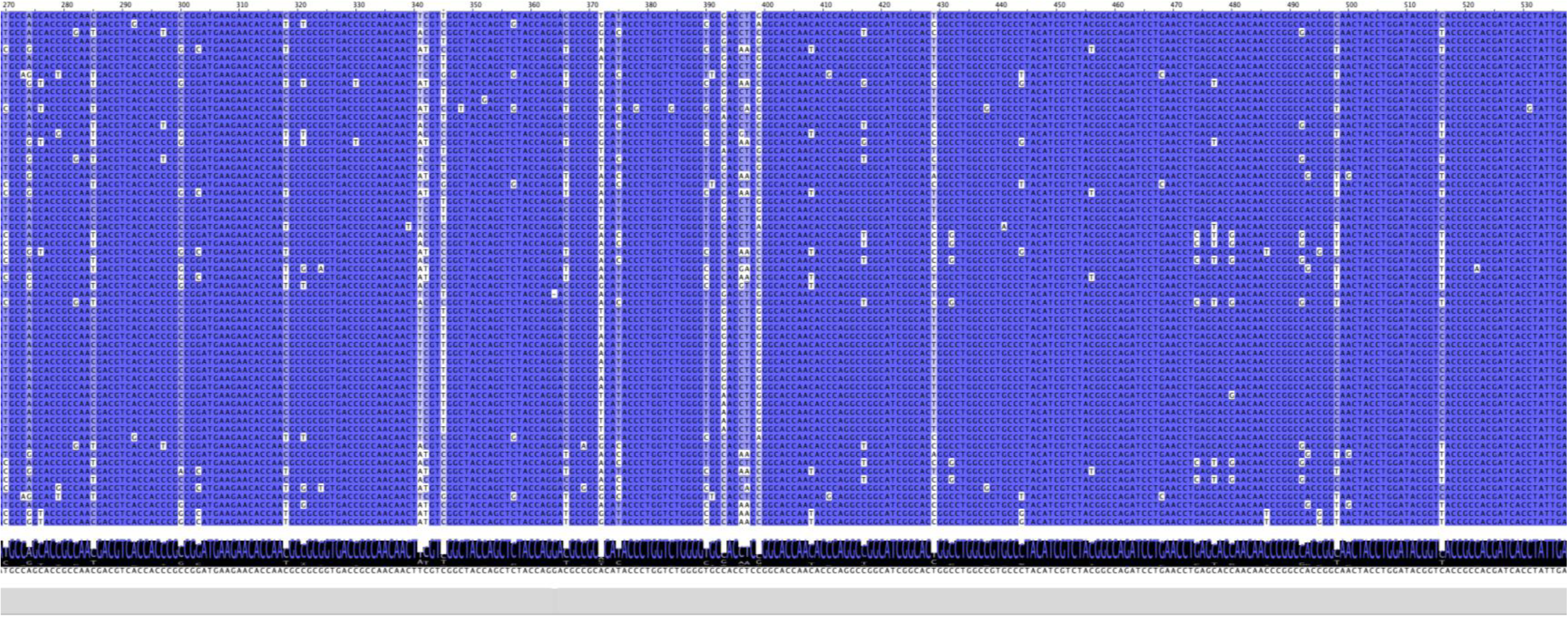
Multiple sequence alignment of the *scs* gene across fully sequenced *Stenotrophomonas maltophilia* strains. Nucleotide sequences of the *scs* gene from all fully sequenced *S. maltophilia* strains available in NCBI were aligned using Jalview. Shades of blue indicate the degree of sequence identity at each nucleotide position, with darker blue representing higher conservation and lighter blue representing lower conservation across strains.

**Supplementary Fig. 5.**
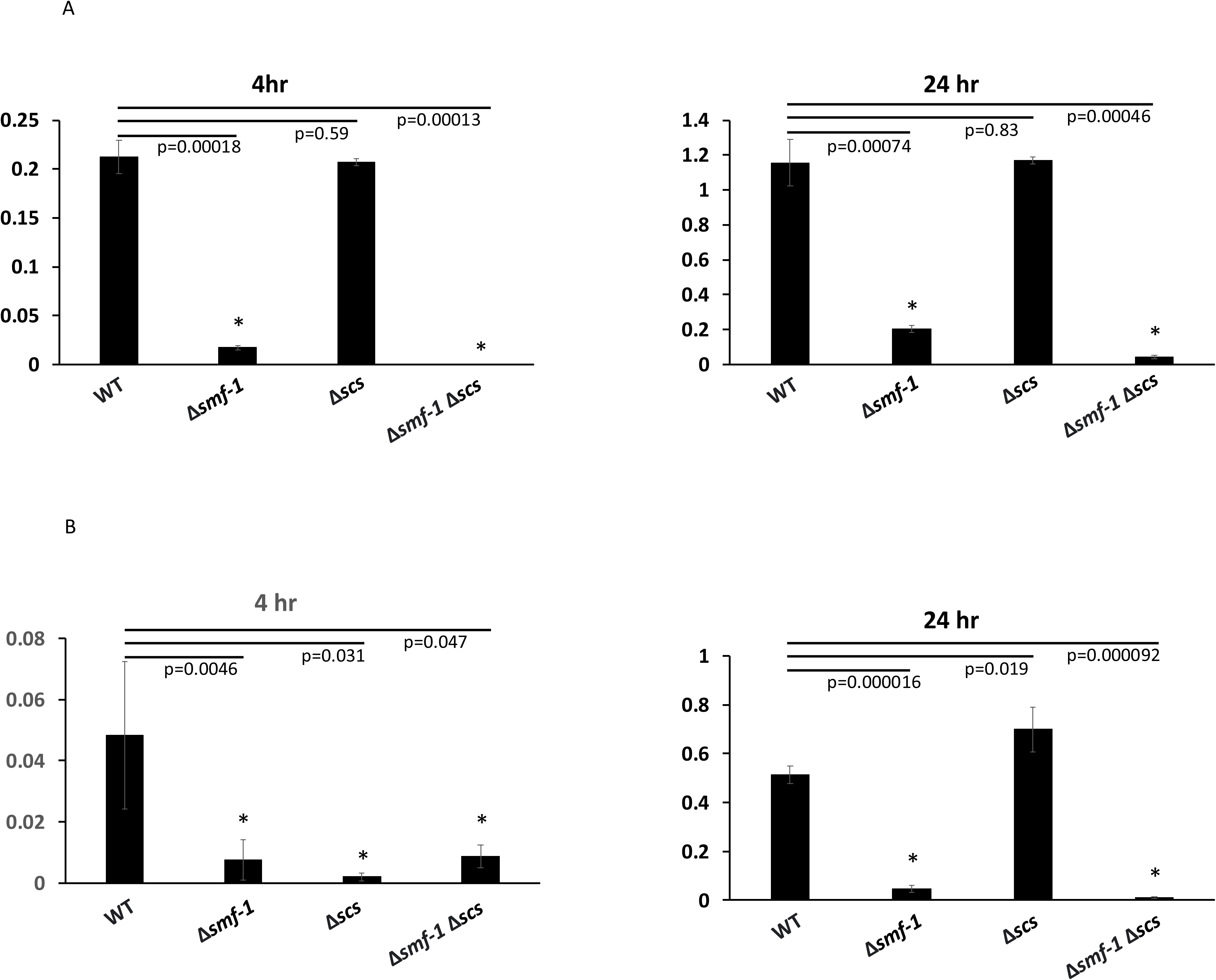
Isogenic deletion of pili genes dramatically reduces *S. maltophilia* biofilm formation in LB medium. Biofilms of *S. maltophilia* wild type (WT), Δ*smf-1,* Δ*scs* and double mutant Δ*smf-1* Δ*scs* in (A) LB medium and (B) SCFM2 medium for 4 hr (left panels) or 24 hr (right panels). *p<0.05. These figures are the combination of three independent experiments (n=3) carried out in quadruplicate and error bars indicate standard deviations.

## 10. Supplementary Table Legends

**Supplementary Table 1.**
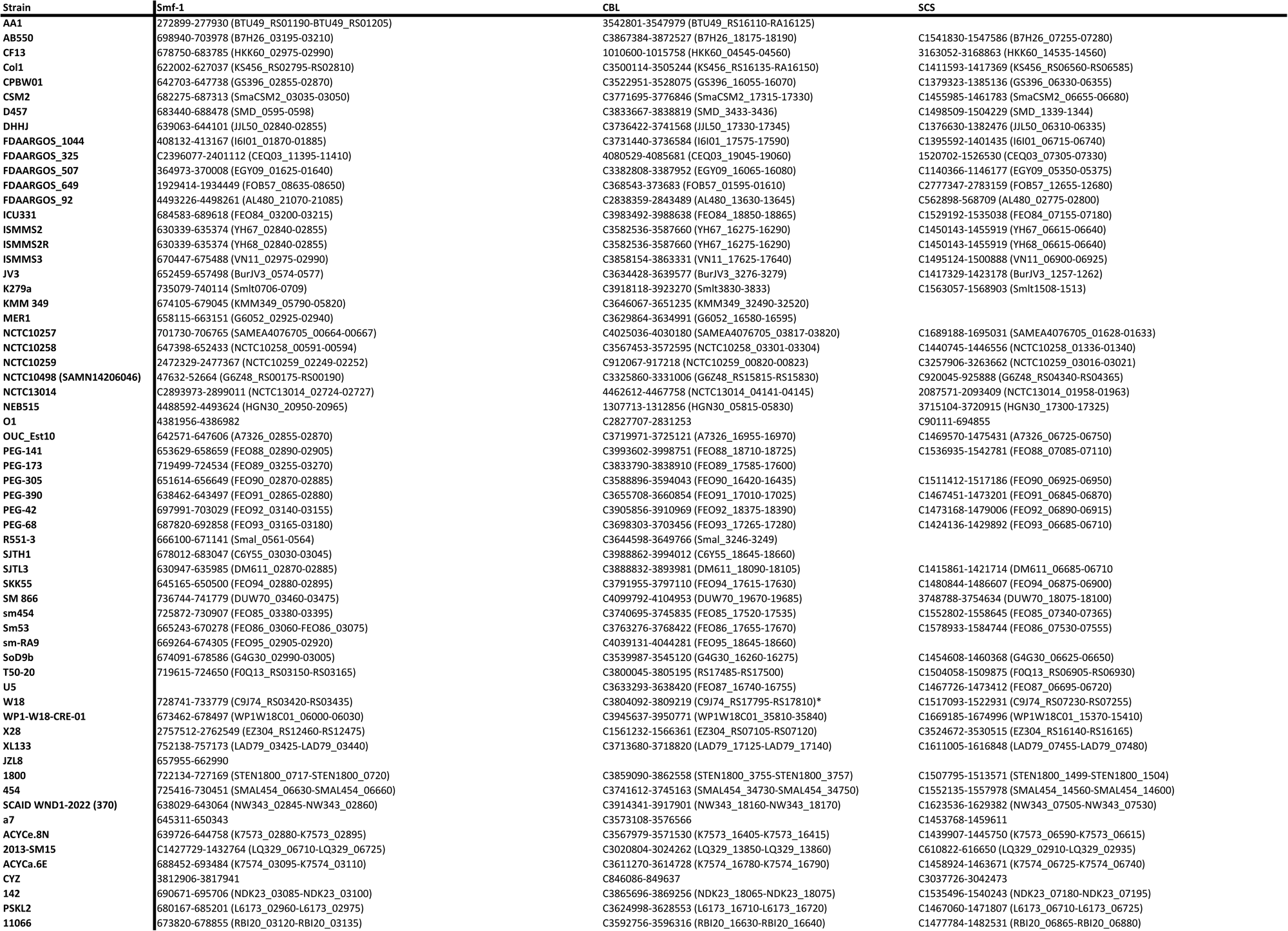
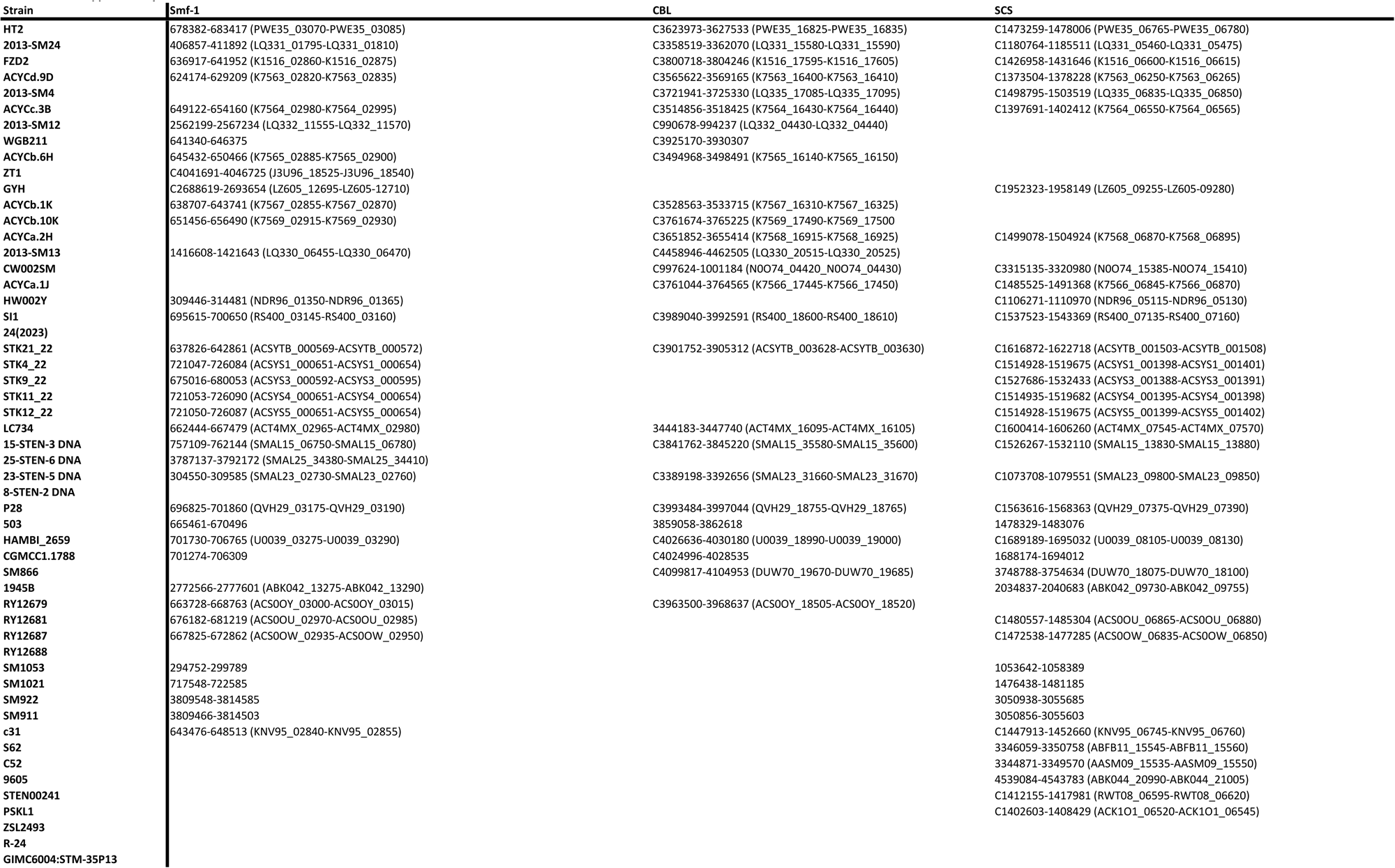
Distribution of major chaperone–usher pilus operons in *Stenotrophomonas maltophilia* strains. Genomic loci corresponding to the major chaperone–usher pilus operons Smf-1, CBL, and SCS across fully sequenced *Stenotrophomonas maltophilia* strains obtained from the NCBI database. Numbers listed indicate the genetic locus positions within each strain genome. A “C” preceding the locus number range denotes that the locus is located on the complementary strand of the genome. Gene designations shown in parentheses represent the annotated genetic nomenclature of each locus in the NCBI genome database.

**Supplementary Table 2.**
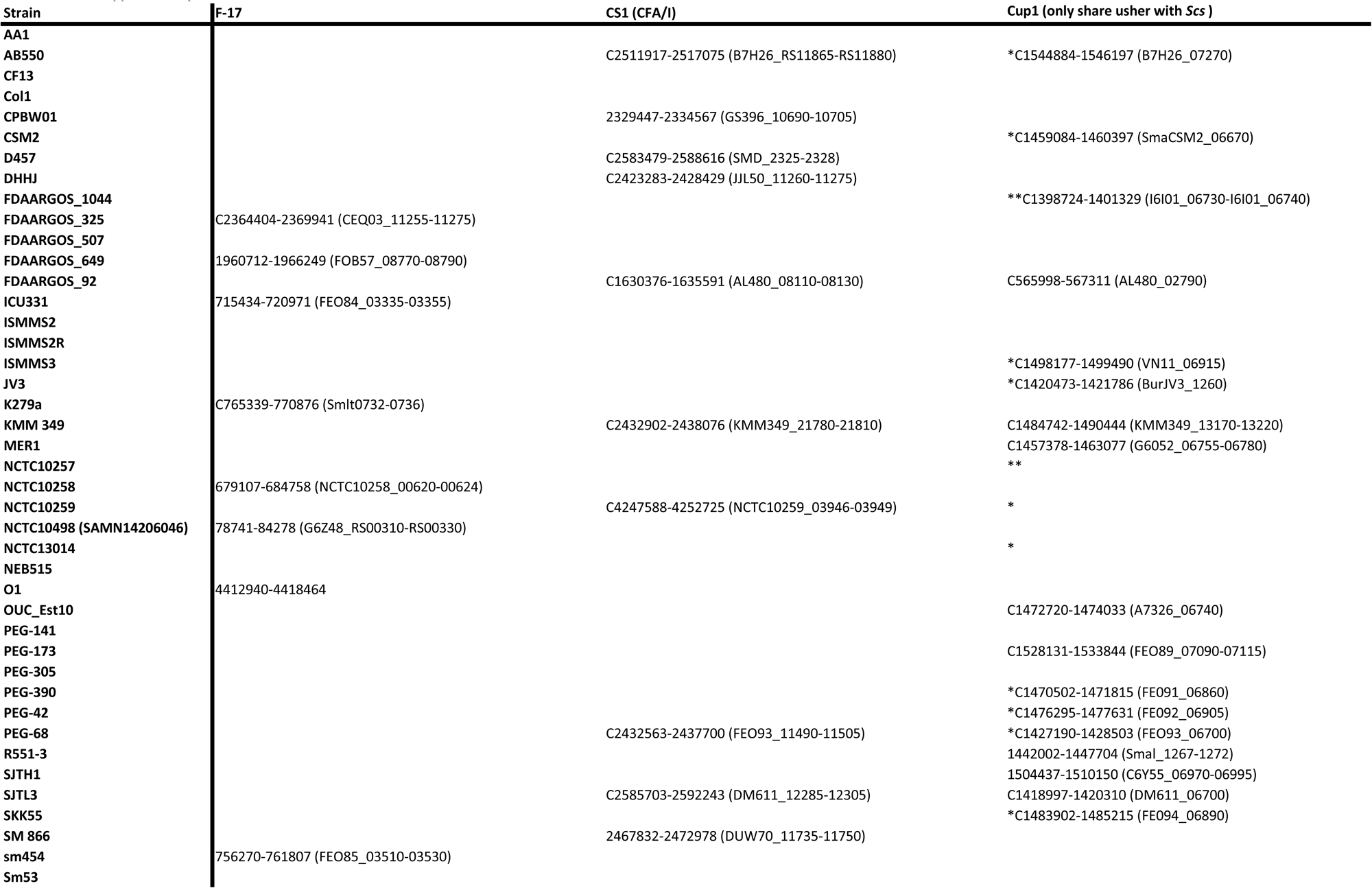
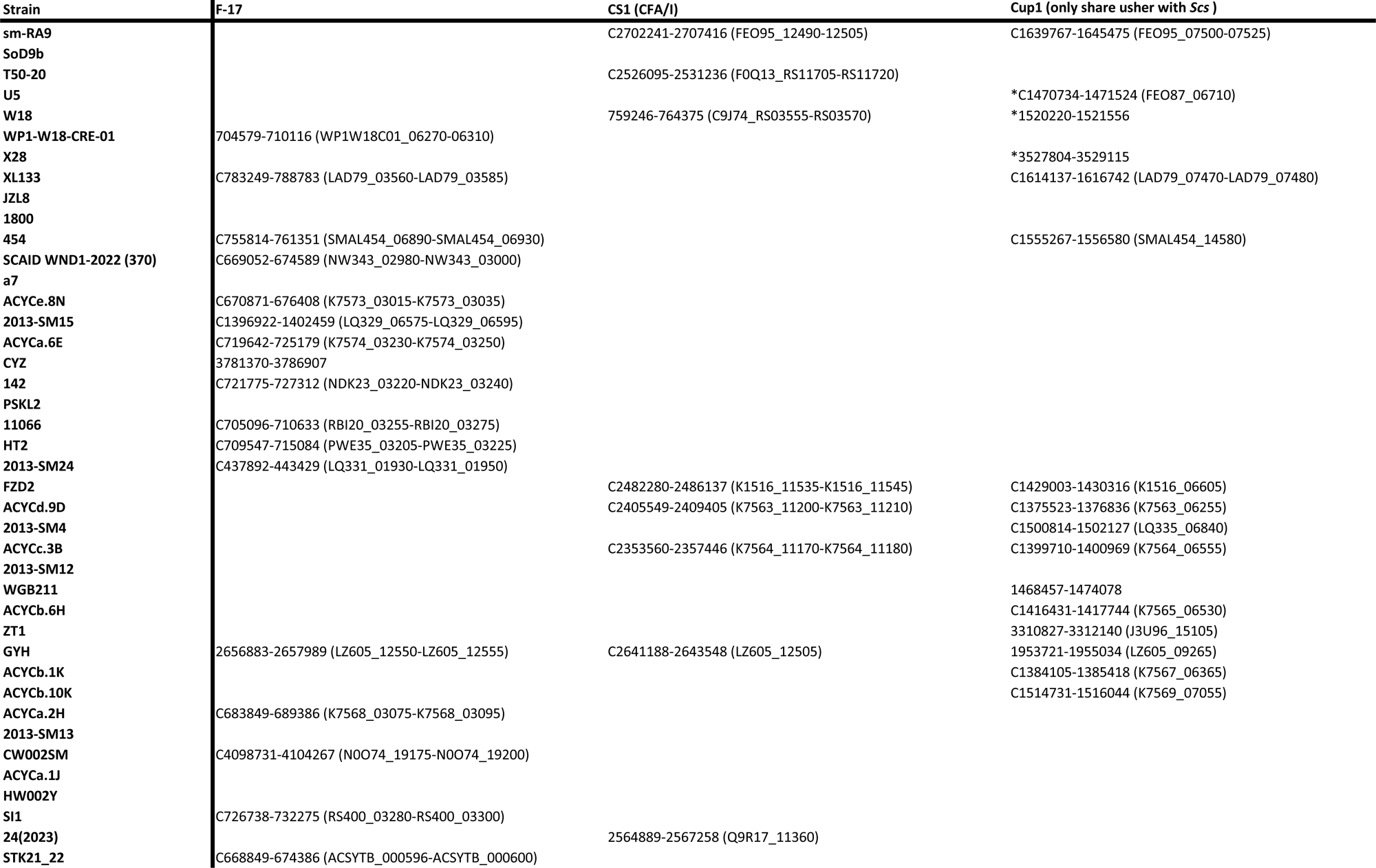
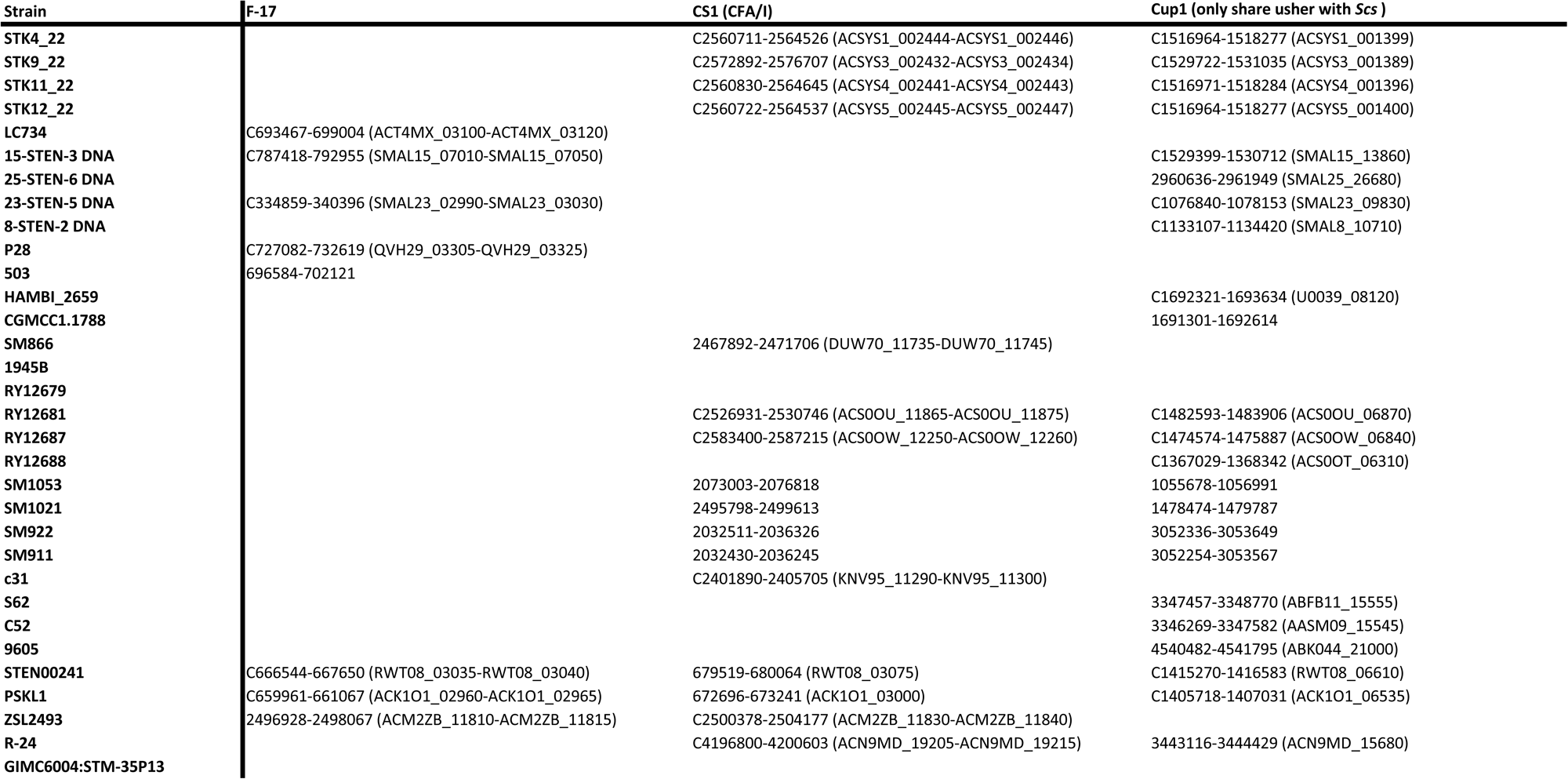
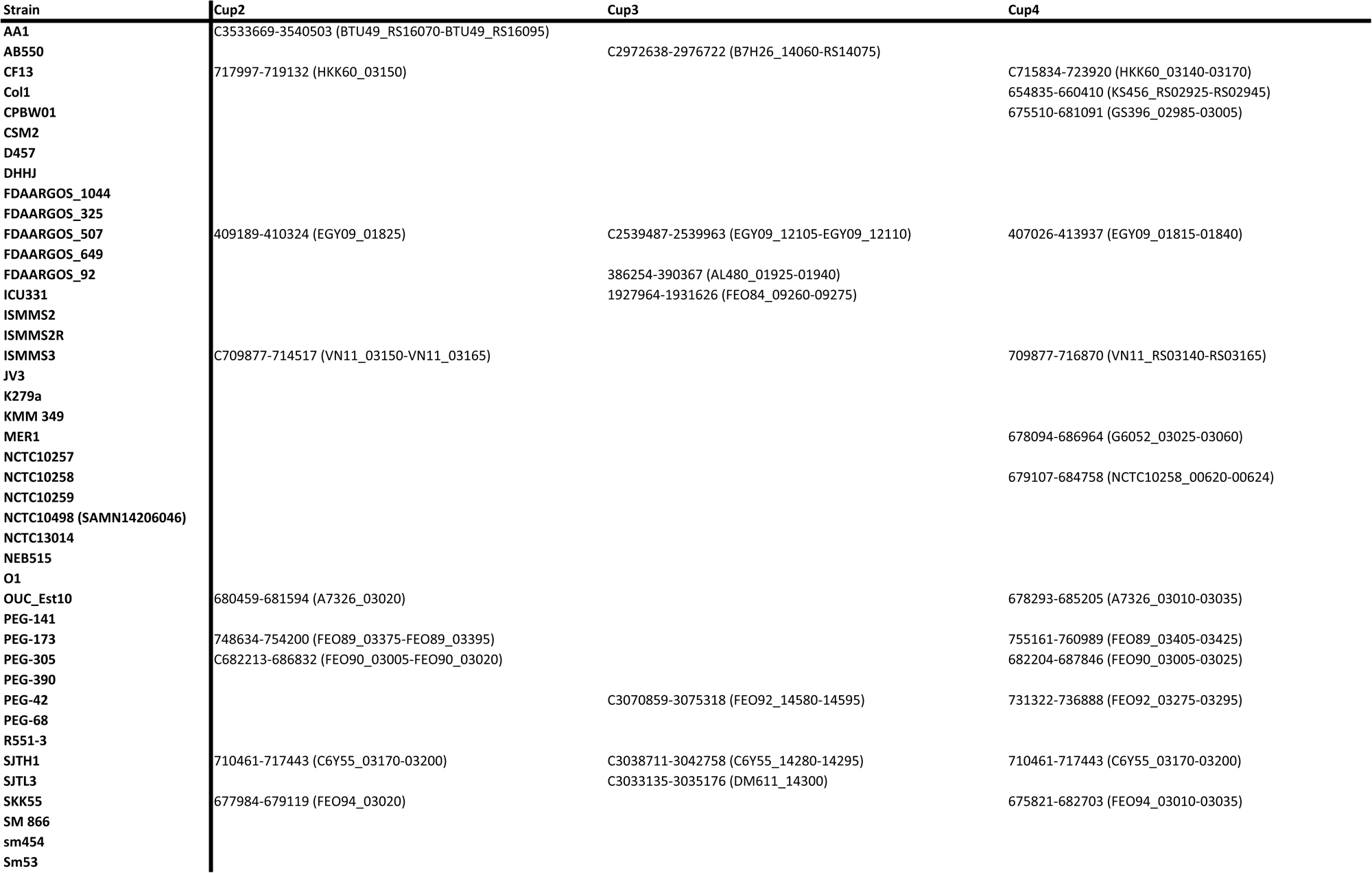
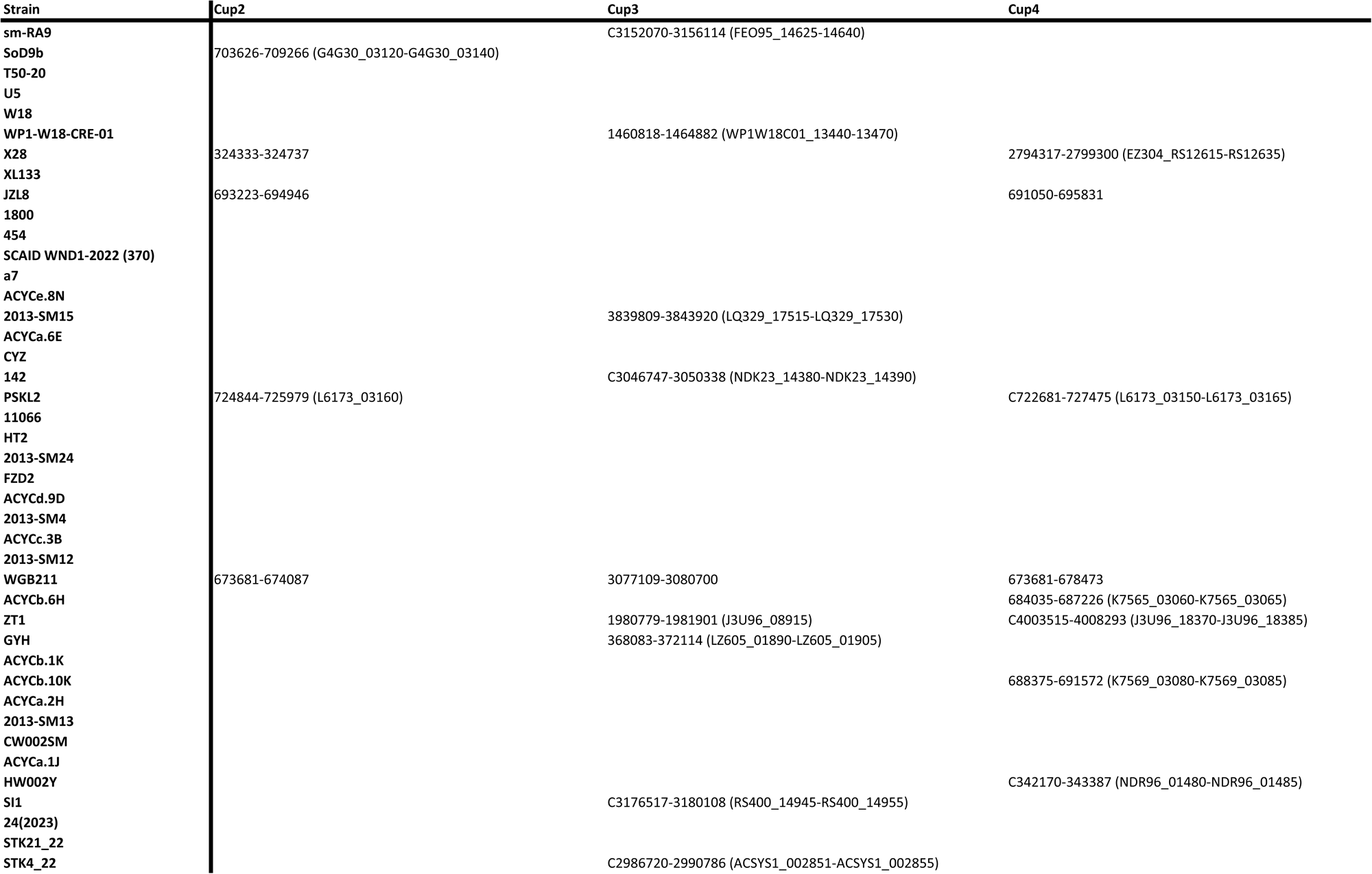
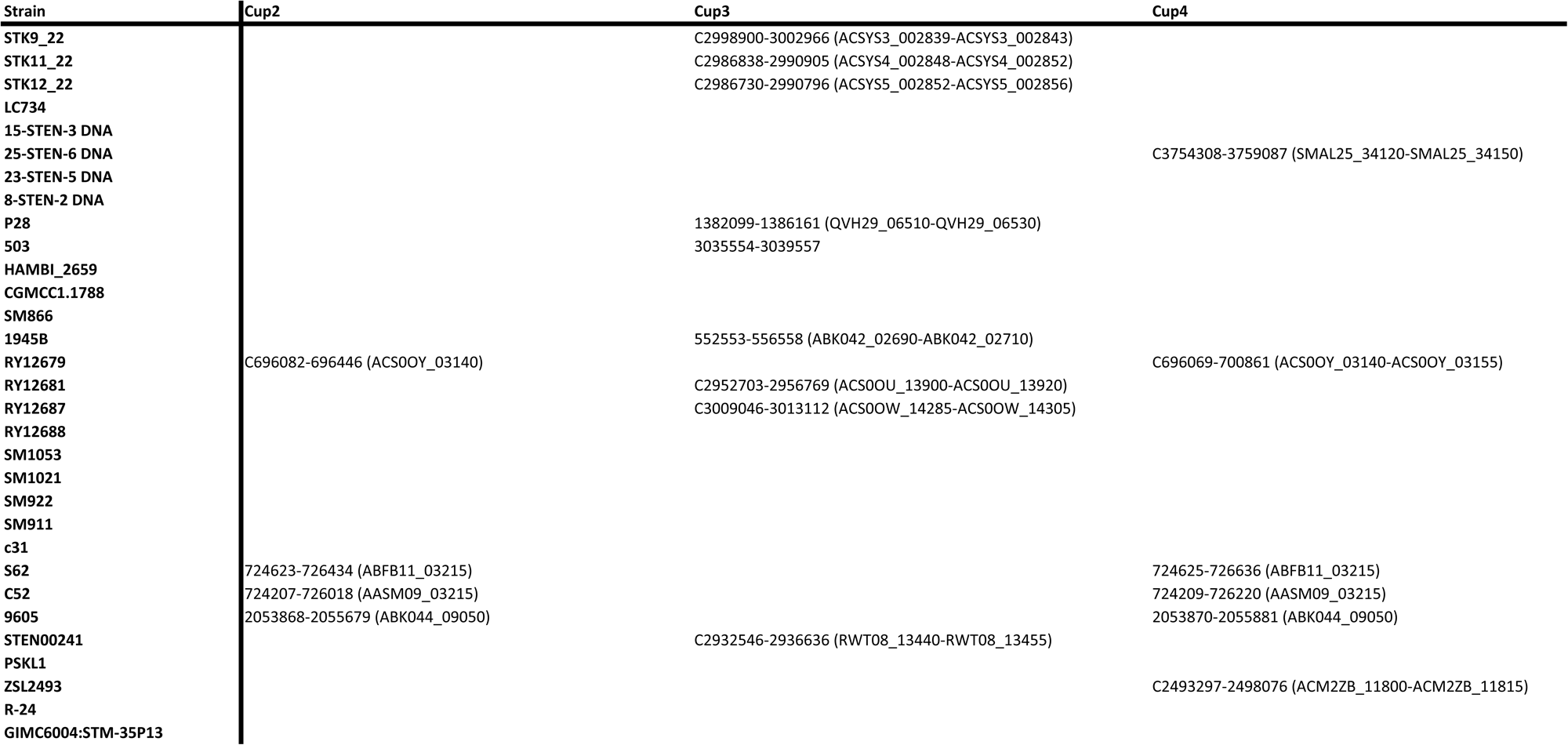
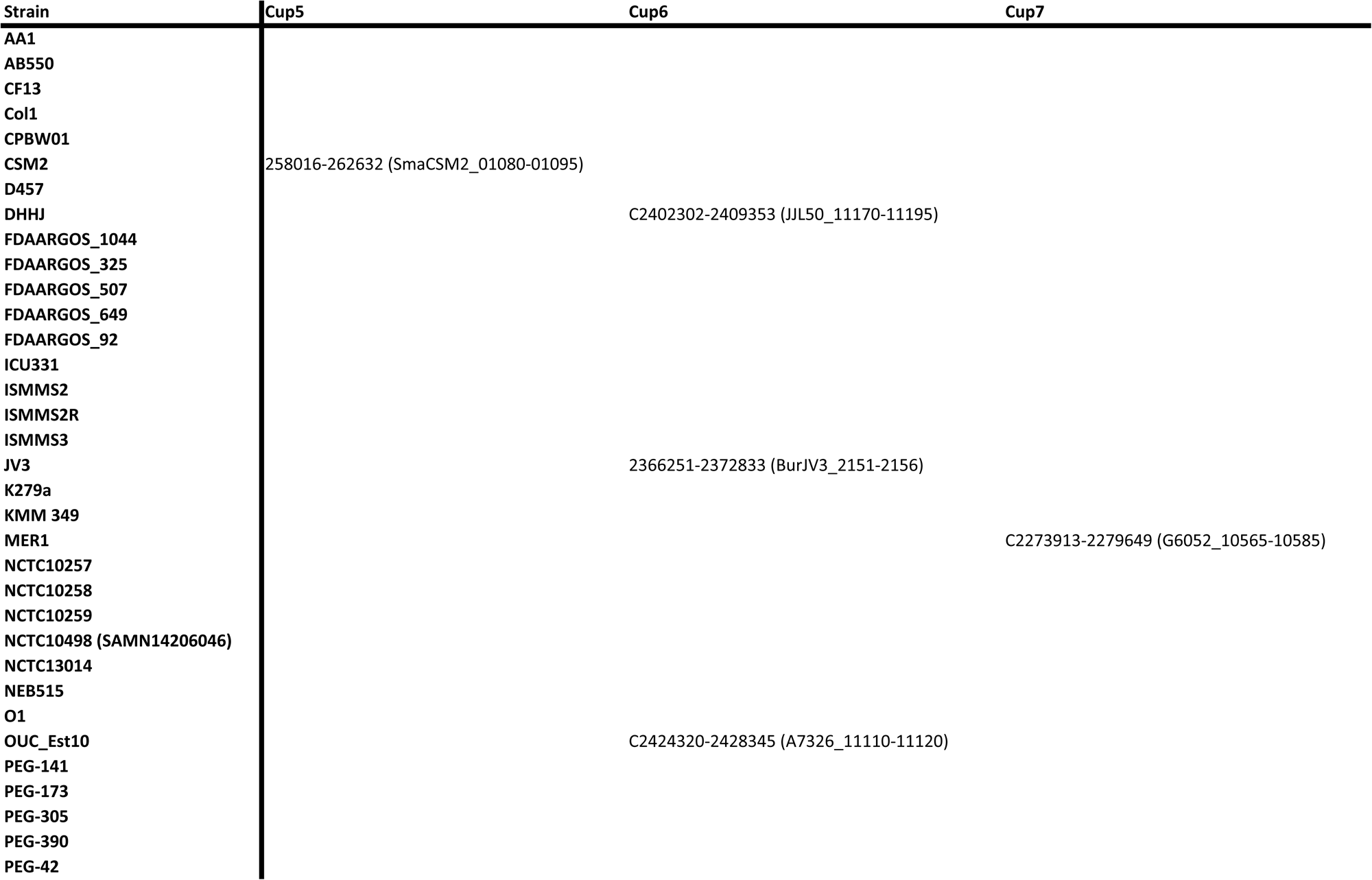
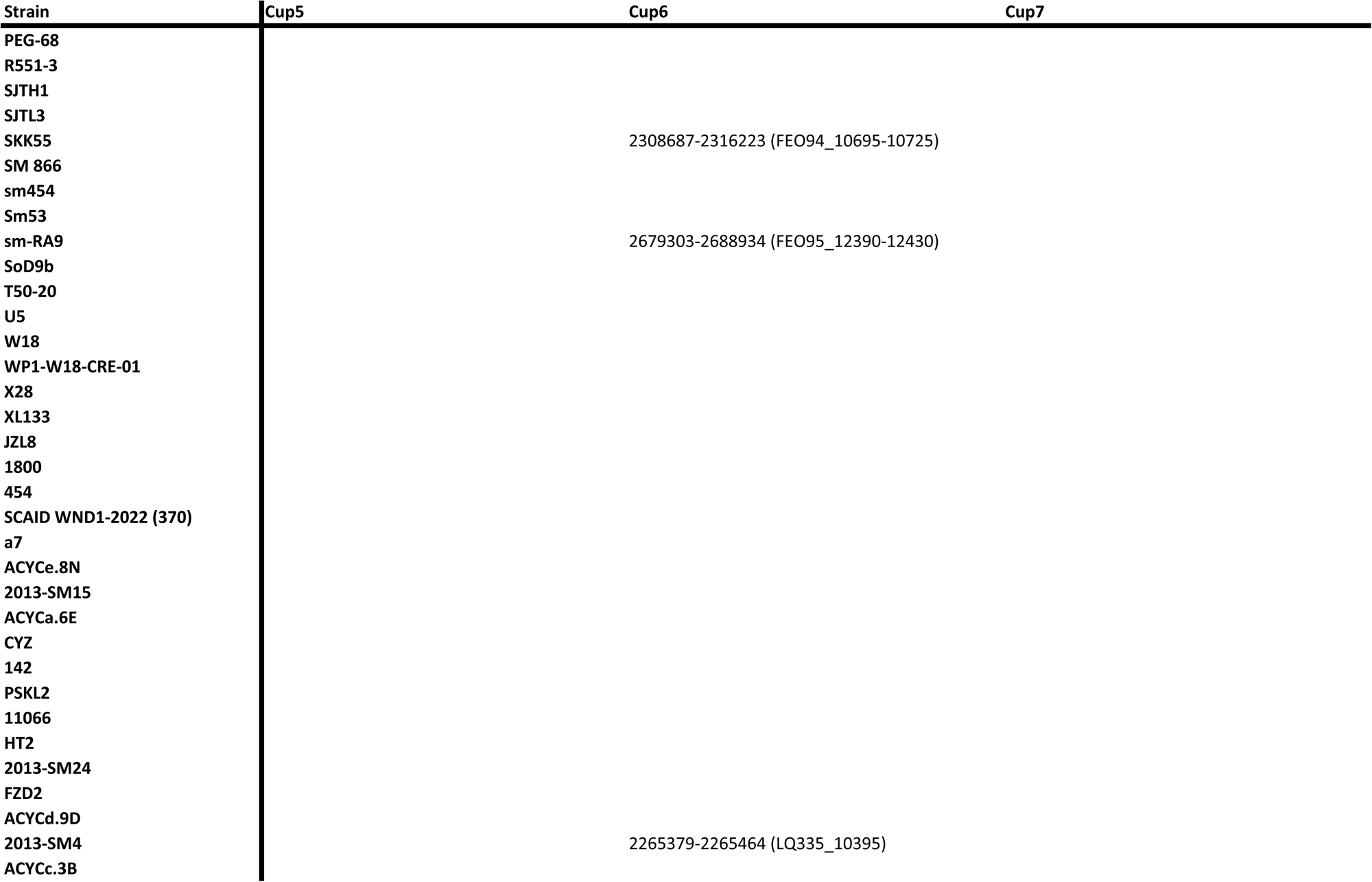
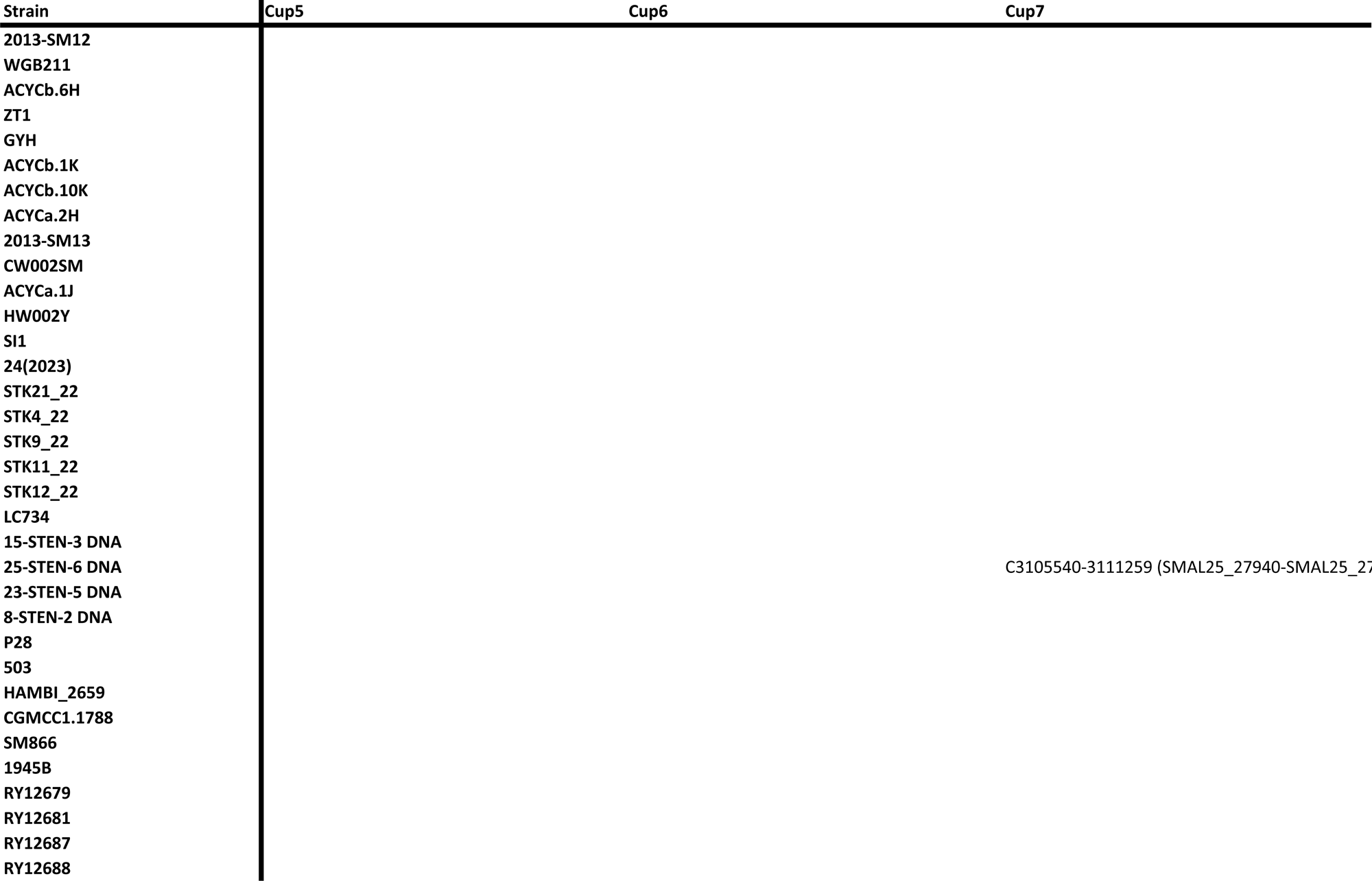
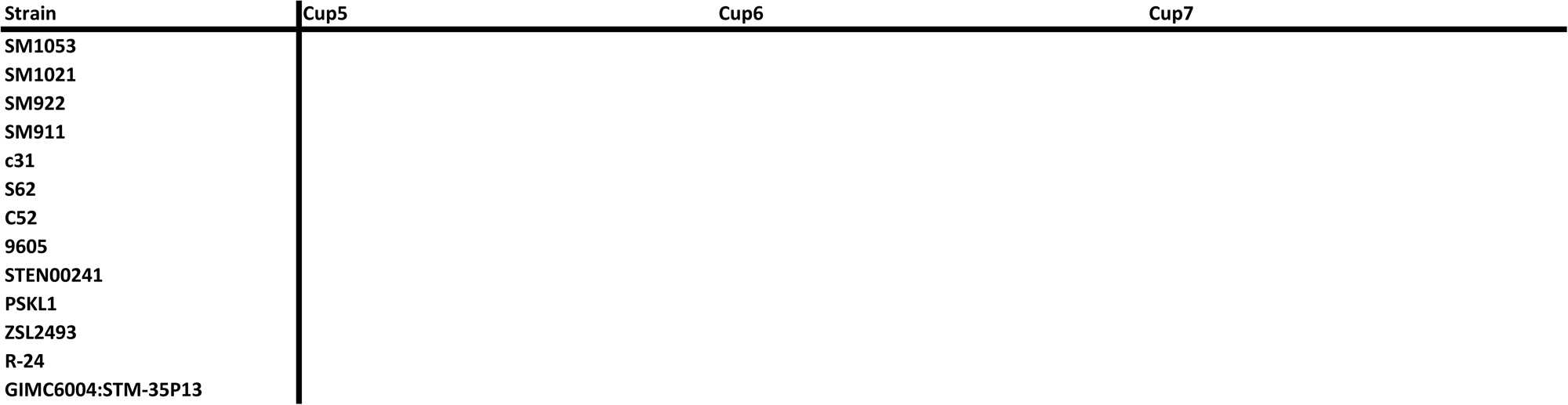
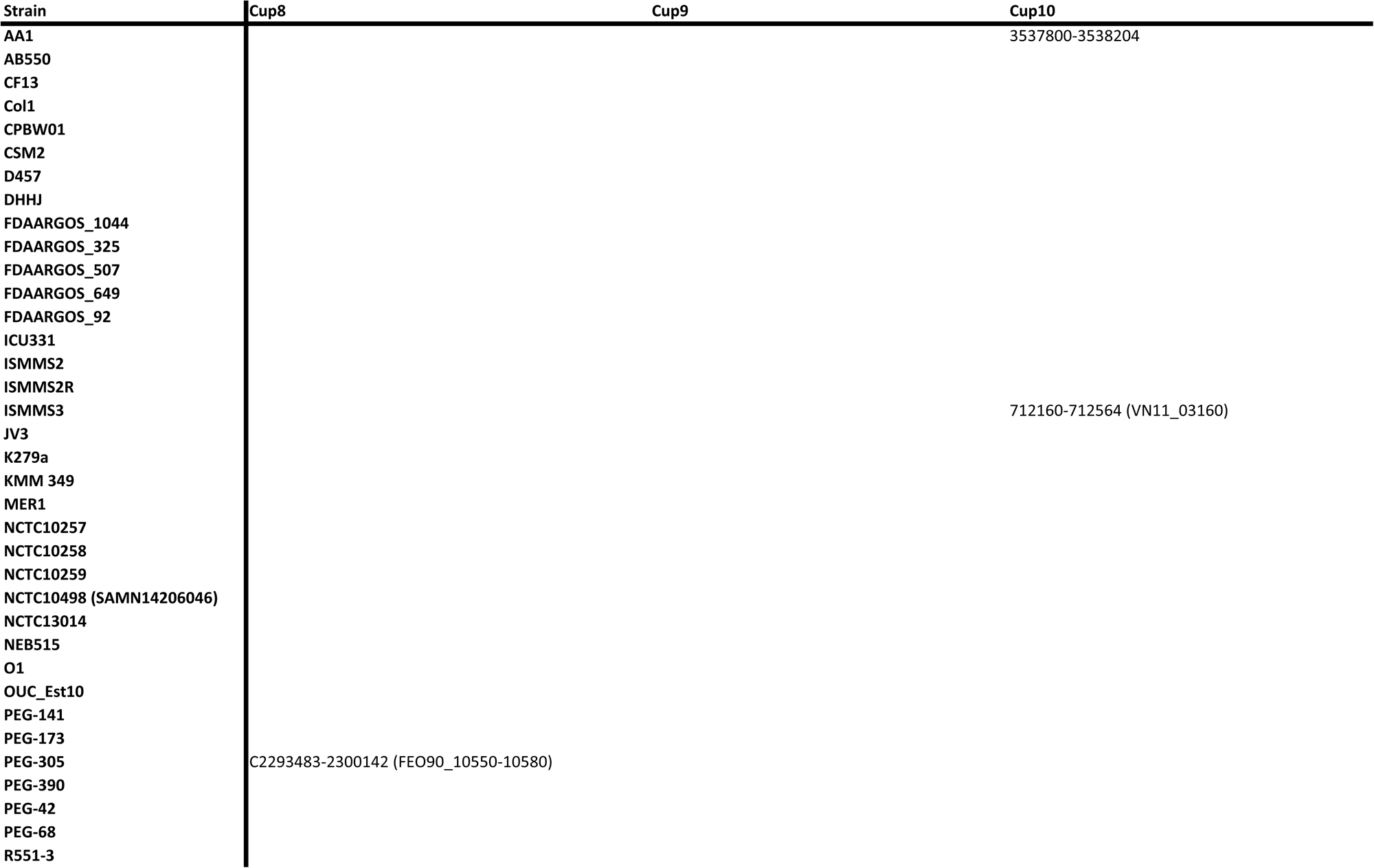
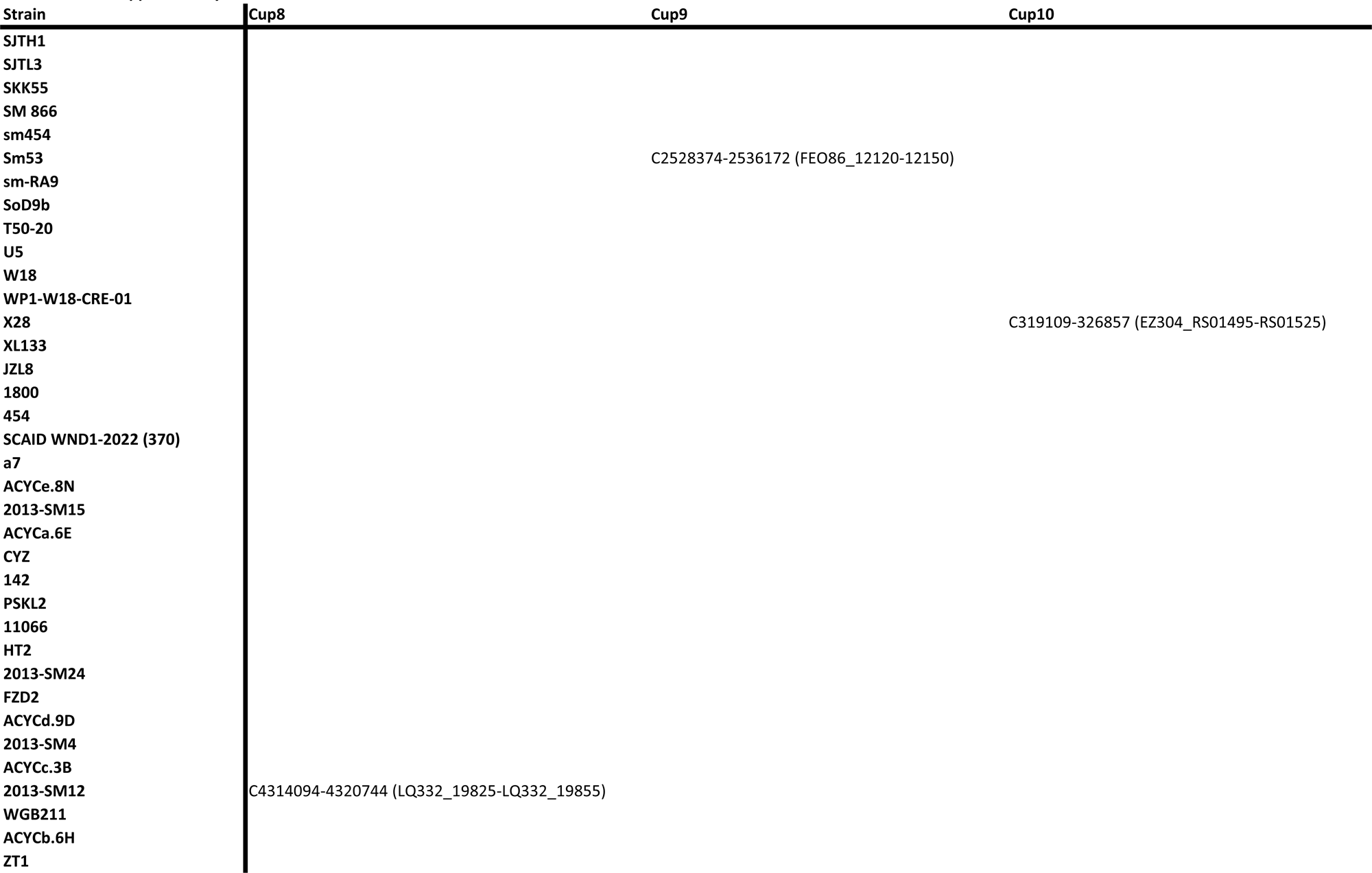
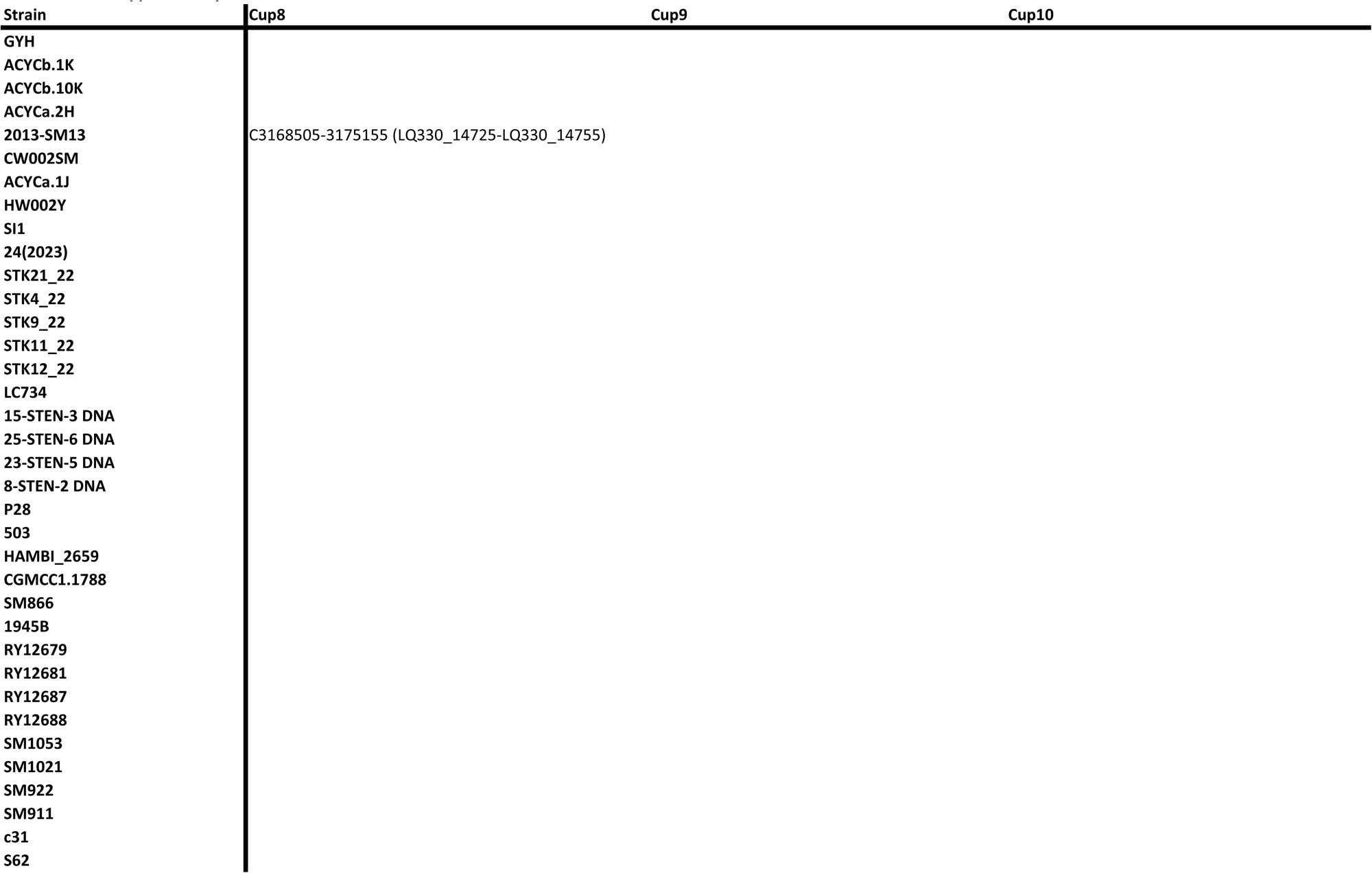
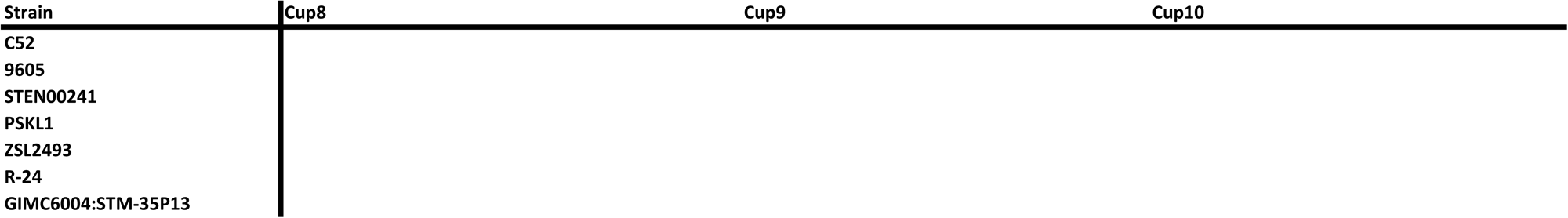
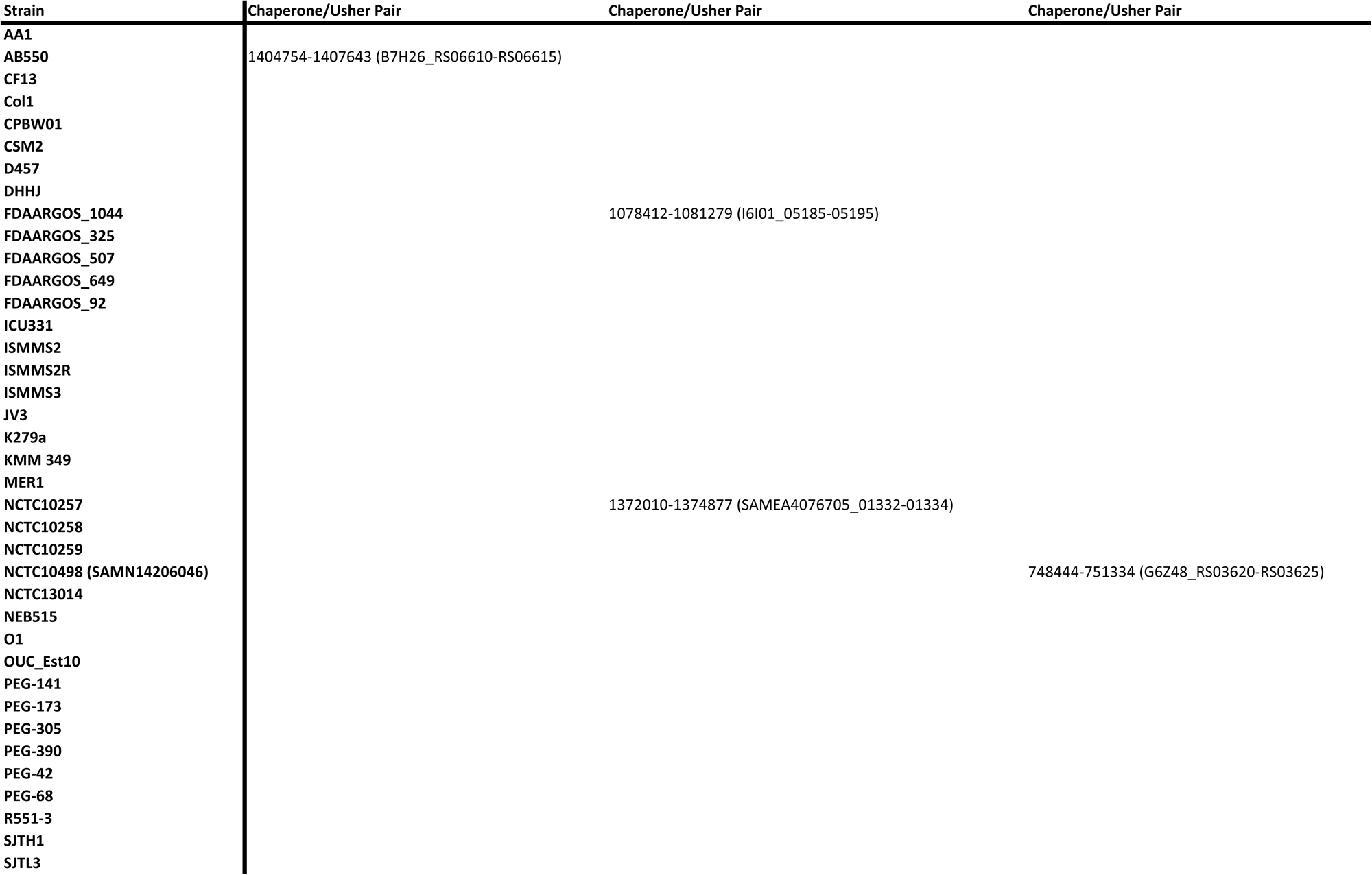
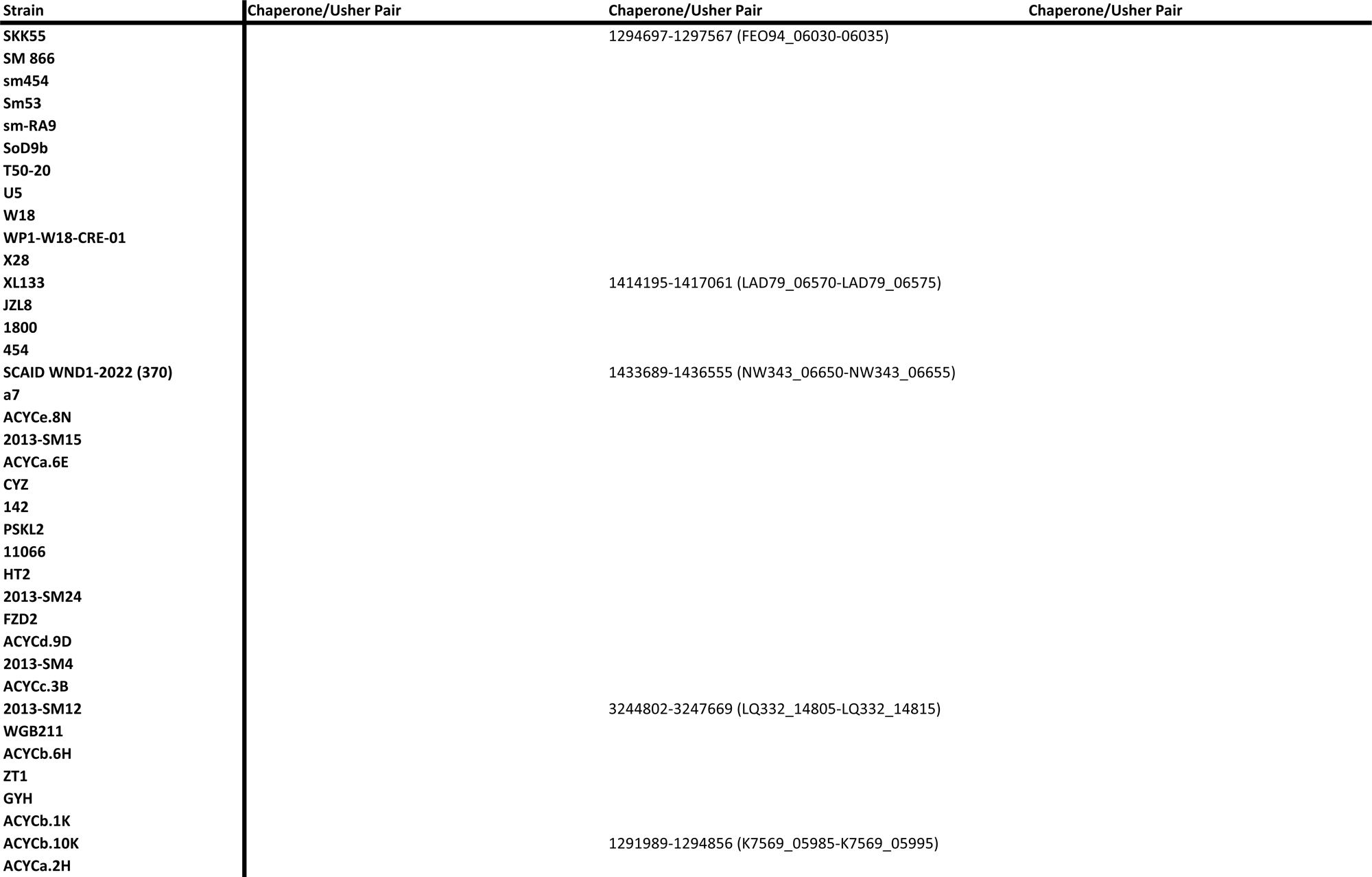
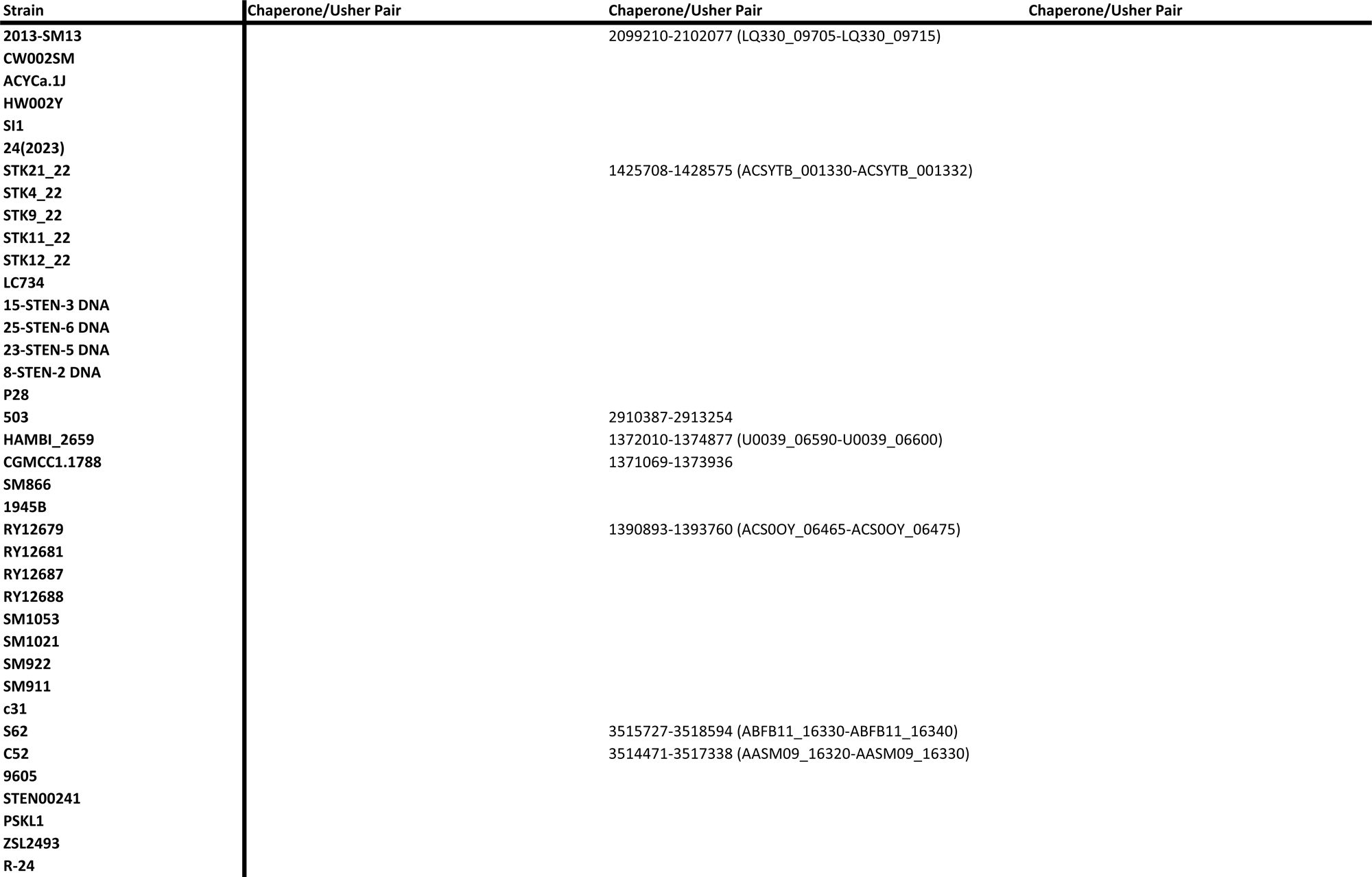
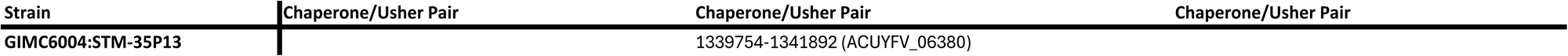
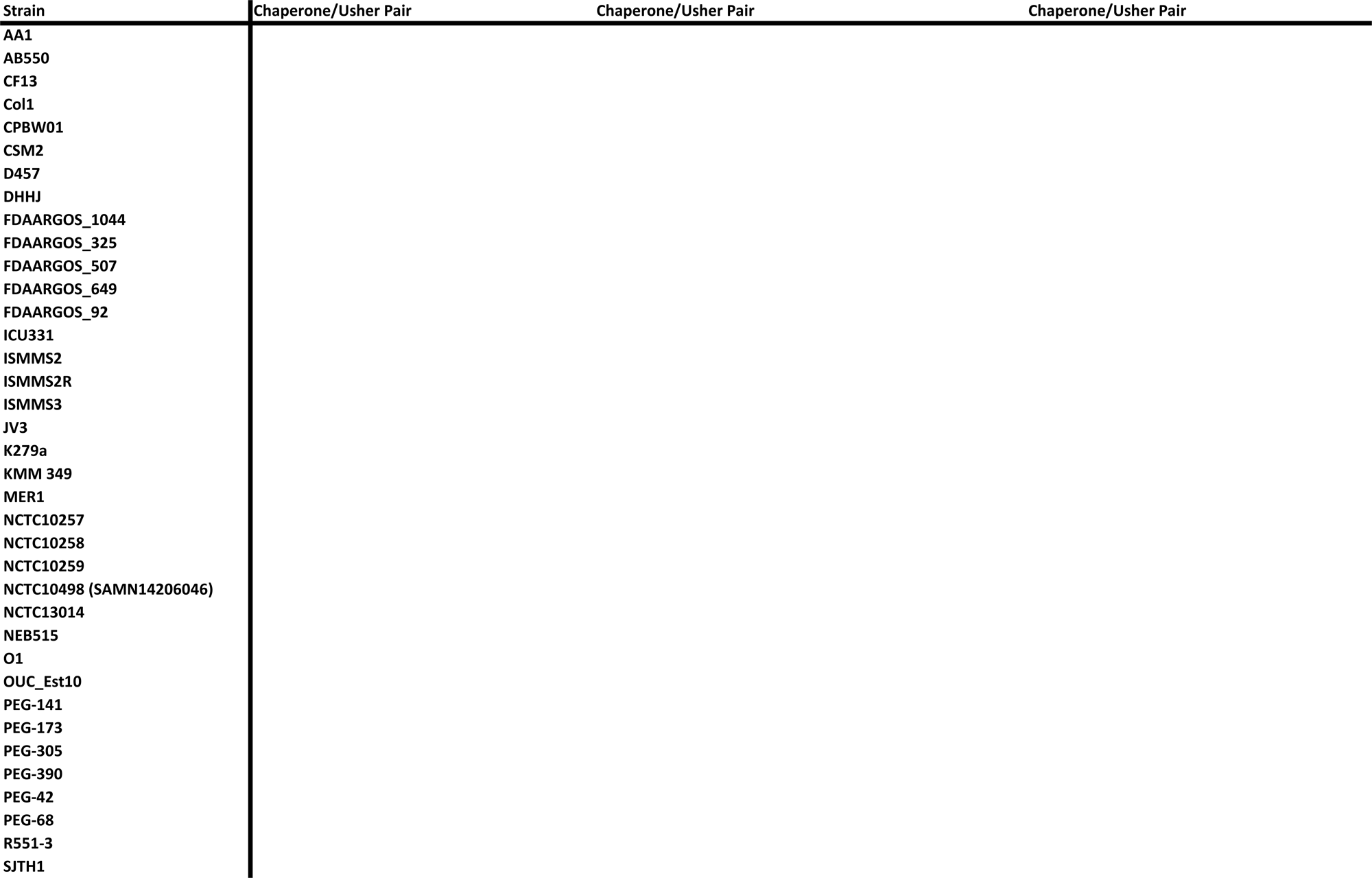
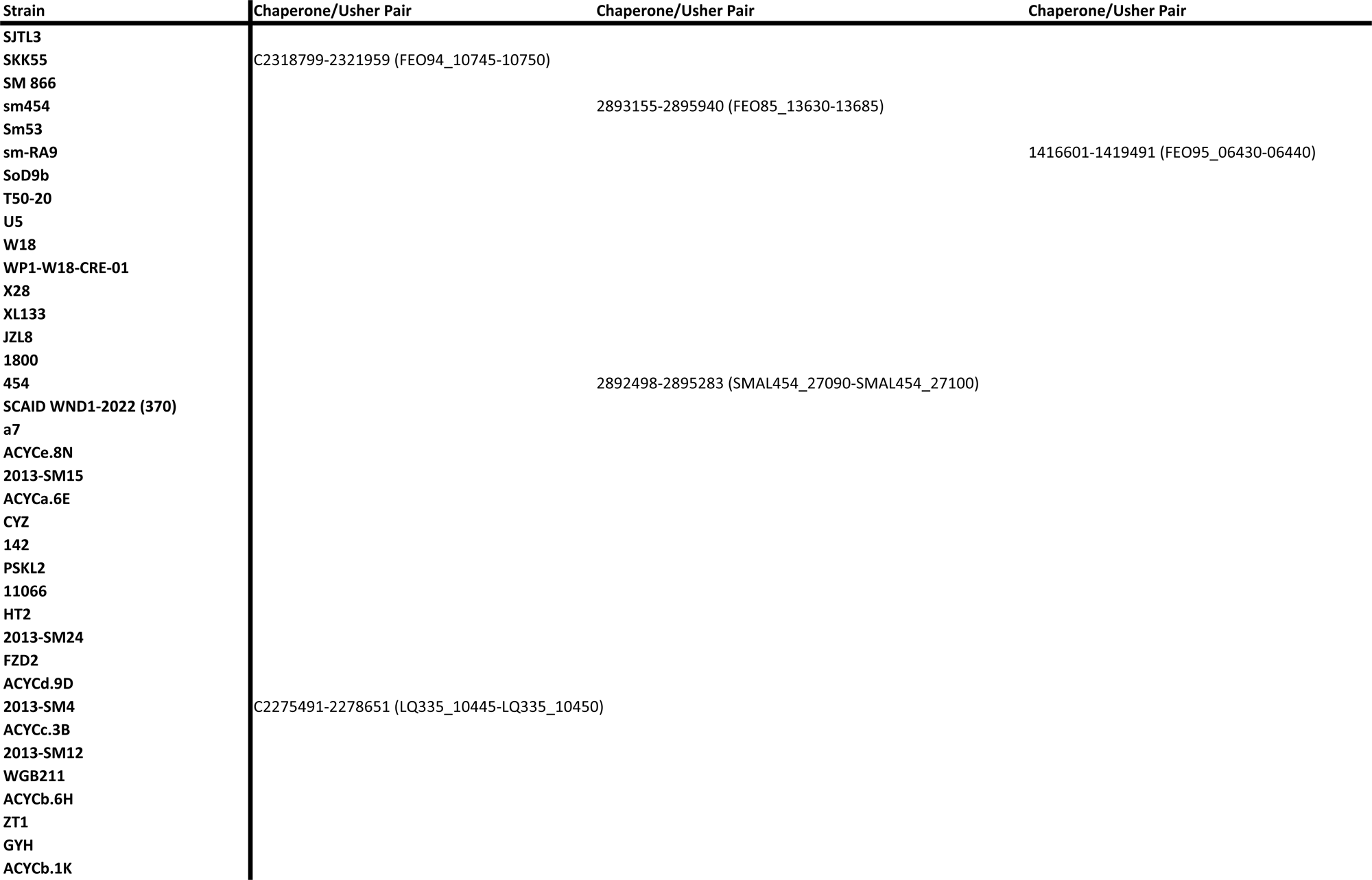
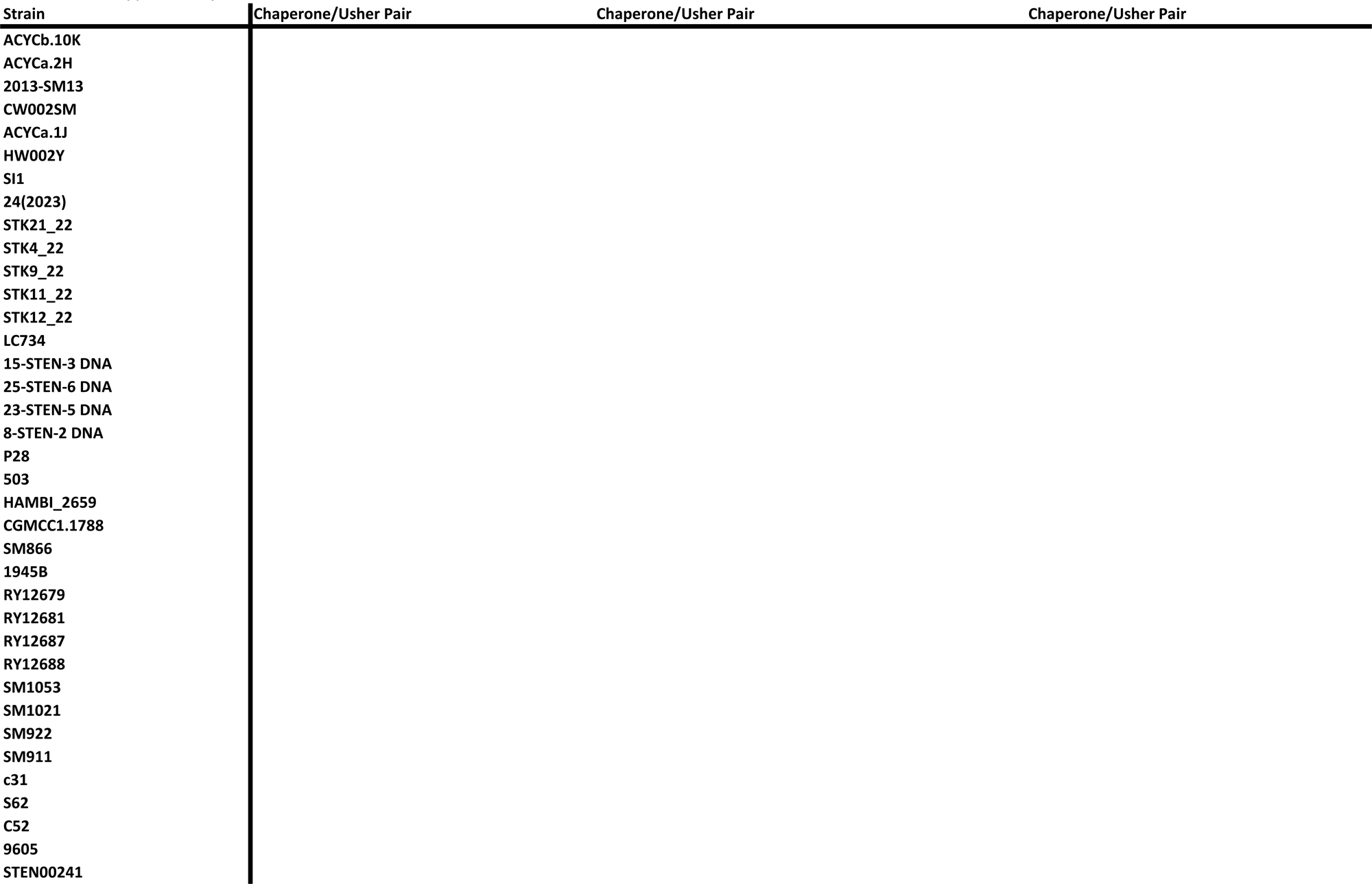
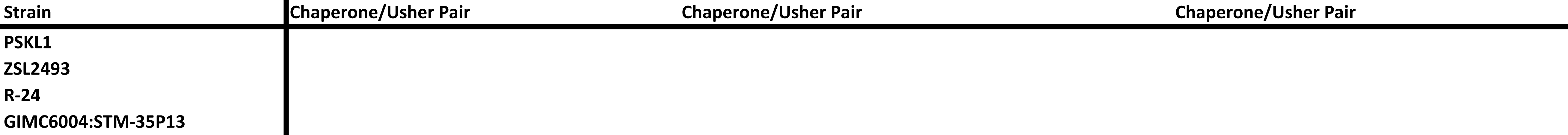
Distribution of additional chaperone–usher pilus loci in *Stenotrophomonas maltophilia*. Genomic loci corresponding to additional predicted chaperone–usher pilus operons identified across fully sequenced *Stenotrophomonas maltophilia* strains from the NCBI database. Numbers listed indicate the genetic locus positions within each strain genome. A “C” preceding the locus number range denotes that the locus is located on the complementary strand of the genome. Gene designations shown in parentheses represent the annotated genetic nomenclature of each locus in the NCBI genome database. An asterisk (*) indicates loci in which only the usher gene shares homology with the Scs operon, whereas a double asterisk (**) indicates loci in which the usher and the first two genes share homology with the Scs operon.

**Supplementary Table 3.** Distribution of additional pilus-associated and adhesive loci in *Stenotrophomonas maltophilia*. Genomic loci corresponding to additional pilus-associated systems identified across fully sequenced *Stenotrophomonas maltophilia* strains, including type IV pili, Tad pili, type II pilus assembly components, and afimbrial adhesins identified through NCBI genome annotations. Numbers listed indicate the genetic locus positions within each strain genome. A “C” preceding the locus number range denotes that the locus is located on the complementary strand of the genome. Gene designations shown in parentheses represent the annotated genetic nomenclature of each locus in the NCBI genome database.

Bhaumik et al. Supplementary Table 3
| Strain | afaD | afaD2 | fimbrial protein |
| --- | --- | --- | --- |
| AA1 |  |  |  |
| AB550 | C4584100-4584549 (B7H26_21590) |  |  |
| CF13 | C4304412-4304861 (HKK60_19845) |  |  |
| Col1 |  |  |  |
| CPBW01 |  |  |  |
| CSM2 | C4451541-4451990 (SmaCSM2_20530) |  |  |
| D457 | C4476199-4476648 |  |  |
| DHHJ |  |  |  |
| FDAARGOS_1044 | C4396881-4397330 (I6I01_20665) |  |  |
| FDAARGOS_325 |  | 3404584-3405045 (CEQ03_15825-CEQ03_15830) |  |
| FDAARGOS_507 | C3986406-3986855 (EGY09_18895) |  |  |
| FDAARGOS_649 | C976598-977047 (FOB57_04415) |  |  |
| FDAARGOS_92 | C3528362-3528811 (AL480_16850) |  |  |
| ICU331 | C4673869-4674318 (FEO84_22035) |  |  |
| ISMMS2 |  | C4222846-4223297 (YH67_19165-YH67_19170) |  |
| ISMMS2R |  | C4222846-4223297 (YH68_19165-YH68_19170) |  |
| ISMMS3 |  |  | 3306013-3306435 (VN11_15225) |
| JV3 | C4255139-4255588 (BurJV3_3851) |  |  |
| K279a | C4530368-4530817 (Smlt4424-Smlt4423) |  |  |
| KMM 349 |  |  |  |
| MER1 |  |  |  |
| NCTC10257 | C4690475-4690924 (SAMEA4076705_04430) |  |  |
| NCTC10258 |  |  |  |
| NCTC10259 | C1519774-1520223 (NCTC10259_01393) |  |  |
| NCTC10498 (SAMN14206046) | C4008717-4009166 |  |  |
| NCTC13014 |  |  |  |
| NEB515 | 684170-684619 (HGN30_02980) |  |  |
| O1 | C3429704-3430153 |  |  |
| OUC_Est10 |  |  |  |
| PEG-141 |  |  |  |
| PEG-173 |  |  |  |
| PEG-305 |  |  |  |
| PEG-390 |  | C4292901-4293362 (FEO91_19945-FEO91_19950) |  |
| PEG-42 |  | C4566799-4567260 (FEO92_21535-FEO92_21540) |  |
| PEG-68 | C4310895-4311344 (FEO93_20105) |  |  |
| R551-3 |  |  |  |
| SJTH1 |  |  |  |
| SJTL3 |  |  |  |
| SKK55 | C4391275-4391724 (FEO94_20380) |  |  |
| SM 866 |  |  |  |

Bhaumik et al. Supplementary Table 3
| Strain | afaD | afaD2 | fimbrial protein |
| --- | --- | --- | --- |
| sm454 | C4377113-4377562 (FEO85_20425) |  |  |
| Sm53 |  |  |  |
| sm-RA9 |  | C4695574-4696035 (FEO95_21575-FEO95_21580) |  |
| SoD9b |  | C4133911-4134372 (G4G30_18980-G4G30_18985) |  |
| T50-20 | C4451091-4451540 |  |  |
| U5 |  | C4261443-4261904 (FEO87_19620-FEO87_19625) |  |
| W18 |  | C4453422-4453883 (C9J74_RS20845) |  |
| WP1-W18-CRE-01 | C4598568-4599017 (WP1W18C01_41720) |  |  |
| X28 |  |  |  |
| XL133 | C4372804-4373253 (LAD79_20225-LAD79_20230) |  |  |
| JZL8 |  | 4351719-4352180 |  |
| 1800 |  |  |  |
| 454 | C4358111-4358560 (SMAL454_40380-SMAL454_40390) |  |  |
| SCAID WND1-2022 (370) | C4525149-4525598 (NW343_20970-NW343_20975) |  |  |
| a7 |  |  |  |
| ACYCe.8N | C4192681-4193130 (K7573_19295-K7573_19300) |  |  |
| 2013-SM15 |  |  |  |
| ACYCa.6E | C4213599-4214048 (K7574_19550-K7574_19555) |  |  |
| CYZ | 246220-246669 |  |  |
| 142 | C4517271-4517720 (NDK23_21025) |  |  |
| PSKL2 | C4281165-4281614 (L6173_19725-L6173_19730) |  |  |
| 11066 |  | C4199366-4199827 (RBI20_19430-RBI20_19435) |  |
| HT2 |  | C4230113-4230575 (PWE35_19620-PWE35_19625) |  |
| 2013-SM24 | C3960765-3961214 (LQ331_18340-LQ331_18345) |  |  |
| FZD2 |  |  |  |
| ACYCd.9D |  |  |  |
| 2013-SM4 | C4350499-4350948 (LQ335_20110-LQ335_20115) |  |  |
| ACYCc.3B | C4125087-4125536 (K7564_19230) |  |  |
| 2013-SM12 |  |  |  |
| WGB211 |  | 4624507-4624968 |  |
| ACYCb.6H |  |  |  |
| ZT1 | 571163-571612 (J3U96_02515-J3U96_02520) |  |  |
| GYH |  |  |  |
| ACYCb.1K |  | C4135580-4136041 (K7567_19060-K7567_19065) |  |
| ACYCb.10K |  | C4364325-4364786 (K7569_20215-K7569_20220) |  |
| ACYCa.2H |  |  |  |
| 2013-SM13 |  |  |  |
| CW002SM | 259533-259982 (N0074_01080-N0074_01085) |  |  |
| ACYCa.1J | C4373674-4374123 (K7566_20260-K7566_20265) |  |  |
| HW002Y |  |  |  |

Bhaumik et al. Supplementary Table 3
| Strain | afaD | afaD2 | fimbrial protein |
| --- | --- | --- | --- |
| SI1 | C4663220-4663669 (RS400_21795) |  |  |
| 24(2023) |  |  |  |
| STK21_22 | C4518232-4518681 (ACSYTB_004193-ACSYTB_004194) |  |  |
| STK4_22 |  |  |  |
| STK9_22 |  |  |  |
| STK11_22 |  |  |  |
| STK12_22 |  |  |  |
| LC734 | C4510463-4510912 (ACT4MX_21140-ACT4MX_21145) |  |  |
| 15-STEN-3 DNA | C4483936-4484385 (SMAL15_41600-SMAL15_41610) |  |  |
| 25-STEN-6 DNA |  |  |  |
| 23-STEN-5 DNA | C4018296-4018745 (SMAL23_37530-SMAL23_37540) |  |  |
| 8-STEN-2 DNA | C4046032-4046481 (SMAL8_37980-SMAL8_37990) |  |  |
| P28 | C4739829-4740278 (QVH29_22215) |  |  |
| 503 | 4548153-4548602 |  |  |
| HAMBI_2659 | C4690477-4690926 (U0039_22060-U0039_22065) |  |  |
| CGMCC1.1788 | 4688423-4688872 |  |  |
| SM866 |  |  |  |
| 1945B | C3782755-3783204 (ABK042_17605-ABK042_17610) |  |  |
| RY12679 |  | C4623208-4623669 (ACS00Y_21485-ACS00Y_21490) |  |
| RY12681 |  |  |  |
| RY12687 |  |  |  |
| RY12688 | C4034233-4034682 (ACS00T_18720-ACS00T_18725) |  |  |
| SM1053 | 3856878-3857327 |  |  |
| SM1021 | 4279591-4280040 |  |  |
| SM922 | 252006-252455 |  |  |
| SM911 | 252005-252454 |  |  |
| c31 | C4242836-4243285 (KNV95_19725-KNV95_19730) |  |  |
| S62 |  |  |  |
| C52 |  |  |  |
| 9605 |  | C957672-958132 (ABK044_04290-ABK044_04295) |  |
| STEN00241 |  |  |  |
| PSKL1 |  |  |  |
| ZSL2493 |  |  |  |
| R-24 |  |  |  |
| GIMC6004:STM-35P13 |  |  |  |

Bhaumik et al. Supplementary Table 3
| Strain | fimbrial protein (pilE1_2)(pilE1_1)(pilE_3) | putative fimbrial biogenesis protein (pilW)(pilF) | putative pilin subunit |
| --- | --- | --- | --- |
| AA1 | 3626905-3627639 (BTU49_RS16485) | 1476409-1477022 |  |
| AB550 | 4306934-4307698 (B7H26_20245) | 1939023-1939808 (B7H26_09190) | 2732217-2732404 (B7H26_12825) |
| CF13 | 4070857-4071615 (HKK60_18725) | C2759015-2759800 (HKK60_12550) | 2024063-2024269 (HKK60_09255) |
| Col1 | 3914127-3914885 (KS456_RS18015) | 1841151-1841936 | 2521753-2521928 |
| CPBW01 | 3915111-3915869 (GS396_17770) | 1808771-1809556 (GS396_08415) | 2544754-2544929 (GS396_11595) |
| CSM2 | 4156103-4156867 (SmaCSM2_19075) | 1939201-1939986 (SmaCSM2_09050) | 2759019-2759205 (SmaCSM2_12710) |
| D457 | 4248729-4249493 (SMD_3781) | 2046392-2047135 (SMD_1851) | 2796472-2796659 (SMD_2505) |
| DHHJ | 4151803-4152561 (JIL50_19195) | 1806168-1806953 (JIL50_08375) | 2643494-2643669 (JIL50_12220) |
| FDAARGOS_1044 | 4107902-4108660 (I6I01_19260) | 1896334-1897119 (I6I01_09205) | 2682723-2682919 (I6I01_12705) |
| FDAARGOS_325 | C3672245-3672997 (CEQ03_17135) | C993436-994221 (CEQ03_04560) | C265087-265293 (CEQ03_01285) |
| FDAARGOS_507 | 3755356-3756114 (EGY09_17775) | 1590282-1591067 (EGY09_07705) | 2304620-2304826 (EGY09_10910) |
| FDAARGOS_649 | 741539-742297 (FOB57_03275) | 3239344-3240129 (FOB57_15050) | 4029587-4029793 (FOB57_18670) |
| FDAARGOS_92 | 3226343-3227113 (AL480_15410) | 1071580-1072365 (AL480_05535) | 1837863-1838069 (AL480_09020) |
| ICU331 | 4358054-4358812 (FEO84_20520) | C2992674-2993459 (FEO84_14200) | 2155637-2155843 (FEO84_10400) |
| ISMMS2 | 3960586-3961344 (YH67_17955) | 1875792-1876577 (YH67_08665) | 2615117-2615286 (YH67_11905) |
| ISMMS2R | 3960586-3961344 (YH68_17955) | 1875792-1876577 (YH68_08665) | 2615117-2615286 (YH68_11905) |
| ISMMS3 | 4239989-4240741 (VN11_19320) | 1968372-1969157 (VN11_09250) |  |
| JV3 | 4023067-4023831 (BurJV3_3632) | 1869353-1870138 (BurJV3_1704) | 2618358-2618544 (BurJV3_2374) |
| K279a | 4297473-4298231 (Smlt4182) | 2082020-2082805 (Smlt2055-Smlt2056) | 2909938-2910144 (Smlt2867) |
| KMM 349 | 4039058-4039816 (KMM349_35930) | 1910218-1911003 (KMM349_17080) |  |
| MER1 |  | 1866180-1866965 (G6052_08755) |  |
| NCTC10257 | 4401496-4402254 (SAMEA4076705_04156) | 2189930-2190715 (SAMEA4076705_02137) |  |
| NCTC10258 | 3940881-3941639 (NCTC10258_03635) | 1847279-1848064 (NCTC10258_01734) |  |
| NCTC10259 | 1296303-1297067 (NCTC10259_01171) | 3662110-3662895 (NCTC10259_03407) |  |
| NCTC10498 (SAMN14206046) | 3767255-3768013 (G6Z48_RS17935) | 1437516-1438296 | 2267687-2267883 |
| NCTC13014 | C4090291-4091043 (NCTC13014_03803) | C1597958-1598743 (NCTC13014_01452) |  |
| NEB515 | C942358-943116 (HGN30_04200) | C3261284-3262069 (HGN30_14945) | C2343672-2343878 (HGN30_10650) |
| O1 | 3197750-3198508 | 1097895-1098682 | 1841412-1841618 |
| OUC_Est10 | 4106843-4107595 (A7326_18700) | 1894529-1895316 (A7326_08745) | 2658141-2658313 (A7326_12105) |
| PEG-141 | 4435916-4436674 (FEO88_20795) | 2079049-2079834 (FEO88_09915) | 2967606-2967781 (FEO88_13975) |
| PEG-173 | 4240599-4241357 (FEO89_19450) | 1983773-1984558 (FEO89_09335) | 2782014-2782189 (FEO89_12940) |
| PEG-305 | 3954162-3954914 (FEO90_18080) | 1925490-1926275 (FEO90_08930) | 2622194-2622380 (FEO90_12060) |
| PEG-390 | 4050663-4051415 (FEO91_18780) | 1922092-1922877 (FEO91_09165) | 2680633-2680802 (FEO91_12570) |
| PEG-42 | 4340152-4340904 (FEO92_20440) | 2009618-2010403 (FEO92_09625) |  |
| PEG-68 | 4080283-4081047 (FEO93_18965) | 1838553-1839338 (FEO93_08710) | 2644588-2644775 (FEO93_12445) |
| R551-3 | 4042995-4043753 (Sma1_3589) | 1868386-1869171 (Sma1_1658) |  |
| SJTH1 | 4362570-4363322 (C6Y55_20345) | 1947125-1947910 (C6Y55_09145) |  |
| SJTL3 | 4291381-4292139 (DM611_19905) | 1913124-1913909 (DM611_09155) | 2803920-2804089 (DM611_13245) |
| SKK55 | 4164455-4165213 (FEO94_19295) | 1870802-1871587 (FEO94_08775) |  |
| SM 866 | 4521459-4522217 (DUW70_21530) | C3099977-3100762 (DUW70_14680) | 2249508-2249683 (DUW70_10785) |
| sm454 | 4116237-4117001 (FEO85_19205) | 1959007-1959792 (FEO85_09370) | 2720803-2720999 (FEO85_12825) |

Bhaumik et al. Supplementary Table 3
| Strain | fimbrial protein (pilE1_2)(pilE1_1)(pilE_3) | putative fimbrial biogenesis protein (pilW)(pilF) | putative pilin subunit |
| --- | --- | --- | --- |
| **Sm53** | 4140735-4141487 (FEO86_19350) | 1994020-1994805 (FEO86_09610) | 2783240-2783436 (FEO86_13205) |
| **sm-RA9** | 4439057-4439752 (FEO95_20395) | 2155459-2156244 (FEO95_10085) |  |
| **SoD9b** | 3903518-3904281 (G4G30_17880) | 1895938-1896723 (G4G30_08800) | 2574790-2574977 (G4G30_11860) |
| **T50-20** | 4201139-4201897 (FOQ13_RS19290) | 1927737-1928522 (not annotated) |  |
| **U5** |  | 1930816-1931601 (FEO87_09045) |  |
| **W18** | 4230241-4230993 (C9J74_RS19765) | 1956590-1957366 |  |
| **WP1-W18-CRE-01** | 4361771-4362535 (WP1W18C01_39480) | 2078322-2079107 (WP1W18C01_19260) | 2837502-2837708 (WP1W18C01_25830) |
| **X28** | 1929899-1930657 (EZ304_RS08755) | 3956649-3956708 | 568649-568849 |
| **XL133** | 4126213-4126971 (LAD79_19030-LAD79_19035) | 2020010-2020795 (LAD79_09460-LAD79_09465) | 2746688-2746894 (LAD79_12740) |
| **JZL8** | 4040924-4041682 | 1903760-1904545 | 2659213-2659382 |
| **1800** | 4309631-4310389 (STEN1800_4195) | 2039133-2039918 (STEN1800_2026-STEN1800_2027) |  |
| **454** | 4115724-4116488 (SMAL454_38080) | 1958355-1959140 (SMAL454_18550-SMAL454_18560) | 2720148-2720344 (SMAL454_25420) |
| **SCAID WND1-2022 (370)** | 4294110-4294868 (NW343_19850-NW343_19855) | 2035111-2035896 (NW343_09530-NW343_09535) | 2869425-2869631 (NW343_13330) |
| **a7** | 3940299-3941063 | 1872110-1872895 | 2609255-2609451 |
| **ACYCe.8N** | 3957066-3957824 (K7573_18155) | 1853589-1854374 (K7573_08625-K7573_08630) | 2602991-2603197 (K7573_12005) |
| **2013-SM15** | C2652839-2653597 (LQ329_12190) | C189921-190706 (LQ329_00865-LQ329_00870) | 4085691-4085897 (LQ329_18725) |
| **ACYCa.6E** | 3983356-3984114 (K7574_18435) | 1869227-1870012 (K7574_08735-K7574_08740) | 2595354-2595560 (K7574_12055) |
| **CYZ** | 477223-477981 | 2591476-2592261 | 1856501-1856707 |
| **142** | 4284781-4285539 (NDK23_19905) | 1940148-1940933 (NDK23_09165-NDK23_09170) | 2817287-2817493 (NDK23_13190) |
| **PSKL2** | 3997307-3998065 (L6173_18390) | 1875218-1876003 (L6173_08700-L6173_08705) | 2619802-2620008 (L6173_12030) |
| **11066** | 3965476-3966240 (RBI20_18295) | 1888811-1889596 (RBI20_08890-RBI20_08895) | 2630611-2630807 (RBI20_12265) |
| **HT2** | 3996447-3997211 (PWE35_18485) | 1886052-1886837 (PWE35_08780-PWE35_08785) | 2663570-2663776 (PWE35_12455) |
| **2013-SM24** | 3728914-3729672 (LQ331_17220) | 1647556-1648341 (LQ331_07830-LQ331_07835) | 2386504-2386710 (LQ331_11190) |
| **FZD2** | 4221407-4222159 (K1516_19500) | 1903072-1903857 (K1516_08940-K1516_08945) |  |
| **ACYCd.9D** |  | 1836768-1837553 (K7563_08595-K7563_08600) | 2623204-2623391 (K7563_12135) |
| **2013-SM4** | 4123849-4124607 (LQ335_19015) | 1890007-1890792 (LQ335_08725-LQ335_08730) | 2730476-2730663 (LQ335_12580) |
| **ACYCc.3B** |  |  | 2526697-2526884 (K7564_11970) |
| **2013-SM12** | 1355736-1356488 (LQ332_06080) | 3859181-3859966 (LQ332_17735-LQ332_17740) | 40163-40349 (LQ332_00165) |
| **WGB211** | 4341763-4342515 | 1951909-1952694 |  |
| **ACYCb.6H** | 3858487-3859239 (K7565_17750) | 1815859-1816644 (K7565_08460-K7565_08465) | 2487150-2487319 (K7565_11460) |
| **ZT1** | C790827-791579 (J3U96_03570) | C2886974-2887759 (J3U96_13055-J3U96_13060) | 2191705-2191880 (J3U96_09940) |
| **GYH** |  |  |  |
| **ACYCb.1K** | 3909602-3910354 (K7567_17995) | 1829199-1829984 (K7567_08540-K7567_08545) | 2525970-2526139 (K7567_11675) |
| **ACYCb.10K** | 4135796-4136548 (K7569_19135) |  | 2746656-2746831 (K7569_12940) |
| **ACYCa.2H** | 4032971-4033729 (K7568_18630) | 1913763-1914548 (K7568_08920-K7568_08925) | 2640346-2640552 (K7568_12205) |
| **2013-SM13** | 210584-211336 (LQ330_00985) | 2713589-2714374 (LQ330_12635-LQ330_12640) | 3507840-3508026 (LQ330_16240) |
| **CW002SM** | C587458-588222 (N0074_02595-N0074_02600) | C2805382-2806167 (N0074_12840-N0074_12845) | C2055058-2055254 (N0074_09405) |
| **ACYCa.1J** | 4135072-4135830 (K7566_19120-K7566_19125) | 1957856-1958641 (K7566_09250-K7566_09255) | 2787848-2788054 (K7566_12995) |
| **HW002Y** | 3603657-3604415 (NDR96_16590-NDR96_16595) | 1530440-1531225 (NDR96_07160-NDR96_07165) | 2233254-2233423 (NDR96_10360) |
| **SI1** | 4422378-4423136 (RS400_20645) | 1953722-1954507 (RS400_09175-RS400_09180) |  |
| **24(2023)** |  |  |  |

Bhaumik et al. Supplementary Table 3
| Strain | fimbrial protein (pilE1_2)(pilE1_1)(pilE_3) | putative fimbrial biogenesis protein (pilW)(pilF) | putative pilin subunit |
| --- | --- | --- | --- |
| STK21_22 | 4288222-4288980 (ACSYTB_003971) | 2028443-2029228 (ACSYTB_001908-ACSYTB_001909) | 2856854-2857060 (ACSYTB_002660) |
| STK4_22 | 4439992-4440756 (ACSYS1_004230) | 2059449-2060234 (ACSYS1_001979-ACSYS1_001980) | 2760900-2761106 (ACSYS1_002617) |
| STK9_22 | 4457114-4457878 (ACSYS3_004222) | 2071523-2072308 (ACSYS3_001967-ACSYS3_001968) | 2773080-2773286 (ACSYS3_002605) |
| STK11_22 | 4440172-4440936 (ACSYS4_004226) | 2059458-2060243 (ACSYS4_001976-ACSYS4_001977) | 2761018-2761224 (ACSYS4_002614) |
| STK12_22 | 4440099-4440863 (ACSYS5_004232) | 2059450-2060235 (ACSYS5_001980-ACSYS5_001981) | 2760911-2761117 (ACSYS5_002618) |
| LC734 | 4280453-4281211 (ACT4MX_20030) | 2024393-2025178 (ACT4MX_09625-ACT4MX_09630) | 2844791-2844997 (ACT4MX_13330) |
| 15-STEN-3 DNA | 4236765-4237523 (SMAL15_39180) | 2048669-2049454 (SMAL15_19260-SMAL15_19270) | 2866615-2866811 (SMAL15_26840) |
| 25-STEN-6 DNA | C463981-464739 (SMAL25_04150) |  |  |
| 23-STEN-5 DNA | 3784201-3784959 (SMAL23_35260) | 1596104-1596889 (SMAL23_15250-SMAL23_15260) | 2414051-2414247 (SMAL23_22860) |
| 8-STEN-2 DNA |  |  | 2418281-2418468 (SMAL8_23070) |
| P28 | 4369745-4370503 (QVH29_20420) | 1978915-1979700 (QVH29_09400-QVH29_09405) | 2981009-2981215 (QVH29_14025) |
| 503 | 4272509-4273267 | 1982670-1983455 |  |
| HAMBI_2659 | 4401498-4402256 (U0039_20665) | 2189931-2190716 (U0039_10600-U0039_10605) | 2976320-2976516 (U0039_14125) |
| CGMCC1.1788 | 4399624-4400382 | 2189944-2190729 | 2975715-2975911 |
| SM866 | 4521459-4522217 (DUW70_21530) | 3099977-3100762 (DUW70_14680-DUW70_14685) |  |
| 1945B | C4063269-4064027 (ABK042_18975) | 1617016-1617801 (ABK042_07675-ABK042_07680) | C773972-774178 (ABK042_03820) |
| RY12679 | 4341857-4342609 (ACS00Y_20200) | 2049231-2050016 (ACS00Y_09820-ACS00Y_09825) |  |
| RY12681 | 4198448-4199212 (ACS00U_19530) | 1928613-1929398 (ACS00U_09190-ACS00U_09195) | 2726876-2727082 (ACS00U_12730) |
| RY12687 | 4295681-4296445 (ACS00W_20240) | 1979042-1979827 (ACS00W_09545-ACS00W_09550) | 2783222-2783428 (ACS00W_13115) |
| RY12688 |  |  | 2483813-2484000 (ACS00T_11685) |
| SM1053 | 3610981-3611745 | 1459804-1460589 | 2274857-2275063 |
| SM1021 | 4033695-4034459 | 1882599-188338 | 2697652-269785 |
| SM922 | 497587-498351 | 2648738-2649523 | 1834266-1834472 |
| SM911 | 497586-498350 | 2648657-2649442 | 1834185-1834391 |
| c31 | 4000768-4001526 (KNV95_18580) | 1872529-1873314 (KNV95_08870-KNV95_08875) | 2601919-2602125 (KNV95_12155) |
| S62 | 4301494-4302252 (ABFB11_19780) | C2913466-2914251 (ABFB11_13425-ABFB11_13430) | 2068310-2068506 (ABFB11_09600) |
| C52 | 4299863-4300621 (AASM09_19760) | C2912311-2913096 (AASM09_13415-AASM09_13420) | 2067586-2067782 (AASM09_09595) |
| 9605 | 718471-719229 (ABK044_03155) | C3980545-3981330 (ABK044_18085-ABK044_18090) | C3300749-3300945 (ABK044_15065) |
| STEN00241 |  |  |  |
| PSKL1 |  |  |  |
| ZSL2493 |  |  |  |
| R-24 |  | C3030904-3031689 (ACN9MD_13700-ACN9MD_13705) | C2239692-2239870 (ACN9MD_10225) |
| GIMC6004:STM-35P13 |  |  |  |

Bhaumik et al. Supplementary Table 3
| Strain | putative TadE family transmembrane pilus related protein | putative fimbriae assembly protein family | putative type IV pilus protein |
| --- | --- | --- | --- |
| AA1 |  |  |  |
| AB550 | 2733037-2733513 (B7H26_12830-12835) |  |  |
| CF13 | C2022956-2023432 (HKK60_09245-09250) | 2019237-2022932 (HKK60_09225-09240) |  |
| Col1 | 2522587-2523063 | 2523087-2526785 |  |
| CPBW01 | 2545588-2546064 (GS396_11600-11605) | 2546088-2549786 (GS396_11610-11625) |  |
| CSM2 |  | 2760341-2764036 (SmaCSM2_12725-12740) |  |
| D457 | 2797292-2797768 (SMD_2506-2507) |  |  |
| DHHJ |  | 2644831-2648529 (JIL50_12235-12250) |  |
| FDAARGOS_1044 | 2683555-2684031 (I6I01_12710-12715) | 2684055-2687750 (I6I01_12720-12735) | C1231038-1232825 (I6I01_06015) |
| FDAARGOS_325 | 263980-264456 (CEQ03_01275-01280) | C260261_263956 (CEQ03_01255-01270) |  |
| FDAARGOS_507 | 2305457-2305933 (EGY09_10915-10920) | 2305957-2309652 (EGY09_10925-10940) | 2504857-2506644 (EGY09_11920) |
| FDAARGOS_649 | 4030424-4030900 (FOB57_18675-18680) | 4030924-4034619 (FOB57_18685-18700) |  |
| FDAARGOS_92 | 1838700-1839176 (AL480_09025-09030) | 1839200-1842895 (AL480_09035-09050) | C421249-423036 (AL480_02145) |
| ICU331 | C2154530-2155006 (FEO84_10390-10395) | C2150811-2154506 (FEO84_10370-10385) | C1963092-1964890 (FEO84-09460) |
| ISMMS2 | 2615916-2616392 (YH67_11910-11915) | 2616416-2620114 (YH67_11920-11935) |  |
| ISMMS2R | 2615916-2616392 (YH68_11910-11915) | 2616416-2620114 (YH68_11920-11935) |  |
| ISMMS3 |  |  |  |
| JV3 |  | 2619680-2623375 (BurJV3_2377-2380) |  |
| K279a | 2910775-2911251 (Smlt2868-2869) | 2911275-2914970 (Smlt2870-2873) | 3114059-3115846 (Smlt3071) |
| KMM 349 | 2653190-2653630 (KMM349_23730) | 2653654-2657352 (KMM349_23740-23770) |  |
| MER1 |  |  |  |
| NCTC10257 |  |  |  |
| NCTC10258 |  |  |  |
| NCTC10259 |  |  |  |
| NCTC10498 (SAMN14206046) | 2268519-2268995 | 2269019-2272714 |  |
| NCTC13014 |  |  |  |
| NEB515 | C2342565-2343041 (HGN30_10640-10645) | 2338847-2342541 (HGN30_10620-10635) |  |
| O1 | 1842249-1842725 | 1842749-1846437 |  |
| OUC_Est10 | 2658944-2659420 (A7326_12110-12115) | 2659444-2663139 (A7326_12120-12135) |  |
| PEG-141 |  | 2968943-2972641 (FEO88_13990-14005) |  |
| PEG-173 | 2782815-2783291 (FEO89_12945-12950) | 2783315-2787007 (FEO89_12955-12970) |  |
| PEG-305 | 2623014-2623490 (FEO90_12065-12070) | 2623516-2627215 (FEO90_12075-12090) |  |
| PEG-390 | 2681434-2681910 (FEO91_12575-12580) | 2681934-2685632 (FEO91_12585-12600) |  |
| PEG-42 | 2827508-2827984 (FEO92_13295-13300) | 2828008-2831702 (FEO92_13305-13320) | 3034256-3036043 (FEO92_14330) |
| PEG-68 | 2645408-2645884 (FEO93_12450-12455) | 2645910-2649605 (FEO93_12460-12475) |  |
| R551-3 | 2608623-2609063 (Smal_2319) | 2609087-2612785 (Smal_2320-2323) |  |
| SJTH1 | 2819684-2820160 (C6Y55_13100-13105) | 2820184-2823879 (C6Y55_13110-13125) | 3005947-3007734 (C6Y55_14045) |
| SJTL3 | 2804721-2805197 (DM611_13250-13255) | 2805221-2808922 (DM611_13260-13275) |  |
| SKK55 |  |  |  |
| SM 866 |  | C2244685-2248383 (DUW70_10755-10770) |  |
| sm454 | 2721635-2722111 (FEO85_12830-12835) | 2722135-2725831 (FEO85_12840-12855) |  |
| Sm53 | 2784072-2784548 (FEO86_13210-13215) | 2784572-2788267 (FEO86_13220-13235) |  |

Bhaumik et al. Supplementary Table 3
| Strain | putative TadE family transmembrane pilus related protein | putative fimbriae assembly protein family | putative type IV pilus protein |
| --- | --- | --- | --- |
| sm-RA9 |  |  | 3087495-3089282 (FEO95_14300) |
| SoD9b | 2575610-2576085 (G4G30_11865-11870) |  |  |
| T50-20 |  | 2789686-2793384 |  |
| U5 |  |  |  |
| W18 | 2771650-2772126 | 2772150-2775844 |  |
| WP1-W18-CRE-01 | 2838339-2838815 (WP1W18C01_25840-25850) | 2838839-2842534 (WP1W18C01_25860-25890) | C1511786-1513573 (WP1W18C01_14080) |
| X28 | 569484-569960 | 569984-573680 |  |
| XL133 | 2747525-2748001 (LAD79_12745-12750) | 2748025-2751720 (LAD79_12755-12770) |  |
| JZL8 | 2660012-2660488 | 2660512-2664210 |  |
| 1800 |  |  |  |
| 454 | 2720980-2721456 (SMAL454_25430-25440) | 2721480-2725176 (SMAL454_25450-25480) |  |
| SCAID WND1-2022 (370) | 2870262-2870738 (NW343_13335-13340) | 2870762-2874457 (NW343_13345-13360) |  |
| a7 | 2610087-2610563 | 2610587-2614282 |  |
| ACYCe.8N | 2603828-2604304 (K7573_12010-12015) | 2604328-2608023 (K7573_12020-12035) |  |
| 2013-SM15 | 4084584-4085060 (LQ329_18715-18720) | 4080865-4084560 (LQ329_18695-18710) | 3875626-3877424 |
| ACYCa.6E | 2596191-2596667 (K7574_12060-12065) | 2596691-2600386 (K7574_12070-12085) |  |
| CYZ | 1855394-1855870 | 1851675-1855370 |  |
| 142 | 2818124-2818600 (NDK23_13195-13200) | 2818624-2822319 (NDK23_13205-13220) | 3010145-3011932 (NDK23_14130) |
| PSKL2 | 2620639-2621115 (L6173_12035-12040) | 2621139-2624833 (L6173_12045-12060) |  |
| 11066 | 2631443-2631919 (RBI20_12270-12275) | 2631943-2635638 (RBI20_12280-12295) |  |
| HT2 | 2664407-2664883 (PWE35_12460-PWE35_12465) | 2664907-2668602 (PWE35_12470-12485) |  |
| 2013-SM24 | 2387341-2387817 (LQ331_11195-LQ331_11200) | 2387841-2391536 (LQ331_11205-11220) |  |
| FZD2 | 2700429-2700905 (K1516-00005-K1516_00010) | 2700929-2704630 (K1516_12505-12520) |  |
| ACYCd.9D | 2624024-2624500 (K7563_12140-12145) |  |  |
| 2013-SM4 | 2731296-2731772 (LQ335_12585-12590) |  |  |
| ACYCc.3B | 2527517-2527993 (K7564_11975-11980) | 2528019-2531714 (K7564_11985-12000) |  |
| 2013-SM12 | 41007-41459 (LQ332_00175) |  |  |
| WGB211 | 2835800-2836276 | 2836300-2839995 | 3028370-3030157 |
| ACYCb.6H | 2487951-2488427 (K7565_11465-11470) | 2488451-2492149 (K7565_11475-11490) |  |
| ZT1 | C2190603-2191079 (J3U96_09930-09935) | C2186881-2190579 (J3U96_09910-09925) |  |
| GYH |  |  | C417640-419427 (LZ605_02200) |
| ACYCb.1K | 2526771-2527247 (K7567_11680-11685) | 2527271-2530969 (K7567_11690-11705) |  |
| ACYCb.10K | C2745554-2746006 (K7569_12930) | C2741832-2745530 (K7569_12910-12925) |  |
| ACYCa.2H | 2641183-2641659 (K7568_12210-12215) | 2641683-2645378 (K7568_12220-12235) |  |
| 2013-SM13 | 3508684-3509136 (LQ330_16250) | 3509162-3512861 (LQ330_16255-16270) |  |
| CW002SM | C2053946-2054422 (N0074_09395-09400) | C2050229-2053922 (N0074_09370-09390) |  |
| ACYCa.1J | 2788685-2789161 (K7566_13000-13005) | 2789185-2792880 (K7566_13010-13025) |  |
| HW002Y | 2234053-2234529 (NDR96_10365-10370) | 2234553-2238251 (NDR96_10375-10390) |  |
| SI1 |  |  | 3125273-3127060 (RS400_14645) |
| 24(2023) |  |  |  |
| STK21_22 | 2857691-2858167 (ACSYTB_002661-ACSYTB_002662) | 2858191-2861886 (ACSYTB_002663-ACSYTB_002666) |  |

Bhaumik et al. Supplementary Table 3
| Strain | putative TadE family transmembrane pilus related protein | putative fimbriae assembly protein family | putative type IV pilus protein |
| --- | --- | --- | --- |
| STK4_22 | 2761737-2762213 (ACSYS1_002618-ACSYS1_002619) | 2762237-2765932 (ACSYS1_002620-ACSYS1_002623) | 2946310-2948097 (ACSYS1_002800) |
| STK9_22 | 2773917-2774393 (ACSYS3_002606-ACSYS3_002607) | 2774417-2778112 (ACSYS3_002608-ACSYS3_002611) | 2958490-2960277 (ACSYS3_002788) |
| STK11_22 | 2761855-2762331 (ACSYS4_002615-ACSYS4_002616) | 2762355-2766050 (ACSYS4_002617-ACSYS4_002620) | 2946428-2948215 (ACSYS4_002797) |
| STK12_22 | 2761748-2762224 (ACSYS5_002619-ACSYS5_002620) | 2762248-2765943 (ACSYS5_002621-ACSYS5_002624) | 2946320-2948107 (ACSYS5_002801) |
| LC734 | 2845628-2846104 (ACT4MX_13335-ACT4MX_13340) | 2846128-2849823 (ACT4MX_13345-ACT4MX_13360) |  |
| 15-STEN-3 DNA | 2867447-2867923 (SMAL15_26850-SMAL15_26860) | 2867947-2871642 (SMAL15_26870-SMAL15_26900) |  |
| 25-STEN-6 DNA | C1837555-1838031 (SMAL25_16510-SMAL25_16520) | C1833833-1837531 (SMAL25_16470-SMAL25_16500) |  |
| 23-STEN-5 DNA | 2414883-2415359 (SMAL23_22870-SMAL23_22880) | 2415383-2419078 (SMAL23_22890-SMAL23_22920) |  |
| 8-STEN-2 DNA |  |  |  |
| P28 | 2981846-2982322 (QVH29_14030-QVH29_14035) | 2982346-2986041 (QVH29_14040-QVH29_14055) | C1417213-1418991 (QVH29_06735) |
| 503 |  |  | 2952967-2954754 |
| HAMBI_2659 | 2977152-2977628 (U0039_14130-U0039_14135) | 2977652-2981347 (U0039_14140-U0039_14155) | C1524635-1526422 (U0039_07410) |
| CGMCC1.1788 | 2976547-2977023 | 2977047-2980732 | 1523667-1525454 |
| SM866 |  | C2244685-2248383 (DUW70_10755-DUW70_10770) |  |
| 1945B | C772865-773341 (ABK042_03810-ABK042_03815) | C769146-772841 (ABK042_03790-ABK042_03805) | C584491-586278 (ABK042_02895) |
| RY12679 | 2929460-2929460 (ACS00Y_13795-ACS00Y_13800) | 2929484-2933179 (ACS00Y_13805-ACS00Y_13820) |  |
| RY12681 | 2727713-2728189 (ACS00U_12735-ACS00U_12740) | 2728213-2731908 (ACS00U_12745-ACS00U_12760) | 2912281-2914068 (ACS00U_13645) |
| RY12687 | 2784059-2784535 (ACS00W_13120-ACS00W_13125) | 2784559-2788254 (ACS00W_13130-ACS00W_13145) | 2968624-2970411 (ACS00W_14030) |
| RY12688 |  |  |  |
| SM1053 | 2275694-2276170 | 2276194-2279889 |  |
| SM1021 | 2698489-2698965 | 2698989-2702684 |  |
| SM922 | 1833159-1833635 | 1829440-1833135 |  |
| SM911 | 1833078-1833554 | 1829359-1833054 |  |
| c31 | 2602756-2603232 (KNV95_12160-KNV95_12165) | 2603256-2606951 (KNV95_12170-KNV95_12185) |  |
| S62 | C2067198-2067674 (ABFB11_09590-ABFB11_09595) | C2063478-2067174 (ABFB11_09570-ABFB11_09585) |  |
| C52 | C2066474-2066950 (AASM09_09585-AASM09_09590) | C2062754-2066450 (AASM09_09565-AASM09_09580) |  |
| 9605 | C3299637-3300113 (ABK044_15055-ABK044_15060) | C3295917-3299613 (ABK044_15035-ABK044_15050) |  |
| STEN00241 |  |  | 2888081-2889868 (RWT08_13190) |
| PSKL1 |  |  |  |
| ZSL2493 |  |  |  |
| R-24 | C2238590-2239030 (ACN9MD_10215) |  |  |
| GIMC6004:STM-35P13 |  |  |  |

Bhaumik et al. Supplementary Table 3
| Strain | putative type II/IV secretion system pilus related ATP-binding protein | putative type II secretion/pilus assembly |
| --- | --- | --- |
| AA1 |  |  |
| AB550 |  | 2738587-2739527 (B7H26_12860) |
| CF13 | C2017903-2019252 (HKK60_09225-09230) | 2016944-2017884 (HKK60_09220) |
| Col1 | 2526770-2528119 |  |
| CPBW01 | 2549771-2551120 (GS396_11620-11625) | 2551139-2552079 (GS396_11630) |
| CSM2 | 2764021-2765370 (SmaCSM2_12735-12740) | 2765389-2766329 (SmaCSM2_12745) |
| D457 |  |  |
| DHHJ | 2648514-2649863 (JL50_12245-12250) | 2649882-2650822 (JL50_12255) |
| FDAARGOS_1044 | 2687735-2689084 (I6I01_12730-12735) | 2689103-2690043 (I6I01_12740) |
| FDAARGOS_325 | C258927-260276 (CEQ03_01255-01260) | 257968-258908 (CEQ03_01250) |
| FDAARGOS_507 | 2309637-2310986 (EGY09_10935-10940) | 2311005-2311945 (EGY09_10945) |
| FDAARGOS_649 | 4034604-4035953 (FOB57_18695-18700) | 4035972-4036912 (FOB57_18705) |
| FDAARGOS_92 | 1842880-1844229 (AL480_09045-09050) | 1844248-1845188 (AL480_09055) |
| ICU331 | C2149477-2150826 (FEO84_10370-10375) | 2148518-2149458 (FEO84_10365) |
| ISMMS2 | 2620099-2621448 (YH67_11930-11935) | 2621467-2622407 (YH67_11940) |
| ISMMS2R | 2620099-2621448 (YH68_11930-11935) | 2621467-2622407 (YH68_11940) |
| ISMMS3 | 2773318-2774662 (VN11_12700-12705) |  |
| JV3 | 2623360-2624709 (BurJV3_2379-2380) | 2624728-2625668 (BurJV3_2381) |
| K279a | 2914955-2916304 (Smlt2872-2873) | 2916322-2917263 (Smlt2874) |
| KMM 349 | 2657337-2658686 (KMM349_23760-23770) | 2658705-2659645 (KMM349_23780) |
| MER1 |  |  |
| NCTC10257 |  |  |
| NCTC10258 |  |  |
| NCTC10259 |  |  |
| NCTC10498 (SAMN14206046) | 2272699-2274048 | 2274067-2275007 |
| NCTC13014 |  |  |
| NEB515 | C2337513-2338862 (HGN30_10620-10625) | C2336554-2337494 (HGN30_10615) |
| O1 | 1846424-1847770 | 1847789-1848727 |
| OUC_Est10 | 2663124-2664473 (A7326_12130-12135) | 2664492-2665432 (A7326_12140) |
| PEG-141 | 2972626-2973975 (FEO88_14000-14005) | 2973994-2974934 (FEO88_14010) |
| PEG-173 | 2786992-2788341 (FEO89_12965-12970) | 2788360-2789300 (FEO89_12975) |
| PEG-305 | 2627200-2628549 (FEO90_12085-12090) | 2628568-2629508 (FEO90_12095) |
| PEG-390 | 2685617-2686966 (FEO91_12595-12600) |  |

Bhaumik et al. Supplementary Table 3
| Strain | putative type II/IV secretion system pilus related ATP-binding protein | putative type II secretion/pilus assembly |
| --- | --- | --- |
| PEG-42 | 2831687-2833036 (FEO92_13315-13320) | 2833055-2833995 (FEO92_13325) |
| PEG-68 | 2649590-2650939 (FEO93_12470-12475) | 2650958-2651898 (FEO93_12480) |
| R551-3 | 2612770-2614119 (SmaI_2322-2323) | 2614138-2615078 (SmaI_2324) |
| SJTH1 | 2823864-2825213 (C6Y55_13120-13125) | 2825232-2826172 (C6Y55_13130) |
| SJTL3 | 2808907-2810256 (DM611_13270-13275) | 2810275-2811215 (DM611_13280) |
| SKK55 |  |  |
| SM 866 | C2243351-2244700 (DUW70_10755-10760) | C2242392-2243332 (DUW70_10750) |
| sm454 | 2725816-2727165 (FEO85_12850-12855) | 2727184-2728124 (FEO85_12860) |
| Sm53 | 2788252-2789601 (FEO86_13230-13235) | 2789620-2790560 (FEO86_13240) |
| sm-RA9 |  |  |
| SoD9b |  |  |
| T50-20 | 2793369-2794718 | 2794737-2795677 |
| U5 |  |  |
| W18 | 2775829-2777178 | 2777197-2778137 |
| WP1-W18-CRE-01 | 2842519-2843868 (WP1W18C01_25880-25890) | 2843886-2844827 (WP1W18C01_25900) |
| X28 | 573665-575014 | 575033-575973 |
| XL133 | 2751705-2753054 (LAD79_12765-12770) | 2753073-2754013 (LAD79_12775) |
| JZL8 | 2664195-2665544 | 2665563-2666503 |
| 1800 |  |  |
| 454 | 2725161-2726510 (SMAL454_25470-25480) | 2726529-2727469 (SMAL454_25490) |
| SCAID WND1-2022 (370) | 2874442-2875791 (NW343_13355-13360) | 2875810-2876750 (NW343_13365) |
| a7 | 2614267-2615616 | 2615635-2616575 |
| ACYCe.8N | 2608008-2609357 (K7573_12030-12035) | 2609376-2610316 (K7573_12040) |
| 2013-SM15 | C4079531-4080880 (LQ329_18695-18700) | 4078572-4079512 (LQ329_18690) |
| ACYCa.6E | 2600371-2601720 (K7574_12080-12085) | 2601739-2602679 (K7574_12090) |
| CYZ | 1850341-1851690 | 1849382-1850322 |
| 142 | 2822304-2823653 (NDK23_13215-13220) | 2823672-2824612 (NDK23_13225) |
| PSKL2 | 2624818-2626167 (L6173_12055-12060) | 2626186-2627126 (L6173_12065) |
| 11066 | 2635623-2636972 (RBI20_12290-12295) | 2636991-2637931 (RBI20_12300) |
| HT2 | 2668587-2669936 (PWE35_12480-12485) | 2669955-2670895 (PWE35_12490) |
| 2013-SM24 | 2391521-2392870 (LQ331_11215-11220) | 2392889-2393829 (LQ331_11225) |
| FZD2 | 2704615-2705964 (K1516_12515-12520) | 2705983-2706923 (K1516_12525) |
| ACYCd.9D | 2628210-2629556 (K7563_12160-12165) |  |

Bhaumik et al. Supplementary Table 3
| Strain | putative type II/IV secretion system pilus related ATP-binding protein | putative type II secretion/pilus assembly |
| --- | --- | --- |
| 2013-SM4 | 2735482-2736828 (LQ335_12605-12610) |  |
| ACYCc.3B | 2531699-2533048 (K7564_11995-12000) | 2533067-2534007 (K7564_12005) |
| 2013-SM12 |  | 46537-47477 (LQ332_00200) |
| WGB211 | 2839980-2841329 | 2841348-2842288 |
| ACYCb.6H | 2492134-2493483 (K7565_11485-11490) | 2493502-2494442 (K7565_11495) |
| ZT1 | 2185547-2186896 (J3U96_09910-09915) |  |
| GYH |  |  |
| ACYCb.1K | 2530954-2532303 (K7567_11700-11705) |  |
| ACYCb.10K | C2740498-2741847 (K7569_12910-12915) | C2739539-2740467 (K7569_12905) |
| ACYCa.2H | 2645363-2646712 (K7568_12230-12235) | 2646731-2647671 (K7568_12240) |
| 2013-SM13 |  | 3514214-3515154 (LQ330_16275) |
| CW002SM | 2048895-2050244 (N0074_09370-09375) | C2047936-2048876 (N0074_09365) |
| ACYCa.1J | 2792865-2794214 (K7566_13020-13025) | 2794233-2795173 (K7566_13030) |
| HW002Y | 2238236-2239585 (NDR96_10385-10390) | 2239604-2240544 (NDR96_10395) |
| SI1 |  |  |
| 24(2023) |  |  |
| STK21_22 | 2861871-2863220 (ACSYTB_002665-ACSYTB_002666) | 2863239-2864179 (ACSYTB_002667) |
| STK4_22 | 2765917-2767266 (ACSYS1_002622-ACSYS1_002623) | 2767285-2768225 (ACSYS1_002624) |
| STK9_22 | 2778097-2779446 (ACSYS3_002610-ACSYS3_002611) | 2779465-2780405 (ACSYS3_002612) |
| STK11_22 | 2766035-2767384 (ACSYS4_002619-ACSYS4_002620) | 2767403-2768343 (ACSYS4_002621) |
| STK12_22 | 2765928-2767277 (ACSYS5_002623-ACSYS5_002624) | 2767296-2768236 (ACSYS5_002625) |
| LC734 |  | 2851176-2852116 (ACT4MX_13365) |
| 15-STEN-3 DNA | 2871627-2872973 (SMAL15_26890-SMAL15_26900) | 2872992-2873932 (SMAL15_26910) |
| 25-STEN-6 DNA | C1832499-1833848 (SMAL25_16470-SMAL25_16480) |  |
| 23-STEN-5 DNA | 2419063-2420409 (SMAL23_22910-SMAL23_22920) | 2420428-2421368 (SMAL23_22930) |
| 8-STEN-2 DNA |  |  |
| P28 | 2986026-2987375 (QVH29_14050-QVH29_14055) | 2987394-2988334 (QVH29_14060) |
| 503 |  |  |
| HAMBI_2659 | 2981332-2982681 (U0039_14150-U0039_14155) | 2982700-2983640 (U0039_14160) |
| CGMCC1.1788 | 2980717-2982066 | 2982085-2983025 |
| SM866 |  |  |
| 1945B | C767812-769161 (ABK042_03790-ABK042_03795) | C766853-767794 (ABK042_03785) |
| RY12679 | 2933164-2934513 (ACS00Y_13815-ACS00Y_13820) | 2934532-2935472 (ACS00Y_13825) |

**Bhaumik et al. Supplementary Table 3**
| **Strain** | **putative type II/IV secretion system pilus related ATP-binding protein** | **putative type II secretion/pilus assembly** |
| --- | --- | --- |
| **RY12681** | 2731893-2733242 (ACS0OU_12755-ACS0OU_12760) | 2733261-2734201 (ACS0OU_12765) |
| **RY12687** | 2788239-2789588 (ACS0OW_13140-ACS0OW_13145) | 2789607-2790547 (ACS0OW_13150) |
| **RY12688** |  |  |
| **SM1053** | 2279874-2281223 | 2281242-228218 |
| **SM1021** | 2702669-2704018 | 2704037-2704977 |
| **SM922** | 1828106-1829455 | 1827147-1828087 |
| **SM911** | 1828025-1829374 | 1827066-1828006 |
| **c31** | 2606936-2608285 (KNV95_12180-KNV95_12185) | 2608304-2609244 (KNV95_12190) |
| **S62** | C2062144-2063493 (ABFB11_09570-ABFB11_09575) | C2061185-2062125 (ABFB11_09565) |
| **C52** | C2061420-2062769 (AASM09_09565-AASM09_09570) | C2060461-2061401 (AASM09_09560) |
| **9605** | C3294583-3295932 (ABK044_15035-ABK044_15040) | C3293624-3294564 (ABK044_15030) |
| **STEN00241** |  |  |
| **PSKL1** |  |  |
| **ZSL2493** |  |  |
| **R-24** | C2233534-2234883 (ACN9MD_10195-ACN9MD_10200) | C2232575-2233515 (ACN9MD_10190) |
| **GIMC6004:STM-35P13** |  |  |

